# Quantifying the Influence of Mutation Detection on Tumour Subclonal Reconstruction

**DOI:** 10.1101/418780

**Authors:** Lydia Y. Liu, Vinayak Bhandari, Adriana Salcedo, Shadrielle M. G. Espiritu, Quaid D. Morris, Thomas Kislinger, Paul C. Boutros

## Abstract

Whole-genome sequencing can be used to estimate subclonal populations in tumours and this intra-tumoural heterogeneity is linked to clinical outcomes. Many algorithms have been developed for subclonal reconstruction, but their variabilities and consistencies are largely unknown. We evaluated sixteen pipelines for reconstructing the evolutionary histories of 293 localized prostate cancers from single samples, and eighteen pipelines for the reconstruction of 10 tumours with multi-region sampling. We show that predictions of subclonal architecture and timing of somatic mutations vary extensively across pipelines. Pipelines show consistent types of biases, with those incorporating SomaticSniper and Battenberg preferentially predicting homogenous cancer cell populations and those using MuTect tending to predict multiple populations of cancer cells. Subclonal reconstructions using multi-region sampling confirm that single-sample reconstructions systematically underestimate intra-tumoural heterogeneity, predicting on average fewer than half of the cancer cell populations identified by multi-region sequencing. Overall, these biases suggest caution in interpreting specific architectures and subclonal variants.

## Introduction

Understanding tumour heterogeneity and subclonal architecture is important for the elucidation of the mutational and evolutionary processes underlying tumorigenesis and treatment resistance^1–4^. Many studies of tumour heterogeneity have focused on small patient cohorts with multi-region sequencing^5–11^. This study design allows the reconstruction of sample trees that illustrate the relationships between multiple primary and metastatic lesions using shared and private mutations^6,11^. Despite their small sample sizes, these studies have provided remarkable insight, demonstrating multiple subclones within a single tumour, clonal relationships between primary and metastatic tumours and evidence for multiple primary tumours within a single patient. Many studies have further delved into intra-tumoural heterogeneity and constructed clone trees that demonstrate the phylogenetic relationship between cancer cell populations that are shared or unique between lesions^5,7,9,12^. The latter analyses not only provide insight to the convergent and branching evolution of cancer, but also characterize cancer cell migration and highlight the subclonal complexity within individual lesions.

Some studies have applied these techniques to large cohorts of single-region tumour whole genomes. For example, we reconstructed the subclonal architectures of 293 localized prostate cancers using whole-genome sequencing (WGS) of a single region of the index lesion^13^. The larger sample sizes of single-region studies allow the identification of mutational events that are biased to occur at specific times during tumour development. Single-region subclonal reconstruction studies have also suggested that patients with less subclonal diversity (*e.g.* with only a single detectable population of cancer cells; termed *monoclonal*) tend to have superior clinical outcomes compared to those with more subclonal diversity (*e.g.* those with highly *polyclonal* tumours)^13^.

A variety of algorithms have been developed to reconstruct the subclonal architecture of cancers from single-region or multi-region bulk DNA sequencing data^14–21^. These algorithms broadly attempt to infer cancer cell populations based on cancer cell fractions (the fraction of cancer cells in which each variant is present) of somatic single nucleotide variants (SNVs) and/or somatic copy number aberrations (CNAs). Several employ Bayesian models to cluster mutations, and estimate the number and prevalence of cancer cell populations^15–17,20,22^. Some algorithms are further able to infer phylogenetic clone trees, thus resolving the evolutionary relationship between mutation clusters^15,21^. However, there has not been a systematic comparison of the features and consistencies of their reconstructions on a large dataset. It is thus unclear to what extent these pipelines agree on large cohorts of real data, whether specific pipelines are biased towards certain types of reconstructions, and to what degree reconstruction results are influenced by the somatic mutation inputs. It is further unclear to what extent single-sample reconstructions differ from multi-region reconstructions, raising questions on the magnitude of underestimation present in large-cohort studies.

To address these gaps in the field, we evaluated pipelines consisting of twenty-two different combinations of well-established and independent SNV detection tools, subclonal CNA detection tools and subclonal reconstruction algorithms. Sixteen pipelines were applied to a set of 293 high-depth tumour-normal pairs^13,23^ and eighteen were applied to 10 tumours with multi-region sequencing^8,24^. Our analyses reveal consistent biases and extensive differences across subclonal reconstruction pipelines in the predictions of subclonal architecture, identification and timing of variants and influence on downstream analyses. We also quantify the extent that single-region reconstructions underestimate intra-tumour heterogeneity as compared to reconstructions based on multiple regions of the tumour. Together, these findings generate guidance for the community and provide a resource for improving existing methods and benchmarking new ones.

## Results

### Overview and Summary of Pipeline Runs

We reconstructed the subclonal architectures of 293 primary localized prostate tumours using sixteen pipelines (**Figure 1**, **Supplementary Data 1-20**). Each patient had WGS of a single region taken from the index lesion (**Methods**) that was macro-dissected to > 70% tumour cellularity (mean coverage ± standard deviation [SD]: 63.9 ± 16.7) and WGS of matched blood reference tissue (mean coverage ± SD: 41.2 ± 9.0), as reported previously^13^. To investigate the influence of variant detection on subclonal reconstruction, we detected CNAs using Battenberg and TITAN^7,25^ and SNVs using SomaticSniper and MuTect^26,27^. We then used the CNAs and SNVs detected by these tools in factorial combinations as inputs for four widely-used subclonal reconstruction algorithms: PhyloWGS^15^, DPClust^16^, PyClone^17^ and SciClone^20^. Each subclonal reconstruction pipeline was thus composed of three algorithms: a SNV detection tool, a subclonal CNA detection tool and a subclonal reconstruction algorithm. Thus “PhyloWGS-comprising pipelines” refers to all pipelines that use PhyloWGS as the subclonal reconstruction algorithm, in combination with any SNV and CNA detection tool. All subclonal reconstruction solutions were subjected to the same post-processing heuristics to minimize bias (**Methods**). We further quantified the variability that arises in subclonal reconstruction from spatially sampling the same tumour, focusing on ten tumours with multi-region WGS (2-4 regions per tumour, total of 30 regions)^8,24^. Multi-region WGS samples were further assessed using FACETS^28^ for subclonal CNA detection, and subclonal reconstruction was performed both with all regions together and with each region individually using PhyloWGS, PyClone and SciClone.

**Figure 1.**
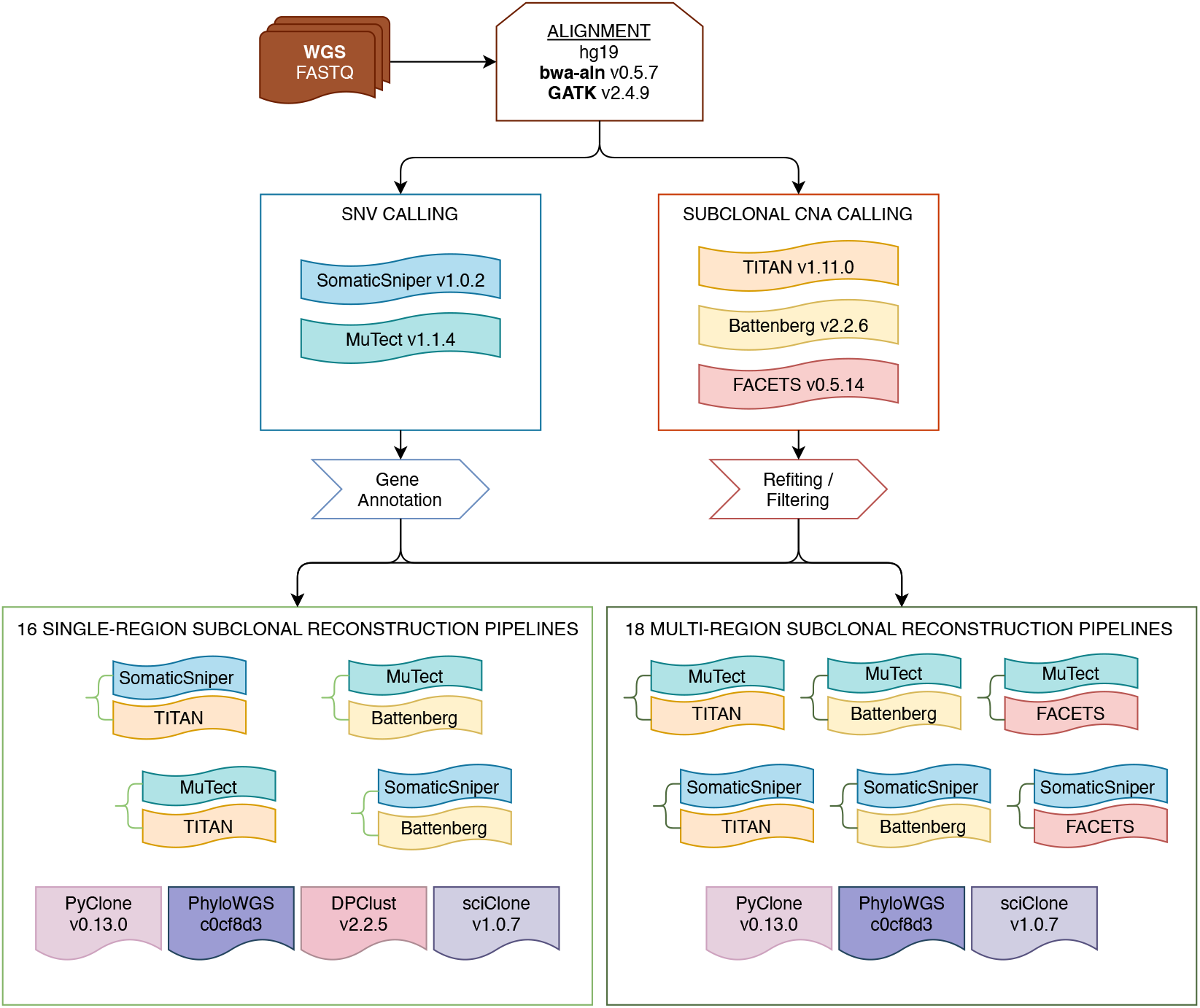
Reconstruction Workflow and Experimental Design. Raw sequencing data from the tumour and normal samples were aligned against the hg19 build of the human genome using bwa-aln and GATK. Somatic SNVs were detected using SomaticSniper and MuTect and annotated for function. Somatic CNAs were detected using TITAN, Battenberg and FACETS and filtered. All single-region tumour samples had their subclonal architectures reconstructed using sixteen pipelines combining one of SomaticSniper and MuTect, one of Battenberg and TITAN, and one of PyClone, PhyloWGS, DPClust and SciClone. For tumours with samples from multiple regions, reconstructions of subclonal architectures were performed by considering each individual region alone and by considering samples from all regions together using eighteen pipelines. SNV, single nucleotide variant; CNA, copy number aberration; WGS, whole-genome sequencing.

Across all samples and pipelines, we attempted to execute 5408 subclonal reconstructions. Of these, 4447 (82.2%) successfully completed their execution (**Supplementary Table 1**). Among pipelines for the single-region subclonal reconstruction of 293 tumours, those using DPClust achieved the lowest failure rates (mean ± SD: 1.4% ± 1.5%), followed by those using PhyloWGS (2.2% ± 1.3%), PyClone (16.3% ± 9.8%) and SciClone (41.2% ± 22.4%; **Supplementary Figure 1A**). The primary reasons of failure for pipelines using DPClust and PhyloWGS were excessive memory requirements (> 250 GB RAM) or run-time (> 3 months). Lack of input SNVs was the largest failure reason for pipelines using PyClone and SciClone, as PyClone exclusively leverage SNVs from clonal CNA regions and SciClone utilizes SNVs in copy number neutral regions. Since we used CNA detection tools that identified subclonal variation, in some cases insufficient clonal CNA regions were available. Post-processing heuristics also contributed to reconstruction failures across pipelines (**Methods**).

Multi-region reconstructions with pipelines using PhyloWGS had the lowest failure rates on the 10 tumours evaluated (mean failure rate ± SD: 5.0% ± 5.5%), followed by PyClone (45.0% ± 26.6%) and SciClone (93.3% ± 10.7%; **Supplementary Figure 1B**). Reasons of failure for pipelines using PhyloWGS include lack of shared CNAs between samples from the same tumour and prediction of poly-tumour architectures (*i.e.*, multiple independent primary tumours; **Methods**). PyClone leverages SNVs in clonal CNA regions that are shared between all samples from the same tumour for multi-region reconstructions and had higher failure rates. Due to similar requirements for SciClone that all SNVs be in copy number neutral regions and shared between all samples from the tumour, multi-region reconstructions using SciClone only succeeded in four cases overall and were excluded from further multi-region reconstruction analyses.

### Consistency of Subclonal Reconstruction from Single Samples

To evaluate subclonal reconstruction solutions for 293 single-region tumours, we first compared tumour cellularity (sometimes called “tumour purity”) estimates across subclonal reconstruction pipelines. Cellularity estimates from CNA detection tools are inputs to PhyloWGS, PyClone and DPClust, and as expected predicted cellularity from pipelines using these algorithms correlated with those from the CNA detection tool used (TITAN: 0.212–0.623, Battenberg: 0.588–0.876, Spearman’s *ρ*; **Figure 2A-B**). By contrast, SciClone predicts sample cellularity using orthogonal evidence (VAF of SNVs in copy number neutral regions). SciClone-estimated cellularity in pipelines using SomaticSniper correlated better with estimates from CNA detection tools (SomaticSniper-TITAN-SciClone *vs.* TITAN: 0.363, SomaticSniper-Battenberg-SciClone *vs.* Battenberg: 0.670, Spearman’s *ρ*) than did pipelines using MuTect (MuTect-TITAN-SciClone *vs.* TITAN: 0.035, MuTect-Battenberg-SciClone *vs.* Battenberg: 0.348, Spearman’s *ρ*). This suggests that the VAFs of SNVs detected by MuTect have biased subclone cellular prevalence estimates. Pipeline-estimated cellularity by pipelines using PhyloWGS, PyClone and DPClust also dropped dramatically in correlation with CNA detection tool estimated cellularity once the latter reached 0.75 (TITAN: −0.478–(-)0.163, Battenberg: −0.396–(-)0.021, Spearman’s *ρ*). This appears to lead to the anecdotal observation that high cellularity results from both Battenberg and TITAN could reflect unsuccessful CNA detection, and should be interpreted with caution and perhaps supported by orthogonal evidence. Finally, Battenberg- and TITAN-estimated cellularities showed poor correlation with each other (0.235, Spearman’s *ρ*). As a result, in 12/12 pipelines using either PhyloWGS, PyClone or DPClust, changing the CNA detection tool influenced cellularity estimates more than changing the SNV detection tool.

**Figure 2.**
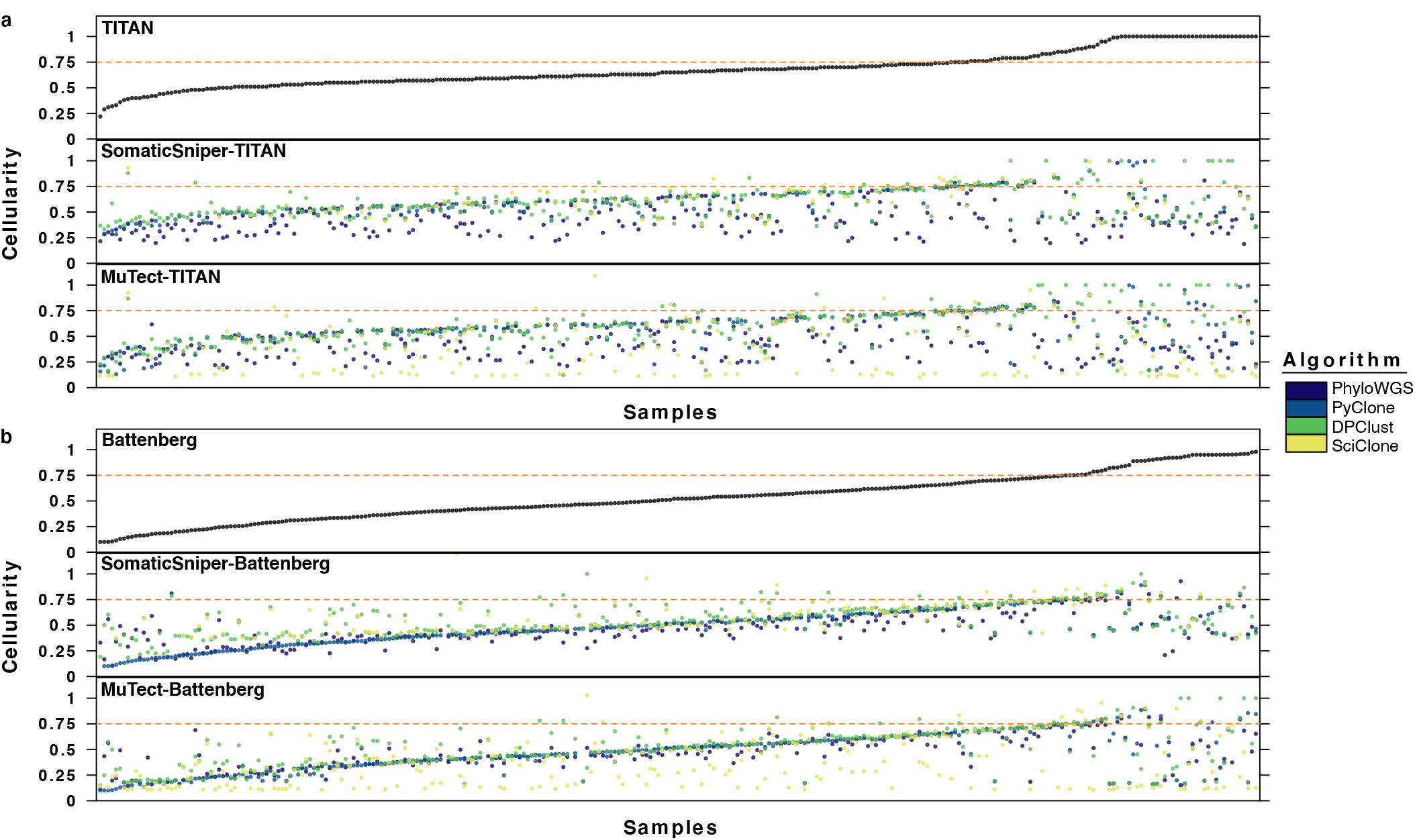
Cellularity Estimates. Cellularity of samples as estimated by the CNA detection tool and by subclonal reconstruction pipelines using the CNA detection tool **a** TITAN and **b** Battenberg. Each dot represents the estimate for a sample and colors delineate subclonal reconstruction algorithms. Mutation detection tool combinations using Battenberg include SomaticSniper-Battenberg and MuTect-Battenberg, and mutation detection tool combinations using TITAN include SomaticSniper-TITAN and MuTect-TITAN. Samples are ordered by cellularity estimates by the CNA detection tool. The horizontal line indicates CNA detection tool estimated cellularity 0.75. TITAN: n=293 biologically independent samples; SomaticSniper-TITAN-PhyloWGS: n=289; SomaticSniper-TITAN-PyClone: n=221; SomaticSniper-TITAN-DPClust: n=293; SomaticSniper-TITAN-SciClone: n=100; MuTect-TITAN-PhyloWGS: n=289; MuTect-TITAN-PyClone: n=221; MuTect-TITAN-DPClust: n=283; MuTect-TITAN-SciClone: n=182. TITAN-estimated cellularity > 0.75: n=73; SomaticSniper-TITAN-PhyloWGS: n=69; SomaticSniper-TITAN-PyClone: n=54; SomaticSniper-TITAN-DPClust: n=73; SomaticSniper-TITAN-SciClone: n=20; MuTect-TITAN-PhyloWGS: n=72; MuTect-TITAN-PyClone: n=53; MuTect-TITAN-DPClust: n=70; MuTect-TITAN-SciClone: n=50. Battenberg: n=293; SomaticSniper-Battenberg-PhyloWGS: n=287; SomaticSniper-Battenberg-PyClone: n=277; SomaticSniper-Battenberg-DPClust: n=291; SomaticSniper-Battenberg-SciClone: n=150; MuTect-Battenberg-PhyloWGS: n=281; MuTect-Battenberg-PyClone: n=262; MuTect-Battenberg-DPClust: n=288; MuTect-Battenberg-SciClone: n=257. Battenberg-estimated cellularity > 0.75: n=47; SomaticSniper-Battenberg-PhyloWGS: n=44; SomaticSniper-Battenberg-PyClone: n=38; SomaticSniper-Battenberg-DPClust: n=47; SomaticSniper-Battenberg-SciClone: n=18; MuTect-Battenberg-PhyloWGS: n=42; MuTect-Battenberg-PyClone: n=37; MuTect-Battenberg-DPClust: n=46; MuTect-Battenberg-SciClone: n=40.

We next assessed if subclonal reconstruction pipelines differed in the number of subclones they predict. For each of the 293 tumours evaluated, up to 16 subclonal reconstruction pipelines were successfully executed, with a median of 14 successful executions (25^th^ quantile [Q1]: 12, 75^th^ quantile [Q3]: 16). Across samples, a median of 7/16 pipelines agreed on the number of subclones predicted (Q1: 6; Q3: 8). The median tumour was predicted to harbor one to three subclones across pipelines (lower range Q1: 1, Q3: 1; upper ranger Q1: 3, Q3: 5), and two randomly-selected successfully-executed pipelines would differ by 1.1 ± 1.3 (mean ± SD) in their predicted number of subclones across samples. These variabilities reflect substantial differences between subclonal reconstruction pipelines. Further, no pair of subclonal reconstruction algorithms consistently produced more similar results across mutation detection tool combinations. Pipelines using SomaticSniper for SNV detection achieved higher levels of agreement across subclonal reconstruction algorithms. All successfully-executed algorithms estimated the same number of subclones in 59.8% of samples in pipelines using SomaticSniper and Battenberg, and in 29.3% of samples in pipelines using SomaticSniper and TITAN, though the agreements were largely driven by concordant monoclonal reconstructions (**Figure 3A-B**). Pipelines using MuTect had much lower levels of agreement across subclonal reconstruction algorithms in pipelines using the same mutation detection tool combination (MuTect-Battenberg: 21.5%, MuTect-TITAN: 12.4%; **Figure 3C-D**), although these results suggest pipelines using SomaticSniper may systematically underestimate subclonal complexity.

**Figure 3.**
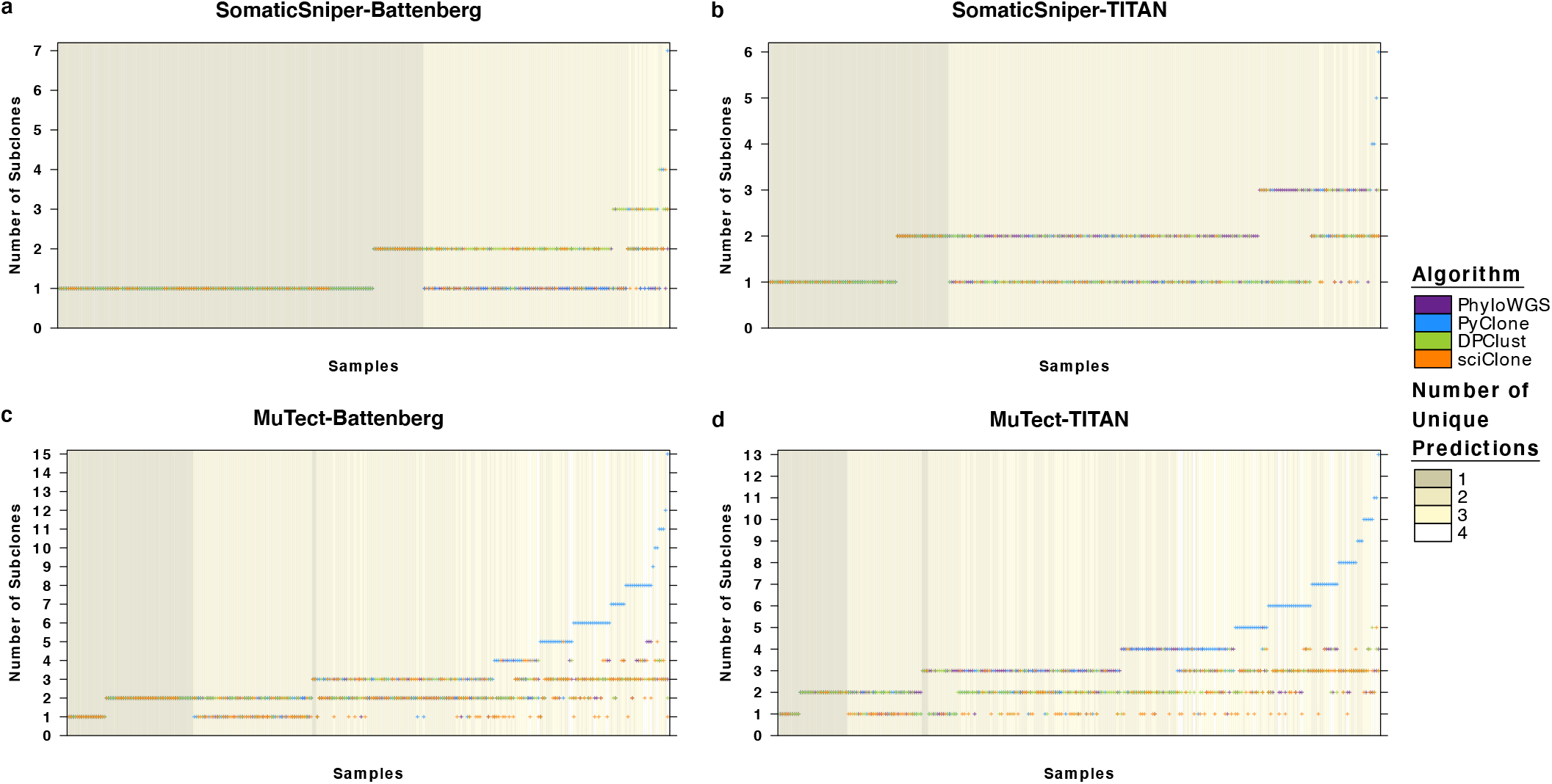
Number of Subclones Detected. Each panel compares the number of subclones predicted for each sample by subclonal reconstruction pipelines using the same mutation detection tool combinations **a** SomaticSniper-Battenberg, **b** SomaticSniper-TITAN, **c** MuTect-Battenberg and **d** MuTect-TITAN. Each marker represents the prediction for a sample by a successful pipeline execution, and the color of the marker represents the subclonal reconstruction algorithm used in the pipeline. In cases where algorithms predicted the same number of subclones, the markers were randomly overlaid. Background color indicates the number of unique subclone number predictions across algorithms that successfully executed for that sample. SomaticSniper-Battenberg: n=291 biologically independent samples; SomaticSniper-TITAN: n=293; MuTect-Battenberg: n=288; MuTect-TITAN: n=290.

To better understand the contribution of mutation detection tools to the discordance in predicted subclonal architectures across pipelines, we compared clonality solutions between pipelines using the same subclonal reconstruction algorithm across mutation detection tool combinations. There are strong interactions between mutation detection tools; for example, predictions by the SomaticSniper-Battenberg-PhyloWGS pipeline agreed poorly with predictions made by other pipelines using PhyloWGS (**Supplementary Figure 2A**). Agreement was highest between the two pipelines using MuTect due to the high number of polyclonal solutions. This overall trend was replicated in pipelines using PyClone, where the SomaticSniper-Battenberg-PyClone pipeline had high agreement with the SomaticSniper-TITAN-PyClone pipeline but differed from pipelines using MuTect (**Supplementary Figure 2B**). DPClust-comprising pipelines using MuTect also predicted high numbers of polyclonal architectures and showed low agreements with other pipelines (**Supplementary Figure 2C**). Finally, results were similar for pipelines using SciClone, with pipelines using the same SNV detection tools achieving the highest agreement (**Supplementary Figure 2D**).

As PhyloWGS is the only one of the four subclonal reconstruction algorithms evaluated that predicts the evolutionary relationship between subclones, we compared the phylogenetic clone trees for each sample as predicted by PhyloWGS-comprising pipelines (**Supplementary Figure 3A**). The most frequently predicted polyclonal architecture was the bi-clonal tree, accounting for 69.8 ± 25.4% (mean ± SD) of polyclonal solutions across pipelines. As multiple phylogenetic clone trees can be inferred from the same data^2,29^, we evaluated prediction stability across the 2,500 Markov chain Monte Carlo (MCMC) iterations of PhyloWGS after burn-in (**Supplementary Figure 3B-E**). Most samples alternated between 1.9 ± 1.2 (mean ± SD) solutions. In 100% of the cases with an alternative phylogeny, the solution alternated at least once between phylogenetic clone trees with different numbers of subclones. Further, when PhyloWGS wavered between solutions that only differed in tree structures (not number of subclones), two alternatives dominated (2.1 ± 0.3, mean ± SD). These data suggest that the uncertainty in phylogenetic clone tree reconstruction comes from the combination of uncertainty from estimating subclone number and resolving their evolutionary relationships.

Taking the consensus across mutation detection tools is a common approach for increasing confidence in mutation detection^30^. We evaluated how subclonal architectures predicted by PhyloWGS-comprising pipelines change when using the union and intersection of detected mutations (**Methods**). MuTect detected significantly more unique SNVs than SomaticSniper (medianUnique SNVs, MuTect = 5,330, medianUnique SNVs, SomaticSniper = 623, p < 2.2 × 10^−16^, Wilcoxon signed-rank test; **Supplementary Figure 4A**). CNAs detected by TITAN and Battenberg were also substantially imbalanced, with a median of 50.2% and 1.2% of the covered genome having uniquely detected CNAs across samples, respectively (p < 2.2 × 10^−16^, Wilcoxon signed-rank test; **Supplementary Figure 4B**). The pipeline using the union of SNVs and the intersect of CNAs predicted clonality with similar skew to the pipeline using the union of both SNVs and CNAs, and the pipeline using the intersection of SNVs and union of CNAs predicted clonality with similar balance to the pipeline using the intersect of both SNVs and CNAs (**Supplementary Figure 4C-F**). This is consistent with our observation that pipeline predictions of complex polyclonal phylogenies using PhyloWGS are primarily driven by large numbers of SNVs detected by MuTect, and complexity in CNAs has a smaller influence on the delineation of cancer cell populations.

Considering the strong influence of SNV detection tools on the number of subclones predicted, we investigated the VAFs and trinucleotide profiles of SNVs detected by MuTect and SomaticSniper. Across all 293 WGS tumour-normal pairs, MuTect-unique SNVs had significantly lower VAFs than those detected only by SomaticSniper or by both tools (median_VAF, MuTect-Unique_ = 9.8%, median_VAF, SomaticSniper-Unique_ = 24.0%, median_VAF, Intersect_ = 28.3%; both p < 2.2 × 10^−16^, Mann-Whitney U-test; **Figure 4A**). This supports the finding that the prediction of a higher number of cancer cell populations is associated with higher numbers of input SNVs with ranging VAFs^15^. SNVs detected by both tools exhibited a trinucleotide profile characterized by Np[C>T]G mutations, while a higher proportion of SomaticSniper-unique SNVs were T>C and MuTect-unique SNVs were characterized by a high proportion of C>A mutations, especially C[C>A]G and T[C>A]G (**Figure 4B-D**). This is suggestive of error profiles related to sequencing or alignment artefacts^31^. As all SNVs detected by SomaticSniper and MuTect were subjected to allow- and deny-list filtering^13,23^ prior to subclonal reconstruction (**Methods**), we also evaluated the effect of filtering on VAFs and trinucleotide profiles. In general, filtering removed low-VAF SNVs, but minimally influenced trinucleotide mutational profiles (**Supplementary Figure 5A-E**).

**Figure 4.**
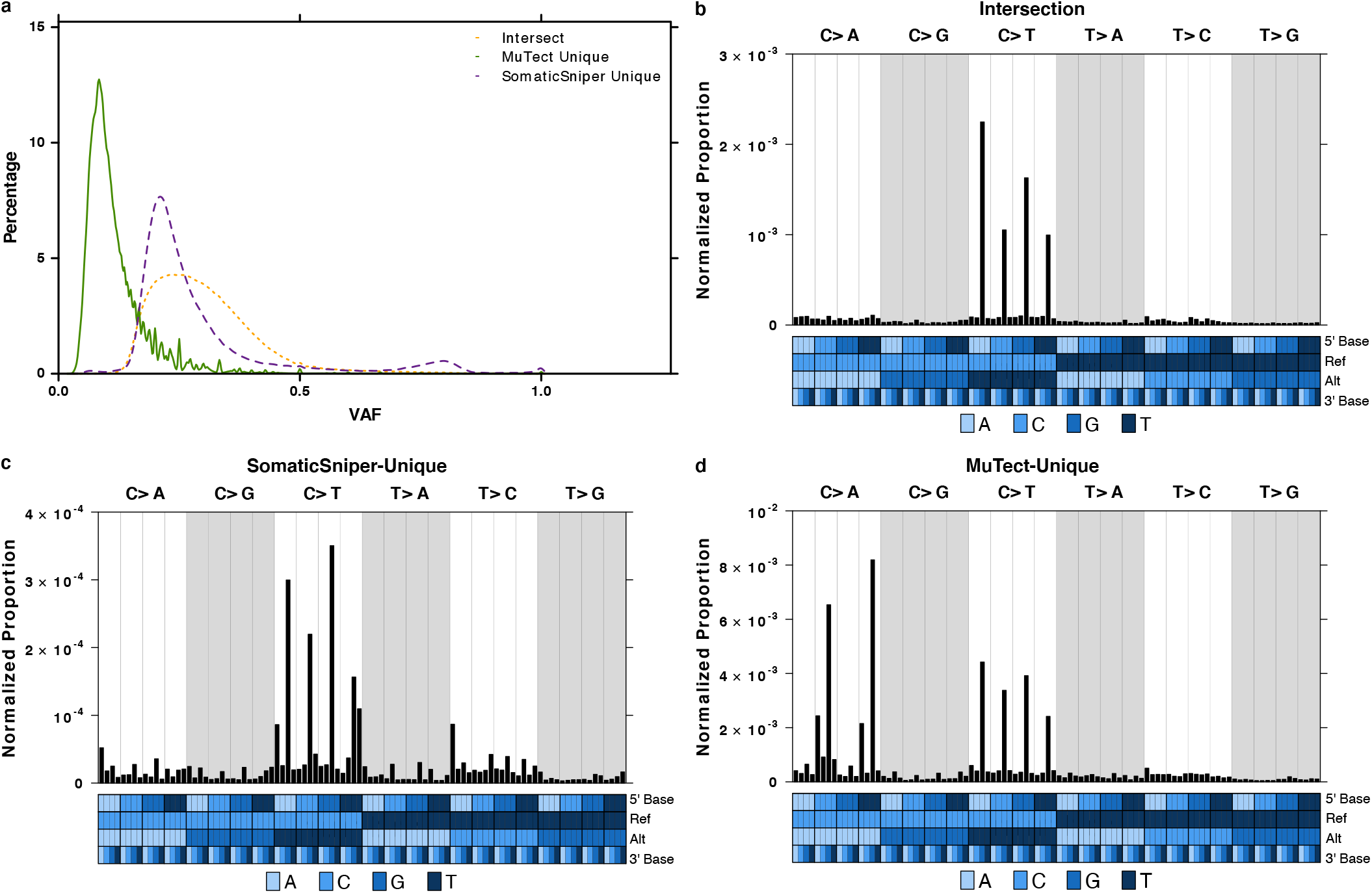
SomaticSniper and MuTect. **a** Density plots of variant allele frequencies for SNVs across all samples that were detected by both SomaticSniper and MuTect (Intersect, short dash orange line), only detected by MuTect (MuTect Unique, solid green line) and only detected by SomaticSniper (SomaticSniper Unique, long dash purple line). VAF, variant allele frequency. **b** Trinucleotide profile of SNVs that were detected by both SomaticSniper and MuTect, where the number of SNVs was normalized by the expected number of each trinucleotide context across the hg19 genome. Trinucleotide profiles for **c** SNVs only detected by SomaticSniper and **d** SNVs only detected by MuTect. Colors in the covariate bar indicate the 5’, reference, alternative and 3’ nucleotides in each trinucleotide context. Ref, reference nucleotide; Alt, alternative nucleotide of variant. Number of SNVs across all samples in Intersect: n=453,855 independent observations; SomaticSniper Unique n=198,623; MuTect Unique n=2,801,546.

### Consistency of SNV Clonality

One goal of subclonal reconstruction is to time when individual mutations occurred during tumour evolution. We therefore compared clonal and subclonal SNV identification for the same set of 293 WGS samples across sixteen pipelines for subclonal reconstruction. As expected from the different types of SNVs leveraged for subclonal reconstruction, algorithms were highly discordant in the numbers of SNVs identified as clonal or subclonal. In samples where the subclonal reconstruction algorithm was successfully executed across all four mutation detection tool combinations, DPClust used and timed the most SNVs on average (2,941 ± 3,929, mean ± SD; **Figure 5A**), followed by PhyloWGS (2,473 ± 1,662, **Figure 5B**), PyClone (1,738 ± 1,580, **Figure 5C**) and SciClone (178 ± 480, **Figure 5D**). As expected from the influence of MuTect on the prediction of subclonal clusters, its use was associated with the identification of an order of magnitude more subclonal SNVs, but similar numbers of clonal SNVs as with use of SomaticSniper.

**Figure 5.**
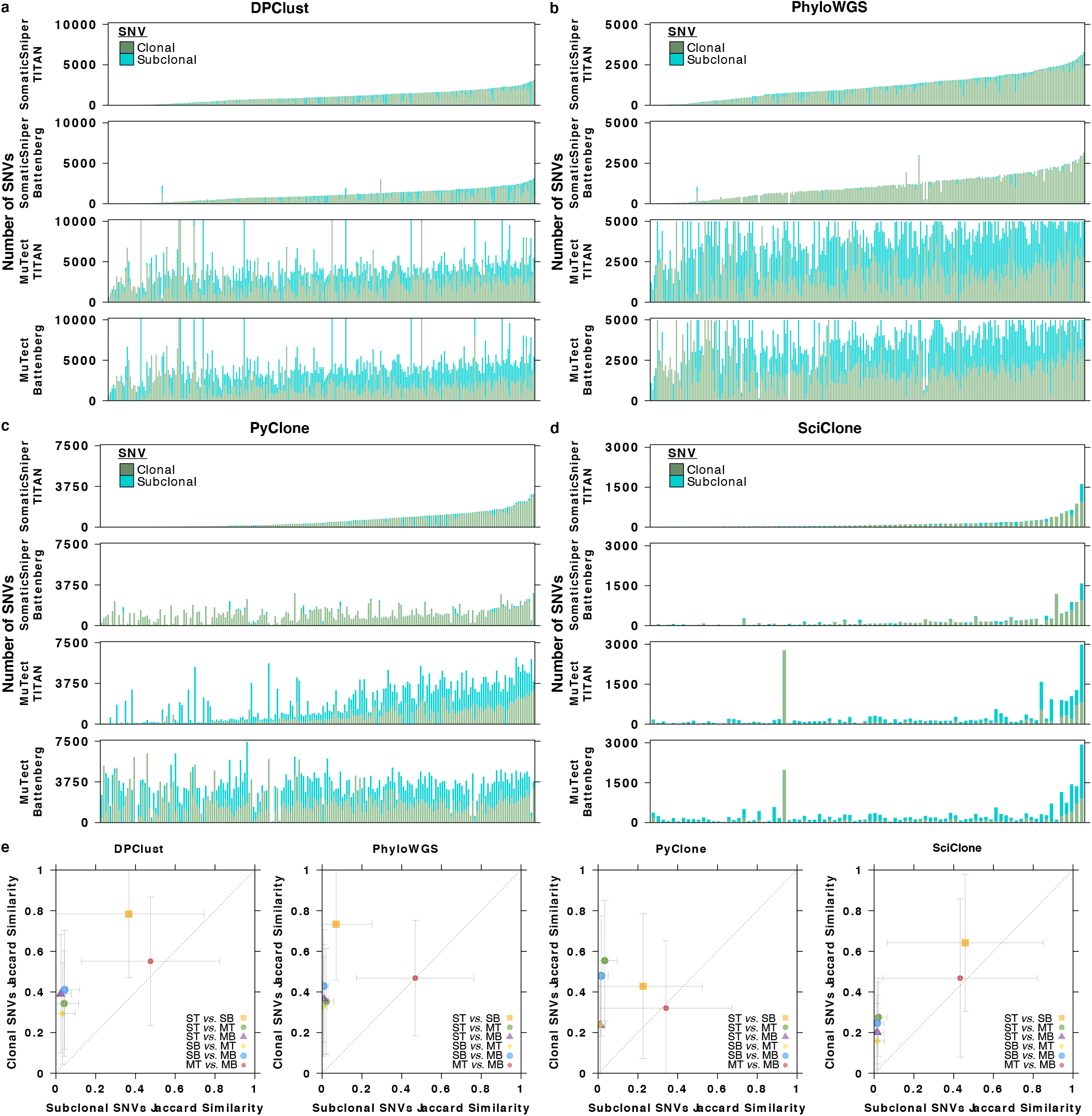
Clonal and Subclonal SNVs. Total number of clonal and subclonal SNVs identified by pipelines using **a** DPClust, **b** PhyloWGS, **c** PyClone and **d** SciClone. Each stacked bar represents one sample and samples are ordered based on the total number of SNVs identified by the pipeline using SomaticSniper and TITAN. Color of the stacked bar reflects the clonality of the SNVs it represents (clonal green, subclonal blue). Only samples with successful executions across all four pipelines using the subclonal reconstruction algorithm are presented. DPClust: n=281 biologically independent samples; PhyloWGS n=274; PyClone n=200; SciClone n=86. **e** Jaccard index of pipeline-identified clonal and subclonal SNVs. Each marker (delineated by shape and color) represents a pipeline pair that is compared, and the *x*- and *y*- axis show subclonal and clonal mean SNV Jaccard indices across samples shared between the pipeline pair, respectively, with error bars indicating one standard deviation. Dashed diagonal line represents the *y* = *x* line. ST, SomaticSniper-TITAN; MT, MuTect-TITAN; SB, SomaticSniper-Battenberg; MB, MuTect-Battenberg. DPClust ST *vs*. SB: n=291 independent observations; DPClust ST *vs*. MT: n=283; DPClust ST *vs*. MB: n=288; DPClust SB *vs*. MT: n=281; DPClust SB *vs*. MB: n=288; DPClust MT *vs*. MB: n=281; PhyloWGS ST *vs*. SB: n=288; PhyloWGS ST *vs*. MT: n=290; PhyloWGS ST *vs*. MB: n=283; PhyloWGS SB *vs*. MT: n=285; PhyloWGS SB *vs*. MB: n=281; PhyloWGS MT *vs*. MB: n=283; PyClone ST *vs*. SB: n=216; PyClone ST *vs*. MT: n=209; PyClone ST *vs*. MB: n=202; PyClone SB *vs*. MT: n=213; PyClone SB *vs*. MB: n=256; PyClone MT *vs*. MB: n=212; SciClone ST *vs*. SB: n=88; SciClone ST *vs*. MT: n=96; SciClone ST *vs*. MB: n=92; SciClone SB *vs*. MT: n=116; SciClone SB *vs*. MB: n=147; SciClone MT *vs*. MB: n=174. Source data are provided as a Source Data file.

To further evaluate how mutation detection tools affect the timing of SNVs, we calculated the Jaccard index of clonal SNVs identified between all pipeline pairs using the same subclonal reconstruction algorithm, and the same for subclonal SNVs (**Figure 5E**). In PhyloWGS-comprising pipelines, clonal SNV identifications were in high agreement (mean Jaccard index ± SD: 44.6 ± 30.2%) but subclonal SNV identifications were significantly less so (10.0 ± 22.4%; p < 2.2 × 10^−16^, Wilcoxon signed-rank test), particularly between pipelines using different SNV detection tools. The results were similar for other algorithms: DPClust (clonal Jaccard index: 46.3 ± 33.2%, mean ± SD; subclonal: 15.4 ± 27.5%), PyClone (clonal: 38.0 ± 32.2%; subclonal: ± 21.6%) and SciClone (clonal: 33.3 ± 31.5%; subclonal: 14.8 ± 29.3%). Overall, we observe diversity in SNV profiles and clonality predictions across pipelines, with extensive diversity in subclonal SNV profiles associated with mutation detection tools.

To better understand how subclonal reconstruction algorithms differ in their prediction of SNV clonality, we next focused on SNVs identified as clonal across all pipelines using the same mutation detection tool combination. For each sample, we assessed the overlap in clonal SNVs identified by each pipeline and found only a small percentage of SNVs per sample that were unanimously identified as clonal: SomaticSniper-TITAN: 2.0 ± 5.8%, SomaticSniper-Battenberg: 3.8 ± 8.0%, MuTect-TITAN: 0.5 ± 2.0%, MuTect-Battenberg: 1.0 ± 3.1% (mean ± SD; **Supplementary Figure 6A-D**). Nevertheless, most SNVs were identified as clonal by more than one algorithm (SomaticSniper-TITAN: 77.4 ± 25.2%, SomaticSniper-Battenberg: 91.9 ± 17.8%, MuTect-TITAN: 48.3 ± 30.9%, MuTect-Battenberg: 71.9 ± 28.5%). Pipelines using Battenberg were characterized by large overlaps in clonal SNV identifications between PhyloWGS, DPClust and PyClone (SomaticSniper-Battenberg: 63.2 ± 34.3%, MuTect-Battenberg: 46.2 ± 33.5%). Pipelines using TITAN were characterized by modest overlaps between these three, but stronger overlap between PhyloWGS and DPClust (SomaticSniper-TITAN: 42.9 ± 35.4%, MuTect-TITAN: 27.1 ± 26.1%). Given the lack of correlation between subclonal reconstruction algorithms in estimating the number of subclones present in a sample, this could suggest that disagreements between subclonal reconstruction algorithms mostly fall in defining the subclonal populations.

### Consistency of CNA Clonality

We also evaluated the influence of mutation detection tools on clonal and subclonal CNA identification. We focused on PhyloWGS, as it was the only algorithm considered here that co-clusters SNVs and CNAs. Previous work on this cohort using the SomaticSniper-TITAN-PhyloWGS pipeline identified four clonal CNA subtypes and three subclonal CNA subtypes^13^, so we first evaluated their robustness across pipelines. In general, clonal subtypes were more robust to pipeline changes, while subclonal subtypes were less so (**Supplementary Figure 7A-B, Supplementary Data 21-32**). Pipelines employing the same CNA detection tool also had more similar profiles then those using different ones.

We next assessed the agreement of these pipelines in their identification of clonal and subclonal CNAs. We calculated the Jaccard index of the identification of 1.0 Mbp genomic bins with CNAs between pipeline pairs, where the direction of aberration (*i.e.,* gain *vs.* loss) must match to be considered as an agreement. We found significantly greater agreement for clonal CNAs compared to subclonal CNAs across all pipeline pairs (mean clonal Jaccard index ± SD: 50.5 ± 21.1%, subclonal Jaccard: 15.6 ± 21.8%; all p < 2.2 × 10^−16^, Wilcoxon signed-rank test; **Supplementary Figure 7C**). Pipelines using the same CNA detection tool tended to agree, although divergence was expected because the reconstructed clonality of CNA segments can be influenced by the VAFs of SNVs in the segment. By contrast, pipelines with different CNA detection tools had less clonal and little subclonal agreement. Thus, for both SNVs and CNAs, clonal mutational landscapes were relatively invariant to pipeline but subclonal ones were not.

### Impact of Reconstruction Variability on Downstream Analyses

Given these differences in SNV and CNA clonality prediction across pipelines, we sought to understand how they might influence the timing of mutations in cancer driver genes. These genes are of particular relevance as they can be actionable as predictive or prognostic biomarkers. We examined the clonality of mutations in five genes driven by recurrent somatic SNVs (*ATM*, *FOXA1*, *MED12*, *SPOP* and *TP53*) and eight driven by recurrent somatic CNAs (*CDH1*, *CDKN1B*, *CHD1*, *MYC*, *NKX3-1*, *PTEN*, *RB1* and *TP53)* in localized prostate cancer^13,23^. Focusing on PhyloWGS-comprising pipelines, these driver events were overwhelmingly predicted to occur early (*i.e.* clonally) during tumour evolution, with 87.2 ± 16.8% (mean ± SD) of SNV and 91.5 ± 6.4% of CNA driver mutations identified as clonal across pipelines (**Supplementary Figure 8A-B)**. There was also broad consensus in these predictions: when a clonal SNV was identified in a specific driver gene and sample by any single pipeline, all four pipelines identified a clonal SNV in that driver gene in the same sample in 39.5 ± 22.5% of cases (mean ± SD). CNAs showed even higher consensus (50.4 ± 14.8%; **Supplementary Figure 8C**). One outlier was *MED12*, where there was disagreement across pipelines with the same SNV detection tools: since *MED12* is located on the × chromosome and Battenberg does not generate copy number status for regions of uncertainty and the sex chromosomes, its mutations were disregarded during subclonal reconstruction because PhyloWGS only considers SNVs with overlapping copy number status.

We then evaluated how CNA clonality predictions would affect the identification of genes as significantly differentially mutated clonally *vs.* subclonally. Within each pipeline we determined whether each 1.0 Mbp genomic bin had different proportions of gains and losses clonally and subclonally (FDR < 0.05, Pearson’s *χ*^2^ Test, clonal: loss, neutral, gain *vs.* subclonal: loss, neutral, gain; **Methods**). The number of genes in regions with CNAs occurring statistically more frequently early or late differed dramatically across PhyloWGS-comprising pipelines (MuTect-TITAN: 5,344; SomaticSniper-TITAN: 5,198; MuTect-Battenberg: 1,498; SomaticSniper-Battenberg: 339). A consensus set of 339 genes showed a bias in timing in all pipelines as preferentially mutated clonally (**Supplementary Figure 9A**, **Supplementary Data 33-36**). These genes were enriched for *TP53*-based regulation of death receptors, *TRAIL* signaling and natural killer cell mediated cytotoxicity (FDR < 0.05; **Supplementary Figure 9B**).

To evaluate whether pipeline differences could influence the accuracy of biomarkers, we focused on biochemical relapse after definitive local therapy. Previous work has identified clonality to be prognostic in this setting, both independently and when combined with an established multi-modal (CNA, SNV, SV and methylation) gene-specific biomarker^13,23^. Discretization by clonality (monoclonal *vs.* polyclonal) only stratified patients by outcome in the SomaticSniper-TITAN-PhyloWGS pipeline (p = 0.004, log-rank test; **Supplementary Figure 10A**), but not any other (all p > 0.05, log-rank test; **Supplementary Figure 10B-P**). The unified biomarker integrating clonality and a multi-modal biomarker achieved prognostic value in more pipelines (p < 0.05 in 14/16 models, log-rank test; **Supplementary Figure 11A-P**), with concordant trends across all pipelines. Thus, the prognostic effect size of clonality in prostate cancer is smaller than the technological effect size in this cohort, with a clinical signal smaller than technical variance. As a result, the translational potential of clonality in localized prostate cancer is improved when it is integrated with complementary gene-specific biomarker information.

### Comparing Reconstructions using Single and Multiple Regions

Our analyses of a large cohort of single-sample reconstructions highlight large inter-pipeline differences in the determination of subclonal architecture and prediction of mutation clonality. To better relate these results to the ground-truth, we focused on a set of ten localized prostate cancers where samples from multiple regions of the tumour were available (30 genomes in total, ranging from 2-4 per patient). These data allowed us to directly compare single-region to multi-region reconstructions using PhyloWGS and PyClone, providing an estimate of the extent to which the former underestimates true clonal complexity.

We first quantified the differences in the number of subclones predicted from single-region and multi-region reconstructions of the ten tumours (**Supplementary Data 37-48**). Multi-region reconstructions predicted more subclones than single-region reconstructions in pipelines using PhyloWGS: 4.6 ± 2.4 (mean ± SD) subclones were predicted with multi-region reconstructions while 2.0 ± 0.9 subclones were predicted with single-region reconstructions (**Figure 6A**). This difference was not seen in pipelines using PyClone (multi-region reconstructions: 2.2 ± 1.7, single-region reconstructions: 2.3 ± 2.0), likely due to the constraint that only mutations present in all samples are used for multi-region reconstruction (**Figure 6B**). These data suggest that the typical single-sample reconstruction identifies fewer than half of the subclones present in the tumour, and this could very well be a lower-bound estimate because of the limited sequencing depth and spatial sampling of this cohort. On the other hand, multi-sample reconstructions also predicted significantly more subclones within the index lesion sample compared to single-sample reconstruction of the index lesion alone in pipelines using PhyloWGS (mean number of subclones in index lesion from multi-region reconstruction ± SD: 2.6 ± 1.5, from single-region reconstruction: 1.9 ± 0.9; p = 2.4 × 10^−4^, Wilcoxon signed-rank test; **Supplementary Figure 12A**), but not those using PyClone (multi-region reconstruction mean ± SD: 2.2 ± 1.7, single-region reconstruction: 2.5 ± 2.5; p ≈ 1, Wilcoxon signed-rank test; **Supplementary Figure 12B)**. Together this suggests that single-region reconstructions are limited by spatial sampling from fully resolving the intra-tumoural heterogeneity of both the overall tumour and the sampled region, for example due to cases where subclones appear with the same CCF and are thus indistinguishable from single-region reconstructions alone^32^.

**Figure 6.**
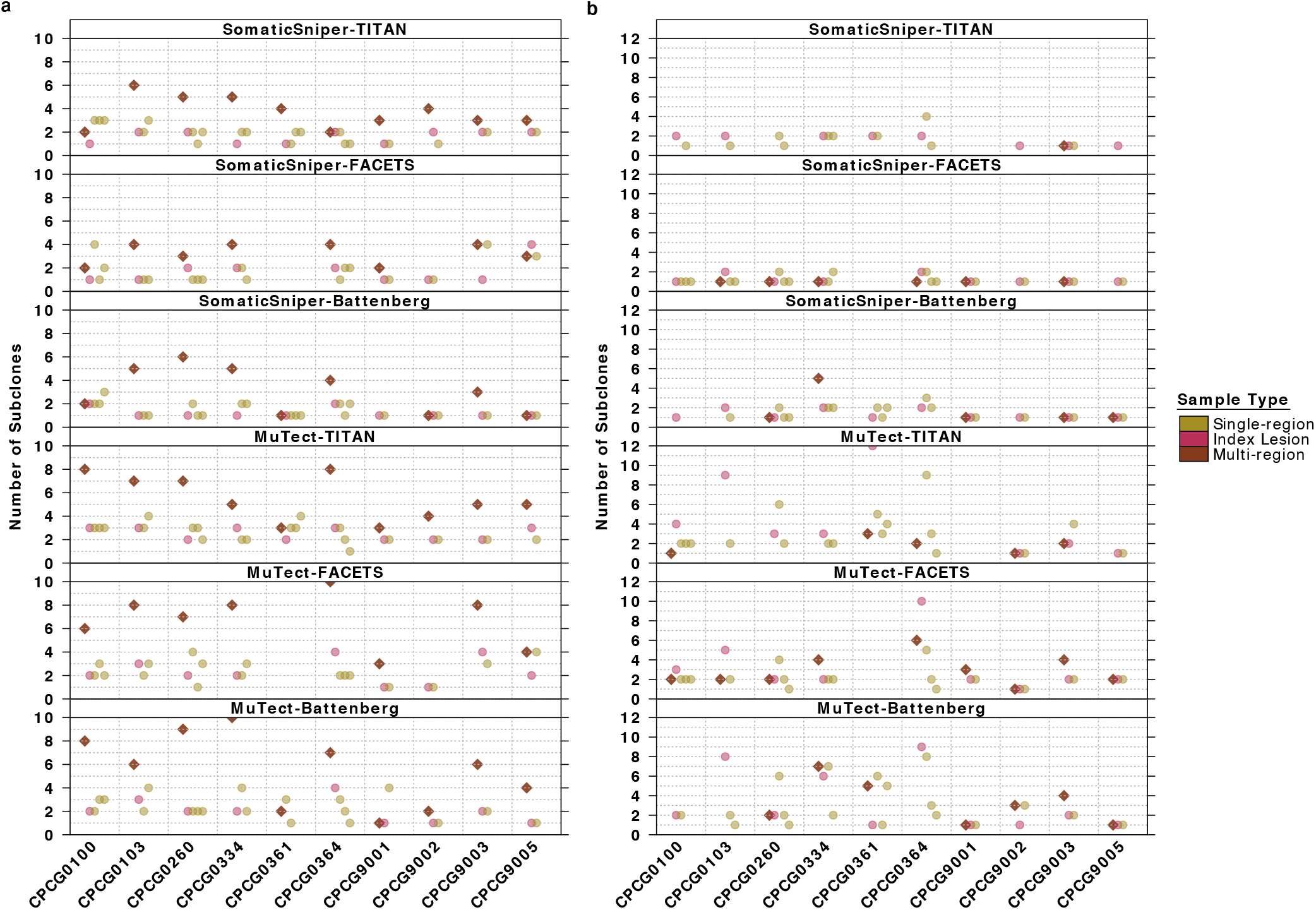
Single- and Multi-Region Subclone Number Prediction. **N** umber of subclones predicted for each tumour from multi-region reconstruction (brown diamond) and reconstructions of each of the individual regions (circle), including the index lesion (pink circle) by pipelines using **a** PhyloWGS and **b** PyClone. Missing values indicate a failed reconstruction. Multi-region reconstruction SomaticSniper-TITAN-PhyloWGS: n=10 biologically independent samples; SomaticSniper-FACETS-PhyloWGS: n=8; SomaticSniper-Battenberg-PhyloWGS: n=9; MuTect-TITAN-PhyloWGS: n=10; MuTect-FACETS-PhyloWGS: n=8; MuTect-Battenberg-PhyloWGS: n=10; SomaticSniper-TITAN-PyClone: n=1; SomaticSniper-FACETS-PyClone: n=6; SomaticSniper-Battenberg-PyClone: n=5; MuTect-TITAN-PyClone: n=5; MuTect-FACETS-PyClone: n=9; MuTect-Battenberg-PyClone: n=7. Single-region reconstruction SomaticSniper-TITAN-PhyloWGS: n=30; SomaticSniper-FACETS-PhyloWGS: n=26; SomaticSniper-Battenberg-PhyloWGS: n=30; MuTect-TITAN-PhyloWGS: n=30; MuTect-FACETS-PhyloWGS: n=26; MuTect-Battenberg-PhyloWGS: n=28; SomaticSniper-TITAN-PyClone: n=18; SomaticSniper-FACETS-PyClone: n=26; SomaticSniper-Battenberg-PyClone: n=25; MuTect-TITAN-PyClone: n=25; MuTect-FACETS-PyClone: n=25; MuTect-Battenberg-PyClone: n=28.

We next sought to determine the extent of variability in SNV clonality predictions between single-region and multi-region reconstructions. We identified SNVs that were predicted be the same clonality (clonal or subclonal) in both single- and multi-region reconstructions (‘Match in Multi and Single’). For SNVs with mismatched clonality, we further categorized them as clonal in multi-region reconstruction and subclonal in single-region reconstruction (‘Clonal in Multi-region’) or *vice versa* (‘Subclonal in Multi-Region’), or SNVs that were uniquely considered in single-region reconstructions (‘Unique in Single-region’) or multi-region reconstructions (‘Unique in Multi-region’). The last category of SNVs is unique to PhyloWGS as it is able to consider SNVs unique to individual samples for multi-region analysis. SNV clonality predictions matched less than half the time for pipelines using PhyloWGS (32.2 ± 24.5%, mean ± SD; **Figure 7A**). Pipelines using PyClone achieved modestly higher clonality agreement, perhaps due to the smaller number of subclones predicted in multi-region reconstructions and the lack of multi-region unique SNVs (38.6 ± 25.4%; **Figure 7B**). Mismatched SNVs tended to be clonal in single-region reconstructions and subclonal in multi-region reconstructions, as expected. Consistent with simulations^33^ and previous observations, multi-region reconstructions are able to better define subclonal populations of cells by identifying and disambiguating those missed or merged by single-region sampling.

**Figure 7.**
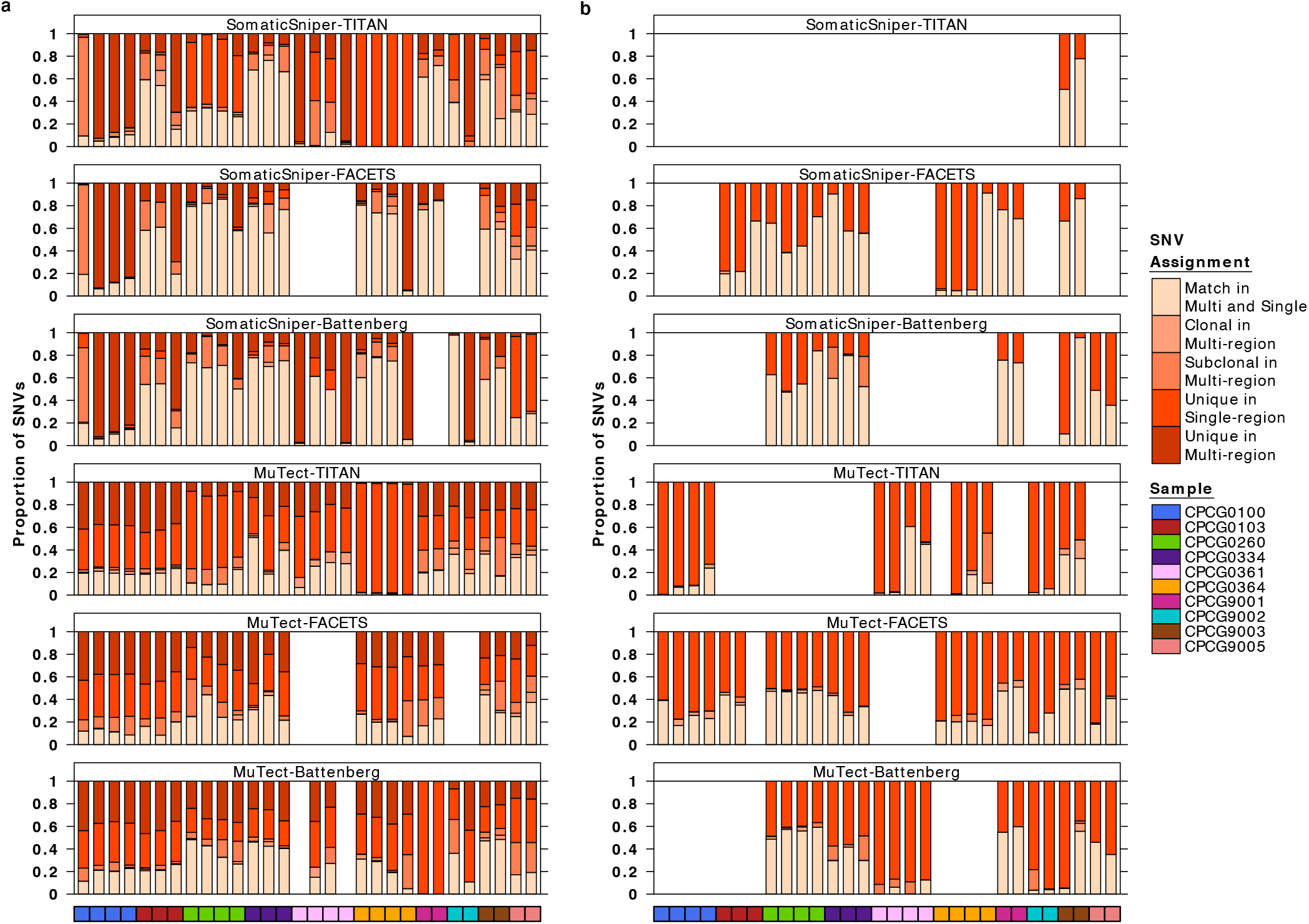
Single- and Multi-Region SNV Clonality Prediction. Comparison of the clonality of SNVs identified by single-region and multi-region reconstructions by pipelines using **a** PhyloWGS and **b** PyClone. Each stacked bar represents a single-region and covariate bar color indicates the identity of the sample. Missing bars indicated failed reconstructions, either single- or multi-region. SNVs were grouped into five categories by color of stacked bar plot: ‘Match in Multi and Single’ if the SNV was predicted to be the same clonality in single- and multi-region reconstructions, ‘Clonal in Multi-region’ if the SNV was clonal in multi-region reconstruction but subclonal in single-region reconstruction, and ‘Subclonal in Multi-region’ if *vice versa*. If a SNV was only analyzed in single-region reconstruction, it was ‘Unique in Single-region’, while SNVs only analyzed in multi-region reconstruction were ‘Unique in Multi-region’. SomaticSniper-TITAN-PhyloWGS: n=30 biologically independent samples; SomaticSniper-FACETS-PhyloWGS: n=24; SomaticSniper-Battenberg-PhyloWGS: n=28; MuTect-TITAN-PhyloWGS: n=30; MuTect-FACETS-PhyloWGS: n=24; MuTect-Battenberg-PhyloWGS: n=28. SomaticSniper-TITAN-PyClone: n=2; SomaticSniper-FACETS-PyClone: n=18; SomaticSniper-Battenberg-PyClone: n=13; MuTect-TITAN-PyClone: n=15; MuTect-FACETS-PyClone: n=25; MuTect-Battenberg-PyClone: n=19. Source data are provided as a Source Data file.

We also examined the agreement between single-region and multi-region reconstruction CNA clonality predictions in pipelines using PhyloWGS (**Supplementary Figure 13**). Agreements were similarly variable, with less than half of CNAs matching in clonality between the single- and multi-region reconstructions and extensive variance across samples and pipelines (35.2 ± 31.5%, mean ± SD). As with SNVs, mismatches mostly involved clonal CNAs in single-region reconstructions that were identified as subclonal in multi-region reconstructions.

To better understand this sampling bias, we analyzed how well the clonal population of the index lesion from single-region reconstruction represents the clonal population of the entire tumour. In PhyloWGS-comprising pipelines, multi-region reconstruction often showed that SNVs identified as clonal in the index lesion were actually subclonal (**Supplementary Figure 14A)**. Nevertheless, the majority of single-region clonal SNVs were truly clonal in multi-region reconstruction (66.6 ± 29.8%, mean ± SD). As before, pipelines using PyClone showed much higher agreement (91.4 ± 23.3%), likely because of the large number of excluded SNVs (**Supplementary Figure 14B**). A similar analysis of subclonal SNVs showed that, as expected, only a small proportion of subclonal SNVs defined by single-region reconstructions of the index lesion was clonal in multi-region reconstructions in pipelines using PhyloWGS and MuTect (12.2 ± 17.5%, mean ± SD). In contrast, multi-region reconstruction pipelines using PhyloWGS and SomaticSniper predicted many subclonal SNVs from single-region reconstructions as clonal (55.2 ± 40.3%). This highlights a potential limitation of multi-region subclonal reconstruction algorithms with a need for shared SNVs or CNAs.

## Discussion

It is difficult to benchmark the accuracy of subclonal reconstruction methodologies since a robust gold-standard experimental dataset does not yet exist. Simulation frameworks are of great value, but might not fully recapitulate the error-profiles and signal-biases of real data^34^. To evaluate the technological variability in estimating aspects of subclonal architecture, we evaluated 293 tumours using sixteen pipelines. These data provide an experimental lower-bound on the algorithmic variability of tumour subclonal reconstruction in a large high-depth whole-genome sequencing cohort, at least for a single cancer type and stage. We complement these data by assessing eighteen subclonal reconstruction pipelines across a set of 10 multi-region tumours to estimate the degree to which single-sample reconstructions underestimate clonal complexity the full tumour.

Subclonal reconstruction algorithms differ substantially in their prediction of subclonal architecture across all mutation detection tool combinations, with no pair of algorithms consistently achieving similar results in cellularity estimates, prediction of subclone number and assignment of mutation clonality. While the subclonal CNA detection tool used mostly influenced cellularity estimates but no other aspects of subclonal architecture, large differences were driven by changing the SNV detection approach. Differences between SNV detection tools led to major divergences in subclonal reconstruction: pipelines using MuTect found extensive subclonal diversity, at least partly due to the greater number of low VAF mutations detected. SNV detection benchmarking efforts^31^ could aid in the further characterization of the error profiles of SNV detection tools and optimize parameter tuning to improve subclonal reconstruction. Future studies might benefit from merging multiple subclonal reconstruction pipelines, for example to provide a potential envelope of upper and lower bounds on different features of the reconstruction.

The potential translational and clinical impact of these technical variabilities is considerable. For example, technological differences between analysis pipelines were larger than the effect size of the association between evolutionary complexity and patient survival. This suggests that estimates of technical variability should be provided for analyses dependent on subclonal architecture, such as in studies mapping evolutionary and migration trajectories between primary and metastatic tumours. Studies identifying clonal and especially subclonal driver mutations should be interpreted with such variability estimates as reference since subclonal mutational landscapes were found to be especially vulnerable to pipelines changes when clonal ones were less so. Articulating how these algorithmic differences relate to the clinical effect-size will greatly improve interpretability of these types of data.

Future studies also need to carefully consider the failure-rates of different reconstruction algorithms, as algorithms leveraging clonal or neutral copy number regions might not be suitable for tumour types characterized by large numbers of CNAs and might call for specific CNA detection strategies. Computational failures are problematic for clinical applications and, in combination with the substantive computational requirements that scale with the number of mutations, could be problematic for cancer types characterized by a high mutational burden.

Our evaluation of subclonal reconstruction using data from spatially distinct regions of tumours found that reconstructions relying on a single sample systematically underestimated the number of subclones in a tumour. Input constraints and non-exhaustive sequencing depth and spatial sampling in multi-region reconstructions also suggest that the current level of underestimation is only the lower-bound. This is in line with previous work in kidney cancer^6,11^. These data also agree with previous work showing the distinct mutational profiles of prostate cancer samples from spatially distinct regions of the same tumour^8^ and reinforces the hypothesis that sufficient sampling will uncover multiple subclones in nearly all cancers. It also suggests that strategies for robust multi-region-aware subclonal mutation detection would be a significant benefit to subclonal reconstruction analyses.

Larger datasets are necessary to better evaluate the performance of subclonal reconstruction methodologies. While simulated data is valuable^34^, single-cell sequencing datasets will likely significantly improve the evaluation of ground truth for subclonal reconstruction algorithms in patient samples. In the meantime, this work involving a large clinical cohort will aid in refining subclonal reconstruction methods and provide guidance for evaluating the subclonal architecture of cancer samples.

## Methods

### Patient Cohort

We aggregated a retrospective cohort of localized prostate tumours with patient consent and Research Ethics Board approval from published datasets, with whole-genome sequencing of tumour samples and matched blood-based normal samples^13,23,24,35–38^. The cohort includes 293 patients with tumour samples from the index lesion and 10 patients with multiple samples from intraductal carcinoma and juxtaposed adjacent invasive carcinoma. For patients receiving radiotherapy, the index tumour was identified on transrectal ultrasound and sampled by needle biopsies (TRUS-Bx) and was deemed the largest focus of disease that was confirmed pathologically. A fresh-frozen needle core ultrasound-guided biopsy to this index lesion was obtained for macro-dissection. For patients receiving surgery, the index tumour was identified macroscopically by a GU expert pathologist at the point of surgery and later sampled and biobanked. A fresh-frozen tissue specimen from the index lesion was then obtained from macro-dissection. Details of the patient cohort have been described previously^13,24^.

We focused on patients with clinical intermediate-risk disease as defined by NCCN, with intermediate-risk factors (T2b or T2c disease, ISUP Grade Group 2 or 3 or pre-treatment prostate specific antigen (PSA) serum levels between 10-20 ng/mL). All patients received either precision image-guided radiotherapy or radical prostatectomy with no randomization or classification and were hormone-naive at time of therapy. Four patients in the multi-region sequencing cohort carried germline BRCA2 mutations and had formalin-fixed paraffin-embedded tissues instead of fresh-frozen (CPCG9001, CPCG9002, CPCG9003, CPCG9005). Sample regions suitable for macro-dissection (tumour cellularity > 70%) were marked by genitourinary pathologists and manually macro-dissected, followed by DNA extraction and sequencing.

### Whole genome sequencing data analysis

Protocols for whole-genome sequencing data generation and processing have been previously described^13,23,24^. Briefly, raw sequencing reads from the tumour and normal samples were aligned against human reference genome build hg19 using bwa-aln (v0.5.7)^39^. Lane-level BAMs from the same library were merged and duplicates were marked using picard (v1.92). Local realignment and base quality recalibration were performed together for tumour/normal pairs using GATK (v.2.4.9)^40^. Tumour and normal sample-level BAMs were extracted separately, had headers corrected with SAMtools (v0.1.9)^41^ and were indexed with picard (v1.107). ContEst (v1.0.24530)^42^ was used to estimate lane-level and sample-level sample mix-up and lane-level cross-individual contamination on all sequences, with no significant contaminated detected.

### Tumour Somatic Mutation Assessment

We detected subclonal copy number aberrations from whole-genome sequencing data using Battenberg (v2.2.6)^7^, TITAN (v1.11.0)^25^ and FACETS (v0.5.14)^28^. First, Battenberg (v2.2.6) was installed with underlying ASCAT (v2.5)^43^ using the installation and running wrapper cgpBattenberg (v3.1.0). Required reference files were downloaded as instructed in https://github.com/Wedge-Oxford/battenberg and further required data files were generated as instructed in https://github.com/cancerit/cgpBattenberg. An ignore file was created for the genome assembly hg19 to exclude all chromosomes not in 1-22. Battenberg (v2.2.6) was run with -gender of XY for male patients and -t of 14 to run using 14 threads, and otherwise default parameters. The resulting primary solution was subjected to manual refitting in situations meeting the following criteria: 1) the solution involved a high copy number segment with high BAF and low logR, indicating an unrecognized homozygous loss event, 2) nearly all copy number aberrations were subclonal, 3) there were unreasonably high copy numbers up to infinity. Refitting was performed until the concerns for refitting were resolved or for three attempts after which the original solution was accepted. The CNAs obtained from the primary solution, along with tumour cellularity and ploidy were used for further analysis. We have described subclonal copy number analysis using TITAN (v1.11.0) previously in detail^13^. Briefly, TITAN (v1.11.0) was run through the Kronos (v1.12.0)^44^ pipeline for whole-genome sequence preprocessing and subclonal copy number assessment. GC and mappability files for bias correction were prepared using HMMcopy (v0.1.1) and bowtie (v2.2.6)^45^ on the hg19 reference genome. Heterogeneous positions in the sequence data were identified by MutationSeq (v4.3.7)^46^ using known dbSNP sites from GATK (v2.4.9). For each whole-genome sequence, TITAN (v1.11.0) made predictions of the existence of one to five subclones based on the given input numClusters and the solution with the lowest S_Dbw validity index^25^ was used to obtain the cellularity, ploidy and subclonal CNAs for downstream analysis. Finally, to prepare inputs for subclonal copy number assessment by FACETS (v0.5.14), the accompanying snp-pileup (v434b5ce) algorithm was installed with underlying htslib (v1.9)^41^. A SNP location VCF file was downloaded as instructed for hg19 with SNP version b151 and human genome build version GrCh37p13 from ftp://ftp.ncbi.nlm.nih.gov/snp/organisms/human_9606_b151_GRCh37p13/VCF/00-common_all.vcf.gz, and snp-pileup (v434b5ce) was run using developer recommended parameters (-g -q15 -Q20 -P100 -r25,0). All FACETS (v0.5.14) runs used the seed 1234 and default parameters for all steps, except for procSample where the developer recommended parameter cval = 150 was used.

We used MuTect (v1.1.4)^27^ and SomaticSniper (v1.0.2)^26^ for the detection of somatic single nucleotide variants from whole-genome sequencing data. MuTect was run to obtain candidate SNVs with dbSNP138^47^, COSMIC (v66)^48^ and default parameters except the -tumor_lod option (tumor limit of detection). The -tumor_lod option was set to 10 to increase the stringency of detection. Outputs that contained REJECT were filtered out and the remaining SNVs were used for downstream analysis. Details for SomaticSniper (v1.0.2) variant detection have been described previously^23^. In short, SomaticSniper (v1.0.2) was used to identify candidate SNVs with default parameters except the -q option (mapping quality threshold), which was set to 1 as per developer recommendation. Candidate SNVs were filtered through standard and LOH filtering using a pileup indel file generated on the sequence data using SAMtools (v0.1.9)^41^, bam-readcount filtering and false positive filtering. Only high confidence somatic SNVs obtained from the high confidence filter using default parameters were used for further analysis, as per developer recommendations. We further performed annotation and filtering on all SNVs, with full details given previously^13^. In brief, SNVs obtained by MuTect (v1.1.4) and SomaticSniper (v1.0.2) were annotated with associated genes and functions by ANNOVAR (v2015-06-17)^49^ using RefGene, subjected to deny-list filtering to remove known germline contaminants and sequencing artifacts and allow-list filtering through COSMIC (v70)^48^. This was done before downstream subclonal reconstruction. SNVs were further subjected to filtering to remove SNVs not at callable bases (where callable bases are those with ≥ 17x coverage for the tumour and ≥ 10x coverage for the normal).

### Subclonal Reconstruction Pipeline Construction

We define a subclonal reconstruction pipeline as comprised of a SNV detection tool, a CNA detection tool and a subclonal reconstruction algorithm. A pipeline is said to be using or comprising of a tool and/or an algorithm when the tool/algorithm is incorporated as one step of the pipeline.

For single-region reconstruction, the SNV detection tools SomaticSniper (v1.0.2) and MuTect (v1.1.4), the CNA detection tools Battenberg (v2.2.6) and TITAN (v1.11.0), and the subclonal reconstruction algorithms PhyloWGS (v3b75ba9), PyClone (v0.13.0), DPClust (v2.2.5) and SciClone (v1.0.7) were combined in factorial combinations to construct 16 pipelines. Subclonal reconstruction was run on the cohort of 293 tumours with index lesion sequencing for single-region subclonal reconstruction.

For multi-region reconstruction, the SNV detection tools SomaticSniper (v1.0.2) and MuTect (v1.1.4), the CNA detection tools Battenberg (v2.2.6), TITAN (v1.11.0) and FACETS (v0.5.14), and the subclonal reconstruction algorithms PhyloWGS (v3b75ba9), PyClone (v0.13.0) and SciClone (v1.0.7) were combined in factorial combinations to construct 18 pipelines. For the 10 tumours with multi-region sequencing, each individual sequencing sample (total 30, 2-4 samples per tumour) was first subjected to single-region subclonal reconstruction using the 18 pipelines, followed by multi-region subclonal reconstruction using the 18 pipelines where all regions of a tumour were provided as input.

### Subclonal Reconstruction of Tumours using PhyloWGS

We used the cnv-int branch of PhyloWGS (https://github.com/morrislab/phylowgs/tree/cnvint, commit: 3b75ba9c40cfb27ef38013b08f9e089fa4efa0c0)^15^ for the reconstruction of tumour phylogenies, as described previously^13^. Briefly, subclonal CNA segments and cellularity inputs were parsed using the provided parse_cnvs.py script (the parse_cnvs.py was custom augmented to process inputs from FACETS [v0.5.14]) and filtered to remove any segments shorter than 10 kbp. The create_phylowgs_inputs.py script was used to generate PhyloWGS (v3b75ba9) inputs for each sample. All default parameters were used, including limiting the number of SNVs considered to 5,000 for the interest of runtime, to launch reconstructions using evolve.py. Multi-region subclonal reconstruction was performed by providing all regions belonging to the same tumour as input for the reconstruction and the procedure was otherwise identical to the single-region reconstructions.

The best phylogenetic clone tree for each run and the CNAs and SNVs associated with each subclone in that structure were determined by parsing the output JSON files for the tree with the largest log likelihood value. In addition to the best tree structure, the output JSON file was also parsed for all predicted tree structures as ordered by log likelihood values to assess the change in predictions across the 2,500 Markov chain Monte Carlo iterations. Only samples with a total of 2,500 complete MCMC iterations were considered, and samples with poly-tumour or overly complex intermediate clone tree structures that were never the final solution for any sample were excluded.

### Subclonal Reconstruction of Tumours using PyClone

We used PyClone (v0.13.0)^17^ for single- and multi-region mutation clustering. A mutation input file was created for each sample by obtaining the tumour reference and variant read counts for each SNV from input VCFs and annotating them with the clonal major and minor copy numbers for the position from CNA inputs. Since PyClone (v0.13.0) leverages SNVs in clonal CNA regions, all SNVs in subclonal CNA regions were not considered. SNVs in regions without copy number information were also discarded, and the normal copy number was set to 2 for autosomes and 1 for chromosomes × and Y. The mutation input file, along with tumour cellularity as predicted by the subclonal CNA detection tool were used as inputs for the run_analysis_pipeline to launch PyClone (v0.13.0)^17^, using 12345 as the seed for all runs. Notably, since PyClone (v0.13.0) was originally developed for deep sequencing (>100x) data, the developer recommended setting the “density” parameter to “pyclone_binomial” to account for characteristics whole-genome sequencing data. The number of Markov chain Monte Carlo iterations were also set to 100,000, with 1,000 burn-ins. Otherwise default parameters were used. PyClone (v0.13.0) outputted ‘cellular prevalence’ as defined by the authors as ‘the proportion of tumor cells harboring a mutation’ fits the definition of cancer cell fraction for this study, and cellular prevalence as defined in this study was calculated by multiplying the outputted ‘cellular prevalence’ with purity estimates from the respective CNA detection tool. Multi-region reconstructions using PyClone (v0.13.0) were launched by including all mutation input files and tumour cellularities prepared for single-region reconstructions as outlined above for all samples of a tumour as input to run_analysis_pipeline. Cellular prevalence as defined in this study was similarly obtained from ‘cellular prevalence’ as outputted by PyClone (v0.13.0) by individually adjusting for the tumour contents for each sample of the tumour.

### Subclonal Reconstruction of Tumours using DPClust

We used DPClust (v2.2.5)^16^ for single-region subclonal reconstruction. DPClust (v2.2.5) was run using the dpc.R pipeline available *via* the DPClust SMC-HET Docker (https://github.com/Wedge-Oxford/dpclust_smchet_docker, commit a1ef254), using also dpclust3p (v1.0.6). The pipeline was customized to process inputs from SomaticSniper (v.1.0.2) and TITAN (v1.11.0). The inputs for each tumour sample were the VCF file provided by the SNV detection tool, and subclonal copy number, cellularity, ploidy, and purity as predicted by the subclonal CNA detection tool, using 12345 as the seed and otherwise default parameters. The results in the subchallenge1C.txt output file were taken as the mutation clustering solution to obtain the number of subclones predicted by DPClust and their cellular prevalences (v2.2.5)^16^. Results in the subchallenge2A.txt output file were taken to define the mutation composition of each cluster.

### Subclonal Reconstruction of Tumours using SciClone

We used SciClone (v1.0.7)^20^ for single- and multi-region subclonal reconstruction. Input VCFs were used to calculate variant allele frequencies (in percentage) and CNA inputs were used to determine regions with loss of heterozygosity. Only SNVs in clonally copy number neutral (major = 1, minor = 1) regions with no subclonal CNAs were considered by SciClone (v1.0.7) and all samples were run using default parameters. Multi-region reconstructions using SciClone (v1.0.7) were run by including inputs for all samples of a tumour. Mutation clusters defined by SciClone (v1.0.7) were characterized using variant allele frequencies, and their VAFs were multiplied by a factor of 2 to convert to cellular prevalence as defined in this study.

### Post Processing of Subclonal Reconstruction Solutions

Since subclones in PhyloWGS (v3b75ba9) trees are numbered based on cellular prevalence instead of evolutionary relationship, trees were transformed to consistent representations to allow comparison across cohorts following two rules: 1) trees are left-heavy, 2) all nodes at a particular tree depth must have numbers greater than that of nodes at lower tree depths, with the root node (normal cell population) starting at 0. Further, pruning of nodes was performed following the heuristic that each node must have at least 5 SNVs or 5 CNAs and a minimum cellular prevalence of 10%, creating a subclonal diversity lower bound for each tumour^13^. A node was pruned and merged with its sibling if their cellular prevalence difference was ≤ 2% and if both were driven purely by SNVs (had ≤ 5 CNAs). A node was merged with its parent node if their cellular prevalence difference was ≤ 2%. When PhyloWGS (v3b75ba9) produced a poly-tumour solution for the best consensus tree, the algorithm was re-run up to 12 times with different random number generator seeds after which the final poly-tumour solution was accepted and considered to be a reconstruction failure. The seeds were applied in the following order: 12345, 123456, 1234567, 12345678, 123456789, 246810, 493620, 987240, 1974480, 3948960, 7897920 and 15795840. In the event PhyloWGS (v3b75ba9) failed to produce a solution due to reconstruction failures or excessive runtime (> 3 months), the sample was excluded from analysis for that pipeline.

PyClone (v0.13.0), DPClust (v2.2.5) and SciClone (v1.0.7) identified subclonal populations were pruned using similar heuristic as that for PhyloWGS (v3b75ba9). Specifically, for each tumour sample, a mutation cluster was pruned if it had fewer than five supporting SNVs or a cellular prevalence below 10% if it is the clonal cluster or below 2% if it is a subclonal cluster. If there were less than 5 total mutations (SNVs) assigned to clusters in a sample, or if all clusters had cellular prevalence of below 10%, a failed reconstruction was designated to the sample. Otherwise pruned clusters were merged with their nearest neighbor in cellular prevalence, and the weighted mean of cellular prevalence was assigned to the merged node. Moreover, two clusters were merged if they differed in cellular prevalence by ≤ 2%. Finally, mutation clusters were ordered by decreasing cellular prevalence and renumbered accordingly, and the cluster with the highest cellular prevalence was treated as the clonal cluster and its cellular prevalence taken as the cellularity estimated by the pipeline. This was a conservative approach as the detection of multiple primary tumours is challenging from single-sample subclonal reconstruction^13^.

### Union and Intersection of Mutation Detection Tools

We obtained the union and intersection of raw SNVs by SomaticSniper (v1.0.2) and MuTect (v1.1.4) for each tumour sample using vcf-isec of vcftools (v0.1.15). The union and intersection sets of SNVs were then annotated and filtered with the same method as described above before being used in subsequent analysis^13^. For the comparison of characteristics between SNVs detected by MuTect (v1.1.4) and SomaticSniper (v1.0.2), all SNVs detected by each tool across all 293 index lesion samples were pooled to assess their VAFs and trinucleotide contexts. SNVs were grouped as intersect if detected by both tools, or as MuTect-unique or SomaticSniper-unique, both pre- and post-filtering. The effect of filtering was assessed by comparing SNVs retained after filtering (‘SomaticSniper’ and ‘MuTect’) with those removed by it (‘Removed SomaticSniper’ and ‘Removed MuTect’). Trinucleotide context profiles for each group of SNVs were normalized by the expected number of each trinucleotide across the hg19 genome.

We determined the union and intersection of CNAs detected by TITAN (v1.11.0) and Battenberg (v2.2.6), first parsed using parse_cnvs.py script of PhyloWGS (v3b75ba9) for consistent formatting, on a per base-pair basis. The intersection of CNAs, based on genomic coordinates and major and minor copy number, was determined using the GenomicRanges (v1.28.6)^50^ package in R (v3.2.5). Regions with disagreeing copy number were identified using bedtools (v2.27.1)^51^ and bedr (v1.0.6)^52^. A region is defined to have a tool-unique CNA if one tool detected a copy number aberration for the region while the other identified it as copy number neutral (major and minor copy number of 1, both clonally and subclonally). Regions were both algorithms detected different copy number aberrations were classified as disagreements. The union set of CNAs thus contained the intersection of CNAs and CNAs unique to either tool, and regions of disagreement were excluded as there was no natural way to resolve discrepancies. In contrast to TITAN, when a region is determined to have a subclonal aberration, Battenberg (v2.2.6) produces two entries, a clonal and subclonal copy number for each genomic region. These regions were labelled Battenberg-unique for its clear delineation of subclonal CNAs. However, the TITAN (v.1.11.0) copy number aberration result for the region (if any) is used in the union of CNAs to avoid conflicting CNAs in the same region, as one cannot combine clonal Battenberg (v2.2.6) results with TITAN (v1.11.0) aberrations. The union and intersection set of CNAs were further filtered to remove any segments under 10 Kbp.

Four pipeline combinations using PhyloWGS (v3b75ba9) and the intersection and union of SNVs and CNAs were executed on 293 single-region samples. The script create_phylowgs_inputs.py was used to combine intersect and union of SNVs and CNAs as inputs for PhyloWGS (v3b75ba9), where no cellularity estimate was provided as there was no obvious way to derive that for the intersect and union of CNAs. The pipelines were run with otherwise identical procedure as single-region reconstructions with PhyloWGS (v3b75ba9).

### Clonality Classification

We classified the phylogenetic clone trees outputted by PhyloWGS (v3b75ba9) and mutation clustering results outputted by PyClone (v0.13.0), DPClust (v2.2.5) and SciClone (v1.0.7) as monoclonal or polyclonal based on the number of subclones they predicted. Solutions where only one subclone was predicted were termed *monoclonal*. In monoclonal reconstructions, the only subclone detected is then termed the clonal node. Solutions where more than one subclone was predicted were termed *polyclonal*. In polyclonal reconstructions, the subclone with the highest cellular prevalence was deemed clonal, and the rest of the subclones were subclonal. In situations where PhyloWGS (v3b75ba9) outputted phylogenies showing a normal root node with more than one direct child, the clone tree was termed *poly-tumour*, suggestive of multiple independent primary tumours. These were excluded from downstream analysis because the reconstruction of these phylogenies, especially from single sequencing samples, is challenging^13^.

CNA and SNV mutations were classified as clonal or subclonal based on their node assignment in the best PhyloWGS (v3b75ba9) consensus clone tree and PyClone (v0.13.0), DPClust (v2.2.5) and SciClone (v1.0.7) mutation clusters. The mutations that define the clonal node were classified as clonal mutations, while all others were classified as subclonal mutations. The cancer cell fraction (CCF) of mutations was calculated by dividing the cellular prevalence of the node that the mutation belonged to by the predicted cellularity of the tumour sample.

### Analysis of Single Nucleotide Variants

We compared the four pipelines using each subclonal reconstruction algorithm for their inference of clonal and subclonal SNVs. In each pairwise comparison, for each sample we noted the clonal SNV set identified by each algorithm and calculated the Jaccard index between the two sets. The analysis was performed separately for clonal and subclonal SNVs.

### Analysis of Copy Number Aberrations

We further filtered the CNAs identified by PhyloWGS using OncoScan data for samples with the data available, removing the identified CNAs that did not overlap any OncoScan CNAs^13^. For samples without OncoScan data, CNAs outputted by PhyloWGS (v3b75ba9) were filtered to retain only those across genomic locations with recurrence of CNAs in OncoScan-filtered samples, with 10 being the established empirical recurrence threshold^13^. Bins of 1.0 Mbp were created across the genome to characterize the copy number profiles for each sample and were assigned the copy number of overlapping genomic segments, either neutral or mutated. Regions not considered by PhyloWGS (v3b75ba9) due to lack of information were assumed to have the normal copy number of two. Profiles were created separately for clonal and subclonal CNAs. We further used previously identified clonal and subclonal subtypes to cluster samples^13^. Samples that were assigned a subclonal subtype in the SomaticSniper-TITAN pipeline^13^ but had no subclonal populations detected in another pipeline were excluded from subclonal subtype analysis for that pipeline. Samples that had no subclonal populations detected in the SomaticSniper-TITAN pipeline and were therefore never assigned to a subclonal subtype were not considered in any subclonal subtype analysis. For each pipeline, we used the copy number profiles of all samples with available data to generate average subtype-specific clonal and subclonal CNA profiles of localized prostate cancer, with standard deviation.

We compared the CNA profiles identified by the four PhyloWGS-comprising pipelines by assessing the difference in clonal and subclonal CNAs between pipeline pairs. For each sample, a clonal CNA set was generated from pipeline results, where the direction of the CNA is taken into account. For example, if a sample was identified with a clonal gain in genomic bin 1 and a clonal loss in genomic bin 2, it would have the clonal CNA set +1, −2. The Jaccard index of clonal and subclonal CNA sets for each sample were calculated between all pipeline pairs.

We identified CNAs that were differentially altered clonally and subclonally. Using 1.0 Mbp bins across the genome, we aggregated the number of samples with and without a CNA overlapping each 1.0 Mbp stretch, with gains and losses considered separately. Clonal and subclonal CNAs were annotated separately, and only samples with *polyclonal* phylogenies were considered, since they have both clonal and subclonal components. Pearson’s *χ*^2^ test was used with multiple testing correction (FDR ≤ 0.05) to define the bins that were significantly enriched for clonal or subclonal CNAs that were gain or loss. CNAs in these bins were thus considered significantly differentially altered, with a predisposition to occur clonally or subclonally as a gain or a loss. Genes affected by differentially altered CNAs were annotated using RefSeq, and the lists of genes considered to have CNA biases by the four pipelines were compared for overlap.

We performed pathway enrichment analysis on the genes that were identified by all four PhyloWGS-comprising pipelines as affected by CNAs biased clonally or subclonally. Using all default parameters of gprofiler2 (v0.1.9) in R (v3.5.3)^53^, statistically significant pathways from Gene Ontology (Biological Process, Molecular Function and Cellular Component), KEGG and Reactome were computed, with no electronic GO annotations. We discarded pathways that involved > 350 or < 5 genes. Cytoscape (v3.4.0) was used to visualize significant pathways^54^. Since all genes identified as significantly differentially altered were biased to be altered clonally, we defined these pathways as differentially altered clonally.

### Driver Mutation Analysis

We gathered a list of known prostate cancer driver genes based on previous large sequencing studies^13,23^. The known CNA-affected driver genes considered were *MYC*, *TP53*, *NKX3-1*, *RB1*, *CDKN1B*, *CHD1*, *PTEN* and *CDH1*. The known SNV-affected driver genes considered were *ATM*, *MED12*, *FOXA1*, *SPOP* and *TP53*. PhyloWGS-comprising pipelines identified CNAs overlapping CNA-affected driver genes and SNVs that occurred in SNV-affected driver genes. These were defined to be driver CNAs and driver SNVs, respectively. A sample was considered to have a consensus driver mutation, CNA or SNV, if the mutation was identified with the same clonality by all four PhyloWGS-comprising pipelines.

Driver SNVs and CNAs of each sample were categorized by the number of PhyloWGS-comprising pipelines they were identified in. Since four PhyloWGS-comprising pipelines were used, in each sample driver SNVs and CNAs could be identified in all four pipelines, three pipelines, two pipelines or one pipeline. Proportions of each category were calculated by dividing the number of samples in that category by the sum of samples assigned to all categories for the driver SNV or CNA. The analysis was done separately for clonal and subclonal mutations, such that the category of the driver SNVs or CNAs in a sample was defined by the most frequent identification of the clonality. For example, if a driver SNV in a sample was identified as clonal by two pipelines, subclonal by one pipeline and wildtype by the last pipeline, it would be counted in both category two for the clonal analysis and in category one for the subclonal analysis.

### Biomarker Survival Analysis

We assessed the utility of clonality (monoclonal *vs.* polyclonal) as a biomarker in all sixteen pipelines used for single region subclonal reconstruction of 293 samples. Tumours were grouped by clonality and the two groups were compared using a log-rank test for differences in outcome. Tumours were also grouped by integrating the previously defined multi-modal biomarker^23^ (groups patients into low risk and high-risk) and clonality, creating unified groups (unified-low: monoclonal low-risk, unified-intermediate: monoclonal high-risk or polyclonal low-risk, unified-high: polyclonal high-risk)^13^ that were compared using a log-rank test. Primary outcome as time to biochemical recurrence (BCR) was described in detail previously^13^. In brief, BCR was defined as PSA rise of ≥ 2.0 ng/mL above the nadir for radiotherapy patients and two-consecutive post-surgery PSA measurements > 0.2 ng/mL (backdated to the date of first increase in PSA) for surgery patients. If a surgery patient had a post-operative PSA ≥ 0.2 ng/mL this was considered primary treatment failure. After salvage radiation therapy, if PSA continued to rise, BCR was backdated to the first PSA measurement > 0.2 ng/mL, but if not then then this was not considered a BCR. Salvage therapy (hormone therapy or chemotherapy) was considered a BCR.

### Comparing Reconstruction using Single and Multiple Regions

For each of the 10 tumours with multi-region sequencing, we compared the subclonal reconstruction solutions from each single region with the solutions obtained from subclonal reconstruction using all tumour regions. In addition to number of subclones predicted, we compared SNV and CNA clonality predictions between single- and multi-region reconstructions. For all SNVs that were identified in a single-region or its corresponding multi-region reconstruction, we calculated the proportion of SNVs in each of the following categories:

1. Multi- and single-region match: same SNV clonality in single- and multi-region.
2. Clonal in multi-region: SNV identified in both single- and multi-region reconstructions, but SNV is clonal in multi-region and subclonal in single-region.
3. Subclonal in multi-region: SNV identified in both single- and multi-region reconstructions, but SNV is subclonal in multi-region and clonal in single-region.
4. Unique in single-region: SNV only present in single-region reconstruction.
5. Unique in multi-region: SNV only present in multi-region reconstruction.

Similarly, all CNAs that were identified in a single-region reconstruction or its matching multi-region reconstruction were assigned to categories defined in a similar fashion. Additional separation was added for CNAs to distinguish between clonal and subclonal predictions.

### Data Visualization and Reporting

Data was visualized using the R statistical environment (v3.2.5 or v3.5.3), and performed using the lattice (v0.20-34), latticeExtra (v0.6-28), VennDiagram (v1.6.21)^55^ and BPG (v5.3.4)^56^ packages. All boxplots show the median (center line), upper and lower quartiles (box limits), and whiskers extend to the minimum and maximum values within 1.5 times the interquartile range (Tukey boxplots). Figures were compiled in Inkscape (v0.91). Standard deviation of the sample mean was reported for point estimates. All statistical tests were two-sided.

## List of abbreviations

CCF: Cancer Cell Fraction
CNAs: Copy Number Aberrations
II: Intersect of SNVs and Intersect of CNAs
IU: Intersect of SNVs and Union of CNAs
MB: MuTect-Battenberg
MCMC: Markov chain Monte Carlo
MF: MuTect-FACETS
MT: MuTect-TITAN
SB: SomaticSniper-Battenberg
SD: standard deviation
SF: SomaticSniper-FACETS
SNVs: Single Nucleotide Variants
ST: SomaticSniper-TITAN
UI: Union of SNVs and Intersect of CNAs
UU: Union of SNVs and Union of CNAs
VAF: Variant Allele Frequency
WGS: Whole-genome Sequencing

## Data Availability

Published data analyzed in this study, publicly available with appropriate Data Access Compliance Office authorization, include:

WGS Data – Baca *et al.*, 2013: dbGaP, phs000447.v1.p1 [https://www.ncbi.nlm.nih.gov/projects/gap/cgi-bin/study.cgi?study_id=phs000447.v1.p1]^35^

WGS Data – Berger *et al.*, 2011: dbGaP, phs000330.v1.p1 [https://www.ncbi.nlm.nih.gov/projects/gap/cgi-bin/study.cgi?study_id=phs000330.v1.p1]^36^

WGS Data – CPC-GENE Espiritu *et al.*, 2018: EGA, EGAD00001001094

[https://www.ebi.ac.uk/ega/datasets/EGAD00001001094]^13^

WGS Data – CPC-GENE Fraser *et al.*, 2017: EGA, EGAD00001001094

[https://www.ebi.ac.uk/ega/datasets/EGAD00001001094]^23^

WGS Data – CPC-GENE Taylor *et al.*, 2017: EGA, EGAD00001002739

[https://www.ebi.ac.uk/ega/datasets/EGAD00001002739]^24^

WGS Data – The Cancer Genome Atlas Research Network, 2015: https://portal.gdc.cancer.gov/projects/TCGA-PRAD^37^

WGS Data – Weischenfeldt *et al.*, 2013: EGA, EGAS00001000400 [https://www.ebi.ac.uk/ega/studies/EGAS00001000400]^38^

Data supporting the conclusions of this article is included within it and its additional files, and at: ICGC Data Portal under the project PRAD-CA [https://dcc.icgc.org/projects/PRAD-CA].

Source Data for Figures 4 is provided at: ICGC Data Portal under the project PRAD-CA [https://dcc.icgc.org/projects/PRAD-CA]. Data is available with appropriate ICGC Data Access Compliance Office approval.

Source data for Supplementary Figure 1 is provided in Supplementary Information.

Source data for Figures 2, 3, 6 and Supplementary Figures 2, 3A, 4C-F, 7, 8B, 9, 10, 11, 12, 13 are provided in Supplementary Data.

Source data for Figures 5, 7 and Supplementary Figures 3B-E, 4A-B, 5, 6, 8AC, 14 are provided in Source Data.

## Code Availability

No custom algorithms or software were developed or utilized in this study. Custom data analysis & data visualization code is available upon request.

## Acknowledgements

The authors thank Dr. Reimand (University of Toronto) for technical support. We also thank all members of the Boutros and Kislinger labs for helpful suggestions and technical support.

This work was supported by Prostate Cancer Canada and is proudly funded by the Movember Foundation - Grant #RS2014-01 to PCB. PCB was supported by a Terry Fox Research Institute and CIHR New Investigator Awards. This work was supported by NSERC Discovery Grants to QDM and PCB. This research is funded by the Canadian Cancer Society (grant #705649) and a Project Grant from CIHR. This work was funded by the Government of Canada through Genome Canada (OGI-125). VB, LYL and AS were supported by Fellowships from the Canadian Institutes of Health Research. The results described here are in part based upon data generated by the TCGA Research Network: http://cancergenome.nih.gov/. This work was supported by the NIH/NCI under awards P30CA016042, 1U01CA214194-01, 1U24CA248265-01 and 1R01CA244729-01.

## Author Contributions

Initiated the Project: VB, LYL, QDM, PCB

Data Analyses: LYL, VB, SMGE

Data Visualization: LYL, VB, AS

Supervised Research: QDM, TK, PCB

Wrote the First Draft of the Manuscript: VB, LYL, PCB

Approved the Manuscript: All Authors

## Ethics Compliance

All tumour samples in this study were obtained with patient informed consent, with approvals by the University Health Network Institutional Research Ethics Board, the Centre Hospitalier Universitaire de Québec Institutional Research Ethics Board and the University of California Los Angeles Institutional Research Ethics Board, and following ICGC guidelines.

