## Supplementary Information for "Quantifying the Influence of Mutation Detection on Tumour Subclonal Reconstruction"

### Supplementary Figure 01

A

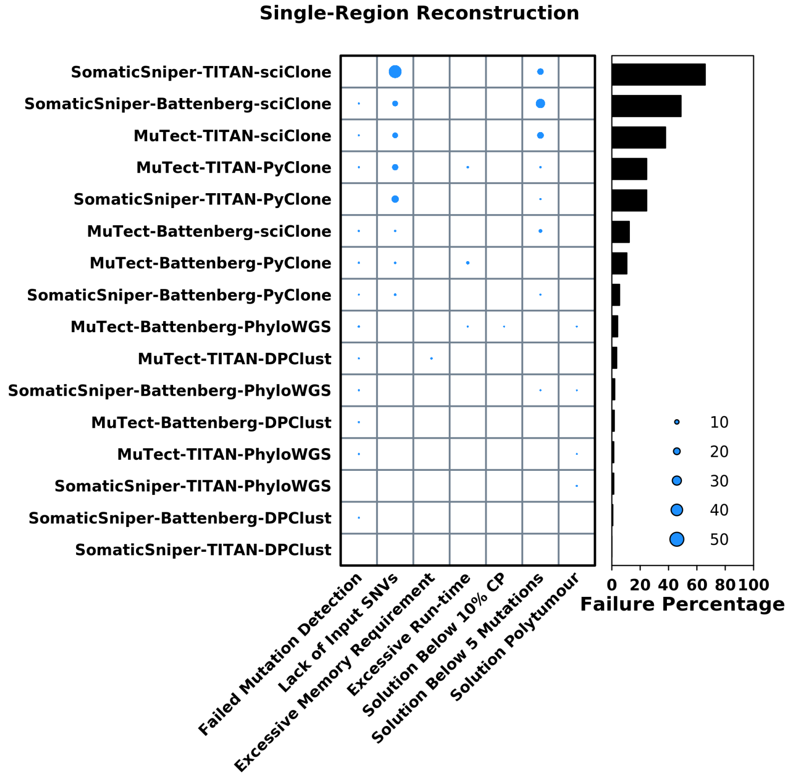


B

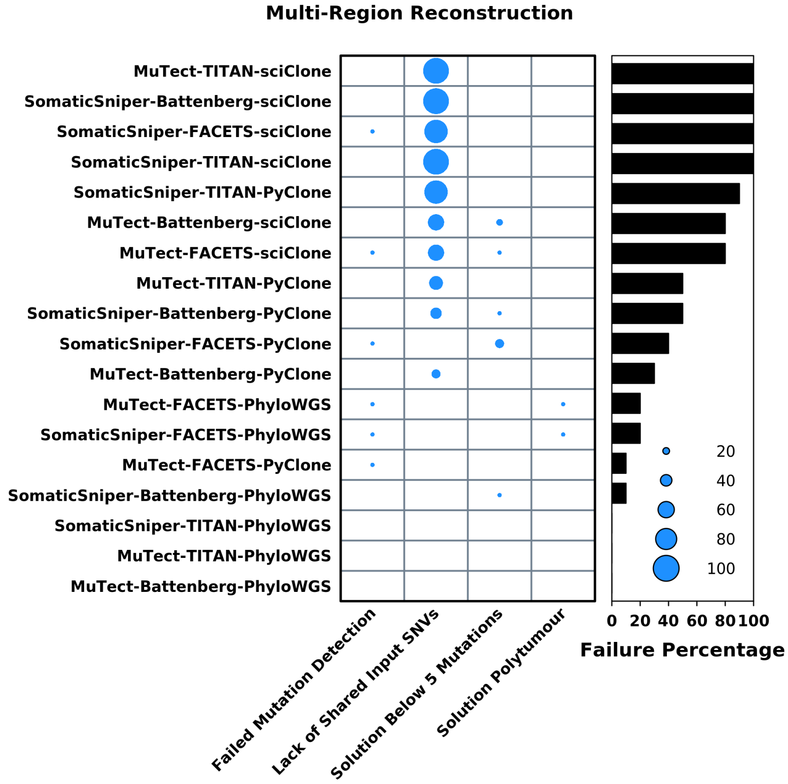


#### Supplementary Figure 1 – Reconstruction Failures

Percentage of 293 samples that failed single-region reconstruction (n=293 biologically independent samples) **A**) and percentage of 10 samples that failed multi-region reconstruction (n=10) **B**). Information is represented by pipeline and reason of failure, where the size of the dot corresponds to percentage of samples.

#### Supplementary Figure 02

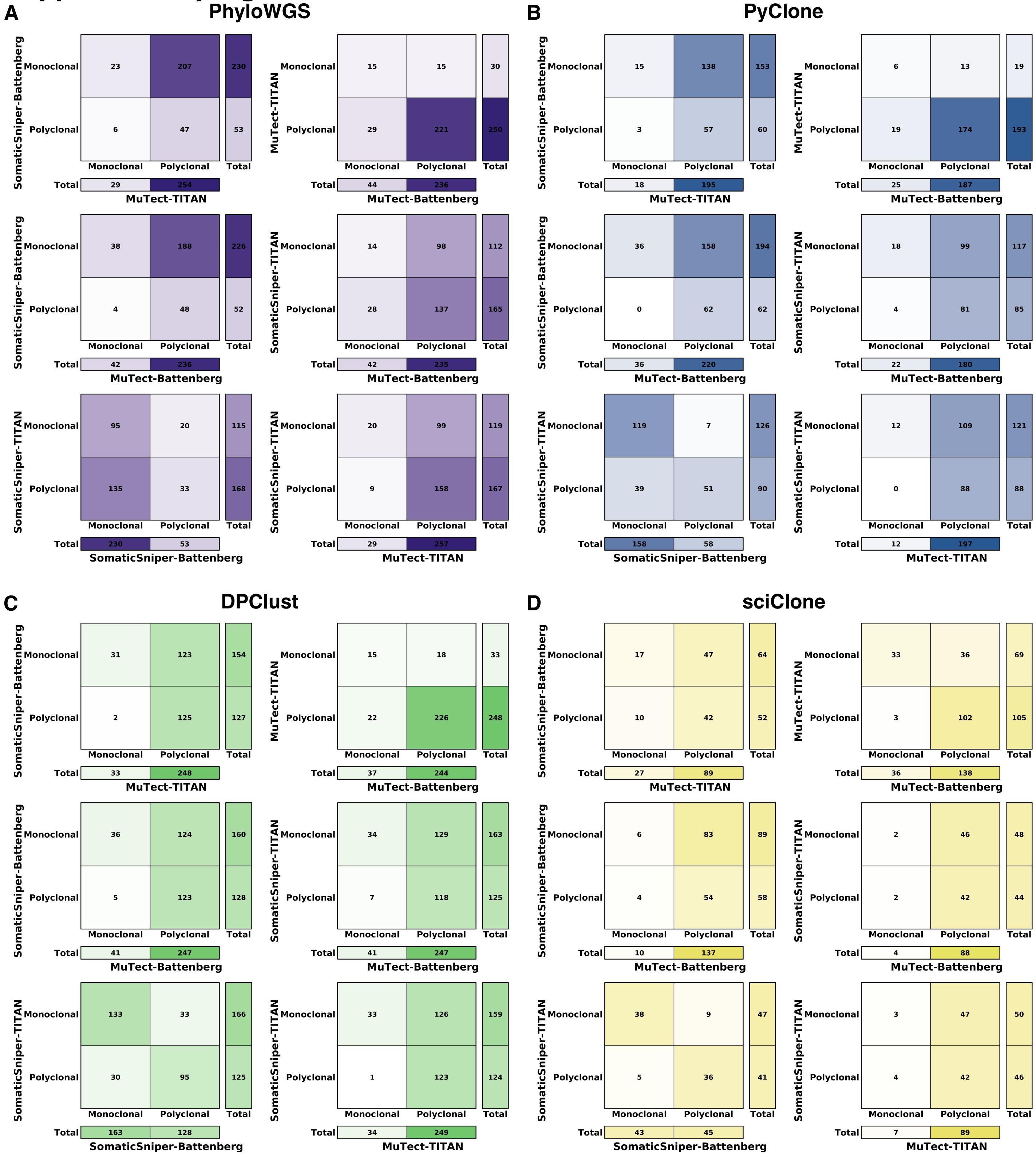


**Supplementary Figure 2 – Clonality between Mutation Detection Tool Combinations**

Comparison of clonality (monoclonal or polyclonal) for shared successfully executed samples between pipeline pairs using the same subclonal reconstruction algorithms PhyloWGS **A**), PyClone **B**), DPClust **C**), SciClone **D**). Darkness of background corresponds to number of samples.

### Supplementary Figure 03

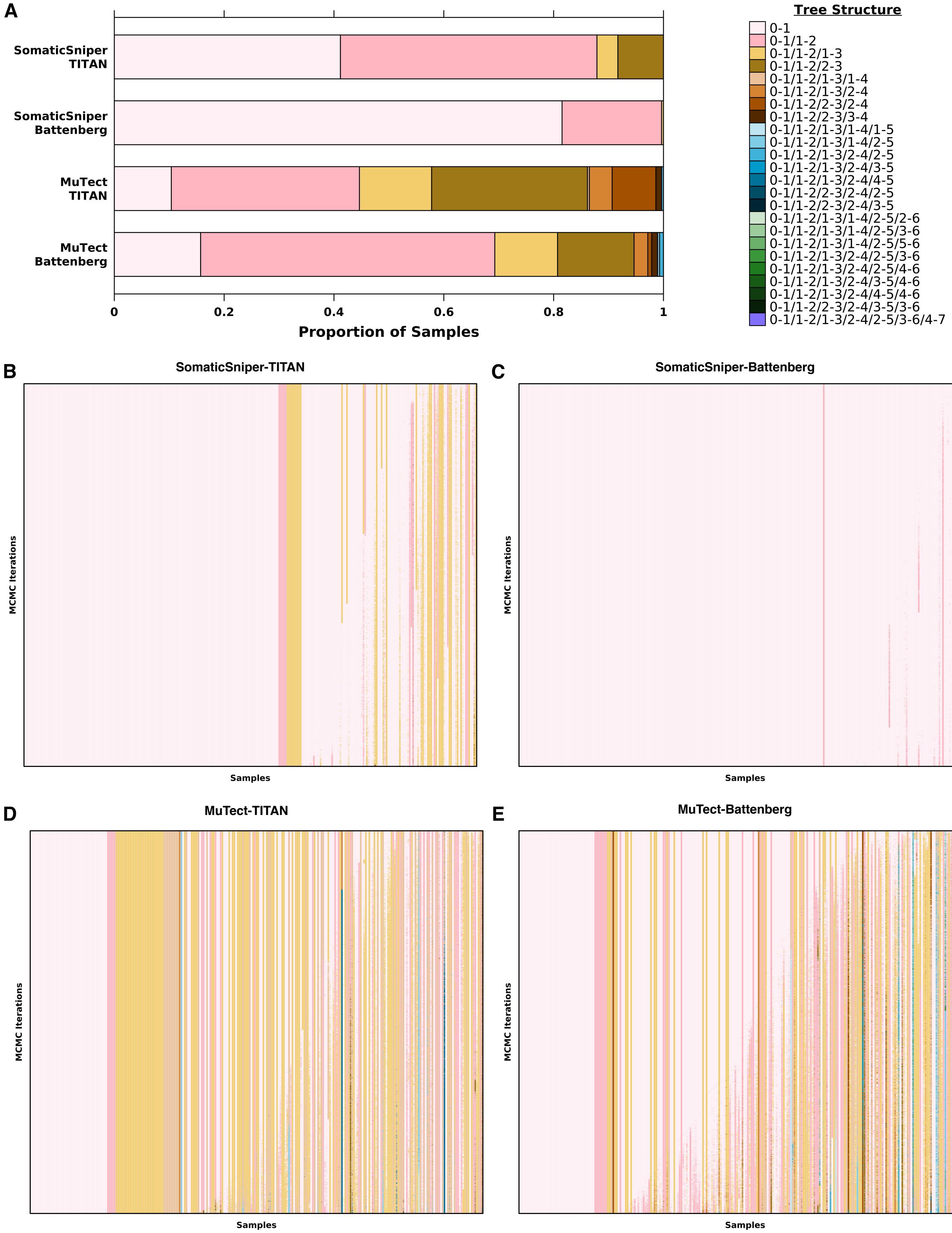

#### Supplementary Figure 3 – Clone Tree Structures between Mutation Detection Tool Combinations

**A**) Proportion of samples with each clone tree structure as predicted by pipelines using PhyloWGS, with each stacked bar representing a pipeline. Tree structures are indicated by color of the stacked bar. SomaticSniper-TITAN: n=289 biologically independent samples; SomaticSniper-Battenberg: n=287; MuTect-TITAN: n=289; MuTect-Battenberg: n=280. Samples with polyclonal solutions SomaticSniper-TITAN: n=170; SomaticSniper-Battenberg: n=53; MuTect-TITAN: n=259; MuTect-Battenberg: n=236. All clone tree structures estimated across 2500 Markov chain Monte Carlo (MCMC) iterations of PhyloWGS for each sample are shown in **B-E**). Each column represents all clone tree structures predicted by 2500 iterations of MCMC for a sample, ordered top to bottom by decreasing log likelihood. Samples are ordered in increasing order by the first ordered iteration where a different clone tree structure was predicted, then by the complexity of the tree structure. Only samples with 2500 complete iterations of MCMC and harboring only the clone tree structures listed are considered. SomaticSniper-TITAN: n=274; SomaticSniper-Battenberg: n=262; MuTect-TITAN: n=265; MuTect-Battenberg: n=252. Number of samples with alternative phylogeny SomaticSniper-TITAN: n=106; SomaticSniper-Battenberg: n=85; MuTect-TITAN: n=176; MuTect-Battenberg: n=199. Number of samples with solutions only differing in clone tree structures SomaticSniper-TITAN: n=8; SomaticSniper-Battenberg: n=8; MuTect-TITAN: n=68; MuTect-Battenberg: n=95. Source data for presented and not presented samples are all provided as a Source Data file.

#### Supplementary Figure 04

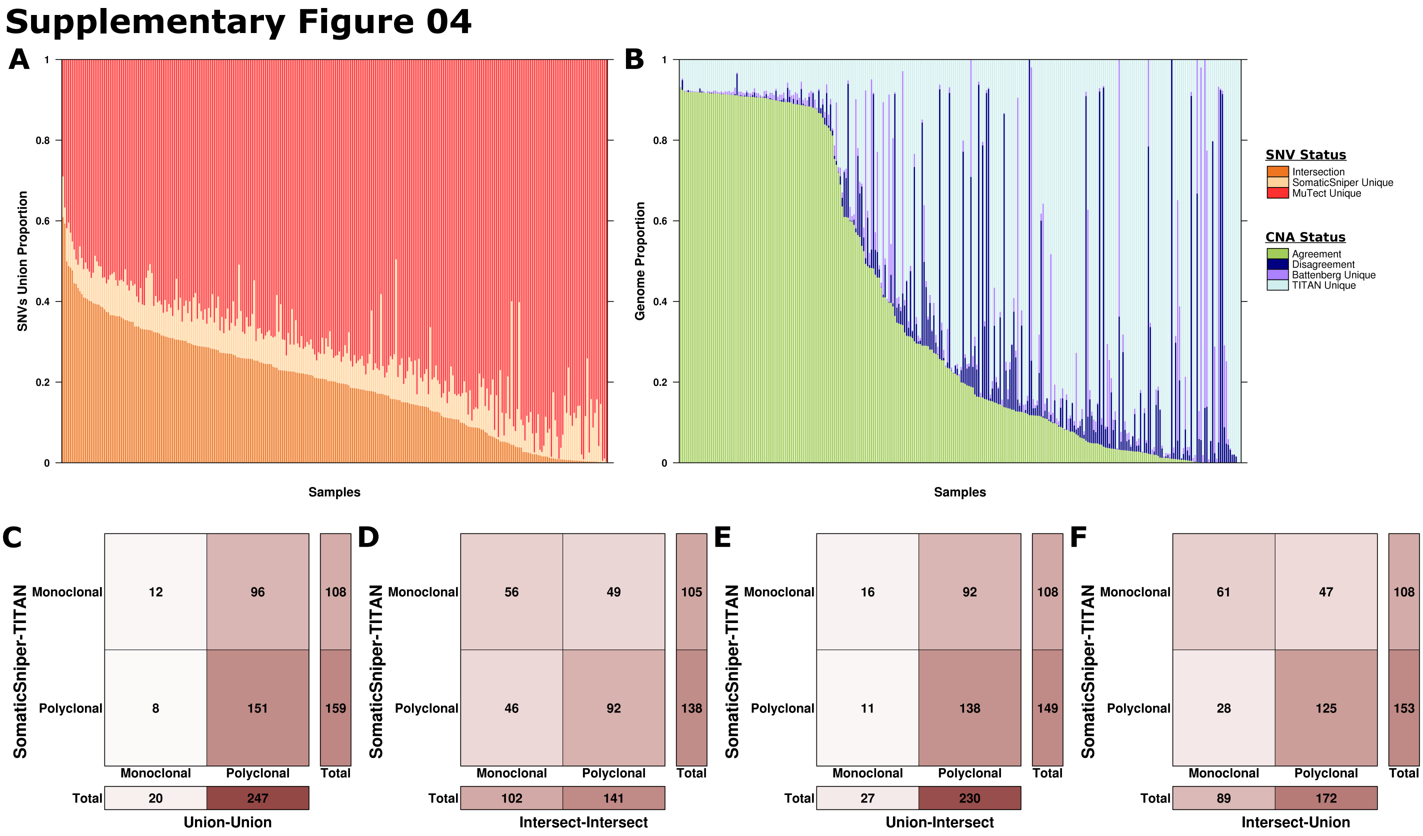


#### Supplementary Figure 4 –Union and Intersection of SNVs and CNAs

**A**) Proportion of unique and intersecting SNVs detected by MuTect and SomaticSniper (n=288 biologically independent samples). **B**) Proportion of the genome covered by unique and intersecting CNAs detected by TITAN and Battenberg (n=288). Each stacked bar represents a sample and the color of the stack bar represents the status of the mutation. **C-F**) Comparing predictions of clonality by the SomaticSniper-TITAN-PhyloWGS pipeline to PhyloWGS-comprising pipelines that had inputs based on the union or intersection of SNVs and CNAs. Darkness of the background corresponds to the number of samples. Source data are provided as a Source Data file.

#### Supplementary Figure 05

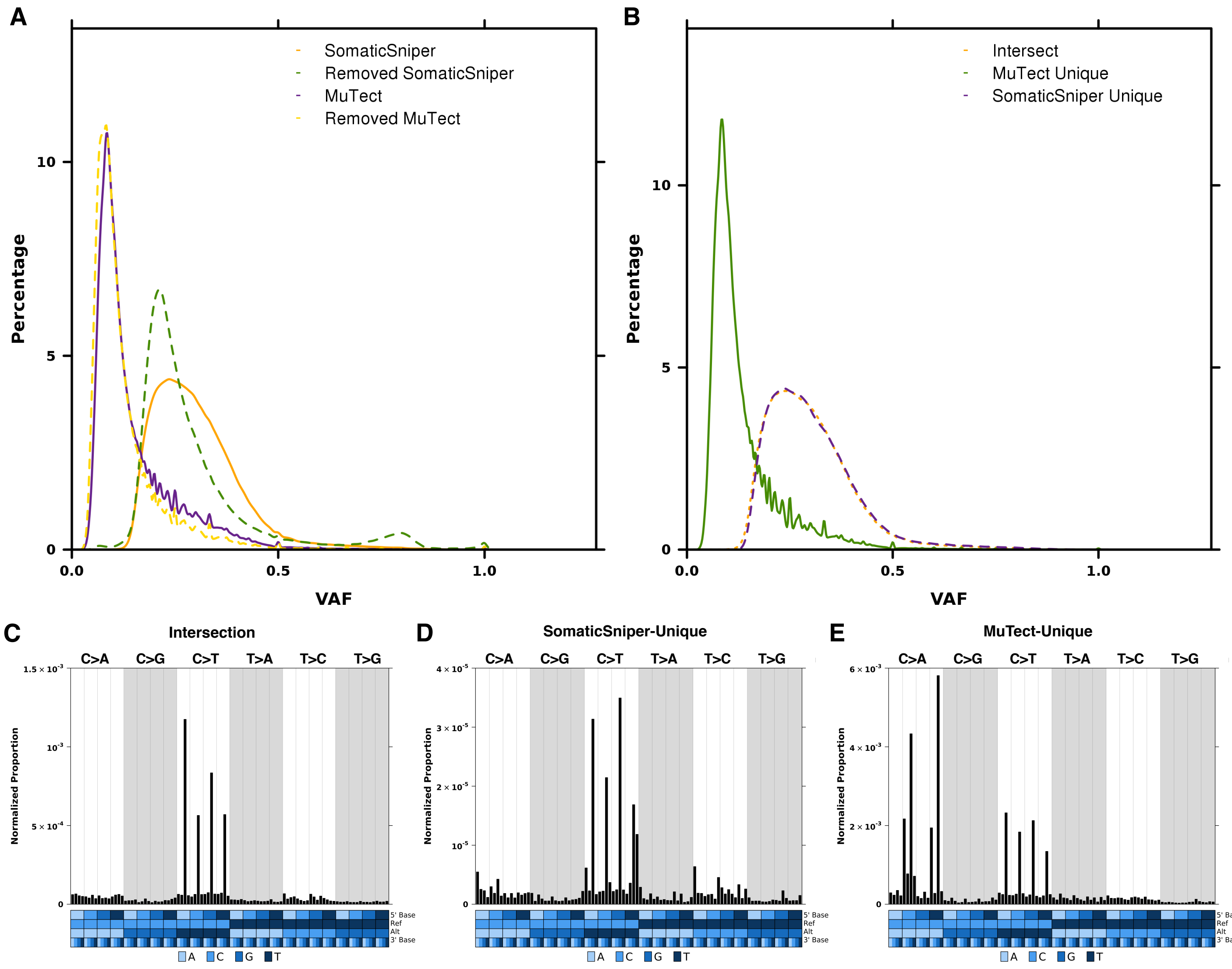


**Supplementary Figure 5 – Effect of SNV Filtering**

**A**) Density plots of variant allele frequencies (VAFs) for SNVs across all samples that were detected by SomaticSniper and MuTect post-filtering (SomaticSniper - long dash yellow line, MuTect - solid purple line), and SNVs that were removed by custom deny-list and allow-list filtering (Removed SomaticSniper - long dash green line, Removed MuTect - solid orange line). **B**) Considering only post-filtering SNVs, density plots of variant allele frequencies (VAFs) for SNVs across all samples that were detected by both SomaticSniper and MuTect (Intersect - short dash orange line), MuTect only (MuTect Unique - solid green line) and SomaticSniper only (SomaticSniper Unique - long dash purple line). **C**) Trinucleotide profile of post-filter SNVs that were detected by both SomaticSniper and MuTect, where the number of SNVs was normalized by the expected number of each trinucleotide context across the hg19 genome. Trinucleotide profiles for post-filter SomaticSniper-unique SNVs **D**) MuTect-unique SNVs **E**). Colors in the covariate bar indicate the 5', reference, alternative and 3' nucleotides in each trinucleotide context. Ref, reference nucleotide; Alt, alternative nucleotide of variant. Number of post-filtering SNVs across all samples in SomaticSniper: n=332,961 independent observations; Removed SomaticSniper: n=319,517; MuTect: n=2,246,971; Removed

MuTect: n=1,008,430; Intersect: n=185,044; SomaticSniper Unique: n=147,917; MuTect Unique: n=2,061,927. Source data are provided as a Source Data file.

#### Supplementary Figure 06

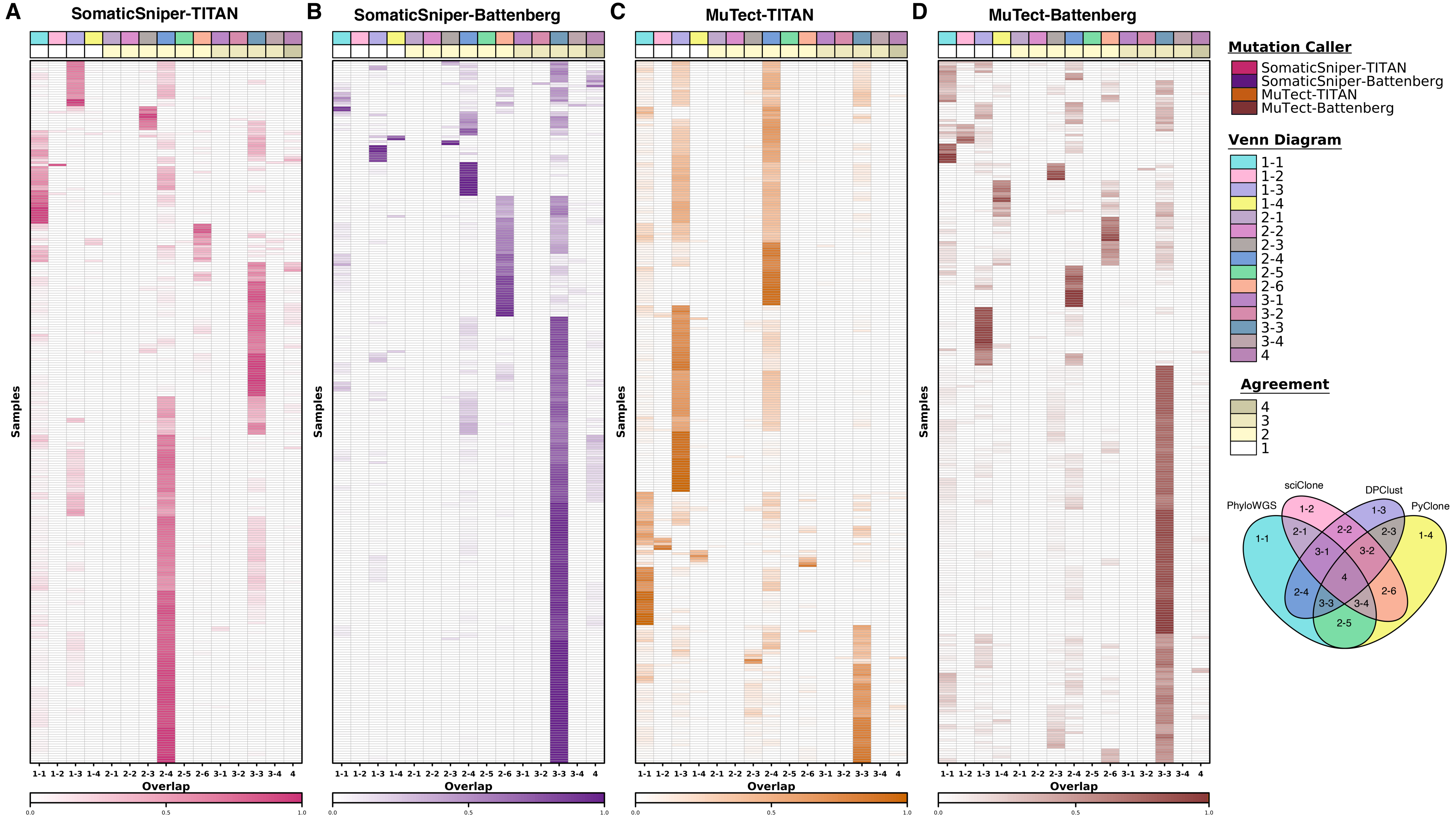

#### Supplementary Figure 6 – Clonal SNVs by Subclonal Reconstruction Algorithm

Proportional overlap between clonal SNVs identified across pipelines using the same mutation detection tool combinations but different subclonal reconstruction algorithms **A-D**). Each row indicates clonal SNV overlap proportions for a sample. Venn diagram indicates the overlap groups of the SNVs, and the first number in the group name indicates the number of subclonal reconstruction algorithms in the overlap. Overlap groups are also represented by colors in the covariate bar, with both legend and Venn diagram reference provided. Covariate bar also indicates, by color, the number of subclonal reconstruction algorithms in agreement for each overlap group. Heatmap colors also correspond with mutation detection tool combinations, with intensity corresponding to proportion of clonal SNVs. SomaticSniper-TITAN: n=293 biologically independent samples; SomaticSniper-Battenberg: n=291; MuTect-TITAN: n=290; MuTect-Battenberg: n=288. Source data are provided as a Source Data file.

### Supplementary Figure 07

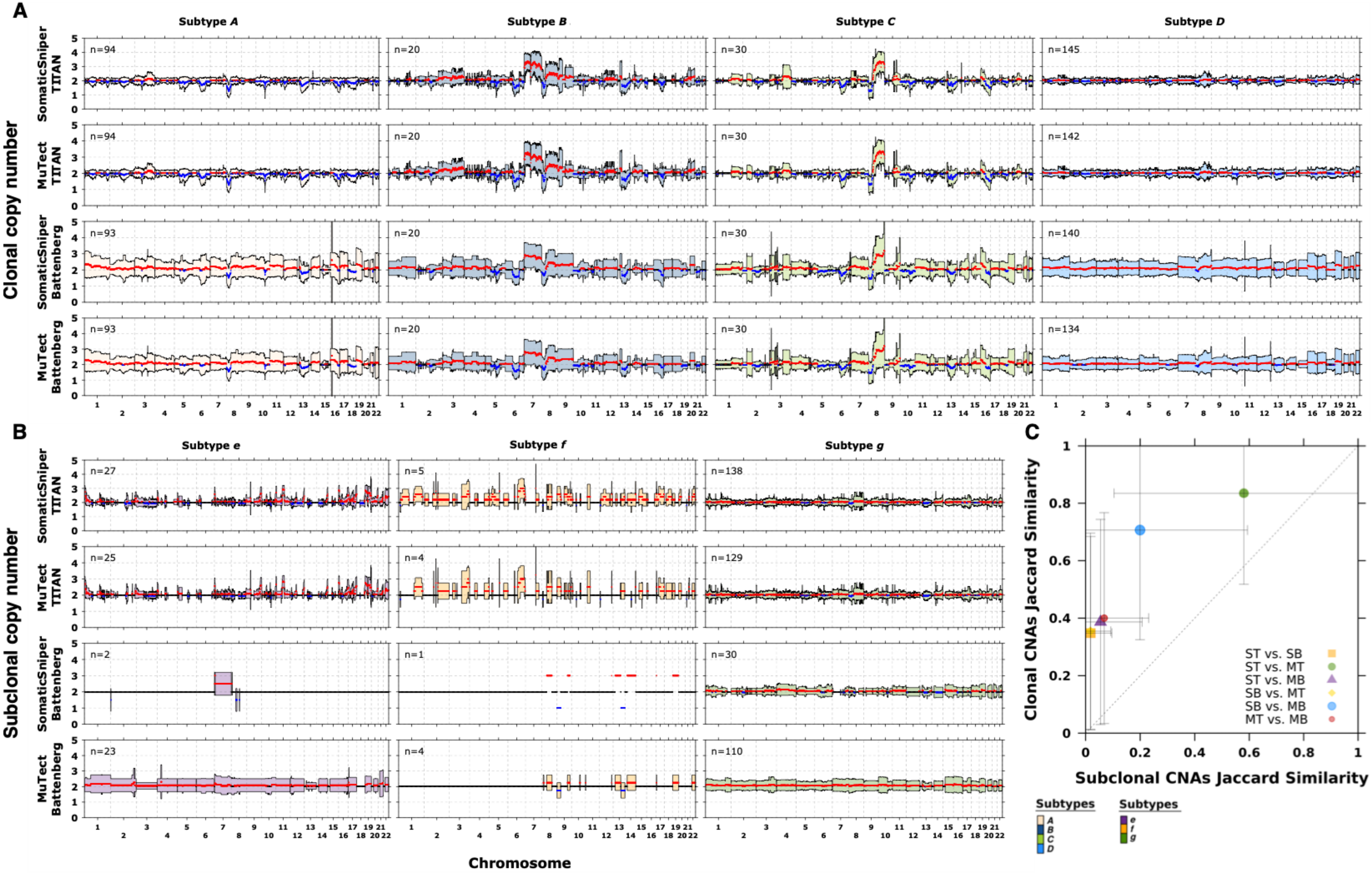

#### Supplementary Figure 7 – Clonal and Subclonal CNAs

Average clonal and subclonal CNA profiles based on four subclonal reconstruction pipelines using PhyloWGS, organized into subtypes determined on the SomaticSniper-TITAN pipeline **A-B**). Chromosomes are shown along the *x*-axis and copy number on the *y*-axis, while each horizontal panel represents the mean clonal or subclonal copy numbers in the CNA profile of one pipeline, based on previously identified clonal and subclonal subtypes. Clonal CNA subtype average profiles are shown in **A**) and subclonal CNA subtypes are shown in **B**). Red indicates that the mean copy number is above the neutral copy number of two while blue indicates that the mean is below two. The shaded regions indicate one standard deviation and is colored to delineate each CNA clonal and subclonal subtype. Sample size is labeled within each panel. **C**) Each marker (delineated by shape and color) represents the comparison between a pair of pipelines and the Jaccard index of 1.0 Mbp genomic bins with clonal and subclonal CNAs. Mean agreement across samples is shown with error bars indicating one standard deviation. Dashed diagonal line represents the $y = x$ line. ST, SomaticSniper-TITAN; MT, MuTect-TITAN; SB, SomaticSniper-Battenberg; MB, MuTect-Battenberg. PhyloWGS ST vs. SB: n=290 independent observations; PhyloWGS ST vs. MT: n=290; PhyloWGS ST vs. MB: n=284; PhyloWGS SB vs. MT: n=287; PhyloWGS SB vs. MB: n=284; PhyloWGS MT vs. MB: n=284.

#### Supplementary Figure 08

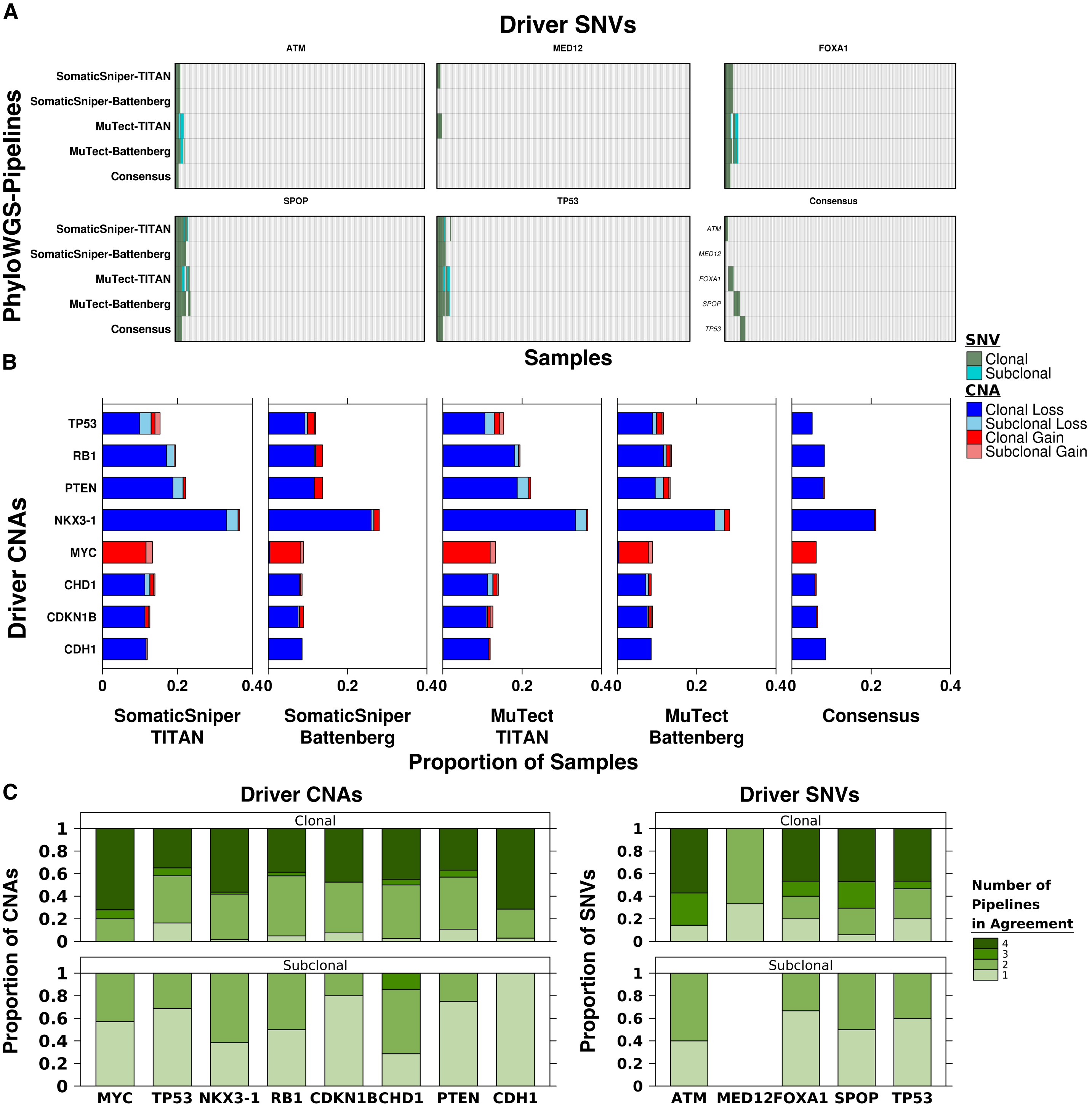


#### Supplementary Figure 8 – Driver CNAs and SNVs

Clonal and subclonal identification of localized prostate cancer driver mutations (single nucleotide variants – SNVs; copy number aberrations - CNAs) based on four pipelines using PhyloWGS and a consensus. **A**) Sub-panels show data for five genes known to be recurrently altered by SNVs in localized prostate cancer. Each column within each sub-panel represent one sample and clonal (green) and subclonal (blue) SNVs detected by each pipeline are indicated by color. **B**) Sub-panels show data for eight genes known to be recurrently altered by CNAs in localized prostate cancer. Clonal and subclonal CNAs identified by each pipeline are shown along with a consensus. Each stacked bar represents a gene and color of the stack bar indicates the type of CNA. **C**) Proportion of samples where each number of pipelines agree on the clonality of driver mutations, assessed separately for clonal and subclonal driver mutations whenever the clonality was predicted by at least one pipeline. Each stacked bar represents information for a gene and darkness of the stacked bar corresponds to the number of pipelines in agreement. SomaticSniper-TITAN: n=289 biologically independent samples; SomaticSniper-Battenberg: n=287; MuTect-TITAN: n=289; MuTect-Battenberg: n=281. Number of SNV driver mutations SomaticSniper-TITAN: n=46 independent observations; SomaticSniper-Battenberg: n=39; MuTect-TITAN: n=59; MuTect-Battenberg: n=56. Number of CNA driver mutations SomaticSniper-TITAN: n=426 independent observations; SomaticSniper-Battenberg: n=299; MuTect-TITAN: n=426; MuTect-Battenberg: n=298. Source data are provided as a Source Data file.

#### Supplementary Figure 09

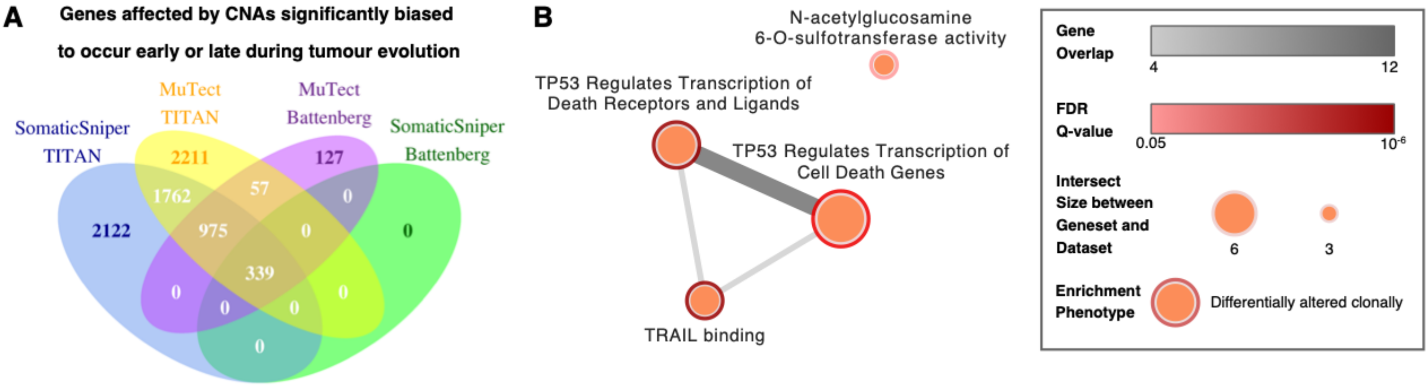


##### Supplementary Figure 9 – Differentially Altered CNAs

**A**) Venn diagram of genes with CNAs that were significantly biased to be altered clonally (*i.e.,* early during tumour evolution) or subclonally (*i.e.,* late during tumour evolution) based on reconstructions from four pipelines using PhyloWGS. Numbers in overlapping areas correspond to the genes with biased timing with the same directionality (*i.e.,* consistent bias for the same gene towards early or late alterations). All 339 genes with biased timing based on all four pipelines were affected by alterations early during tumour evolution. Colors are used to help delineate between Venn's. **B**) Pathway enrichment map based on the 339 genes that were consistently differentially altered clonally. Legend indicates the number of gene overlaps between pathway gene sets (darkness of edges between vertices), false-discovery-rate (FDR) adjusted Q-values for each enriched pathway (darkness of the rims of vertices), and the number of shared genes between gene sets and the dataset (size of vertices).

### Supplementary Figure 10

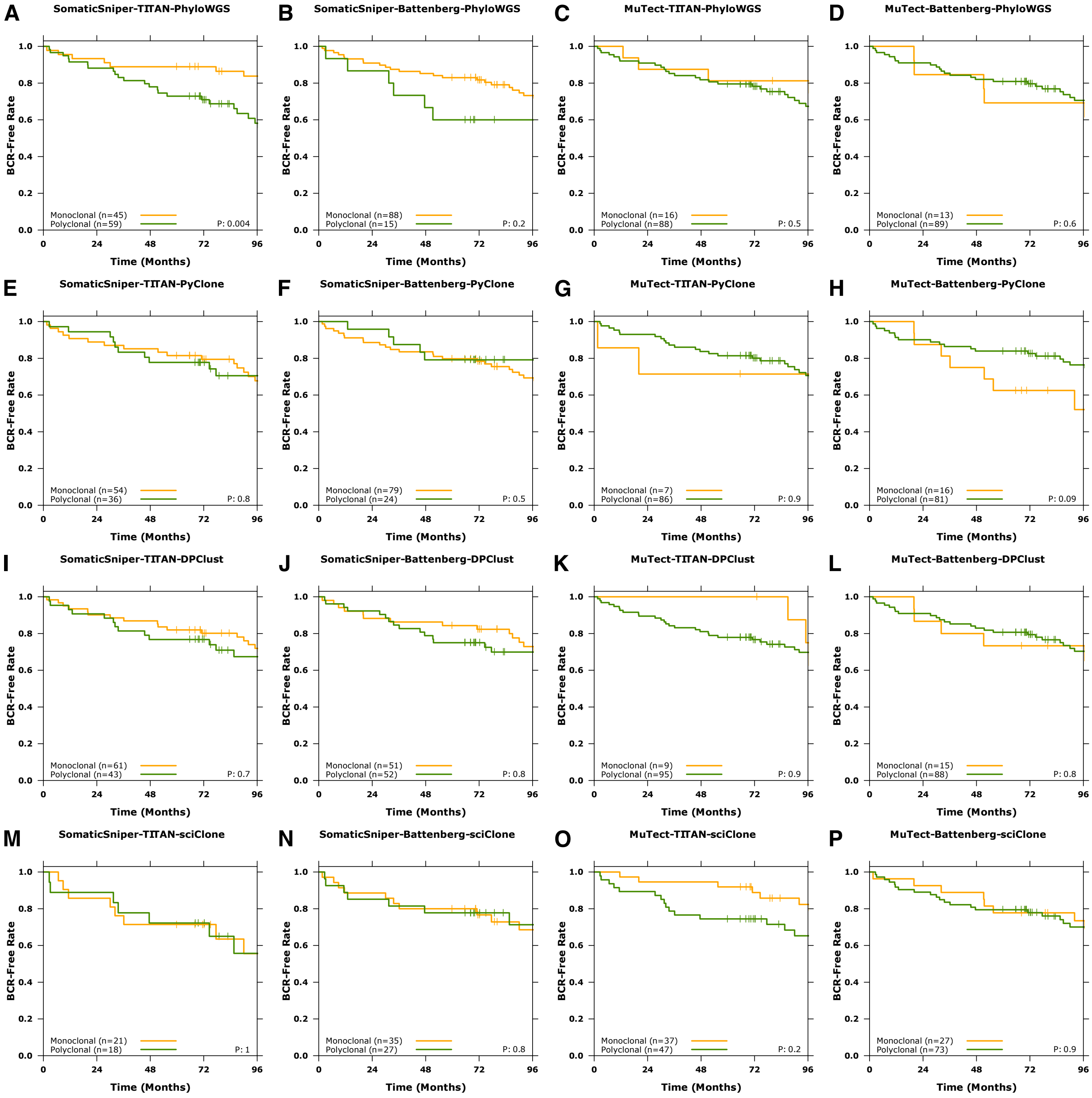


#### Supplementary Figure 10 – Clonality Biomarker

Patients stratified using clonality (monoclonal or polyclonal) as a biomarker across all sixteen subclonal reconstruction pipelines for single-region reconstruction and groups were tested for associations with biochemical recurrence (BCR) **A-P**). Univariate modelling was carried out using log-rank tests, no covariates were included. Color of line in the KM-plots indicate patient groups, monoclonal - yellow, polyclonal - green. Sample size of each group is given as labeled.

#### Supplementary Figure 11

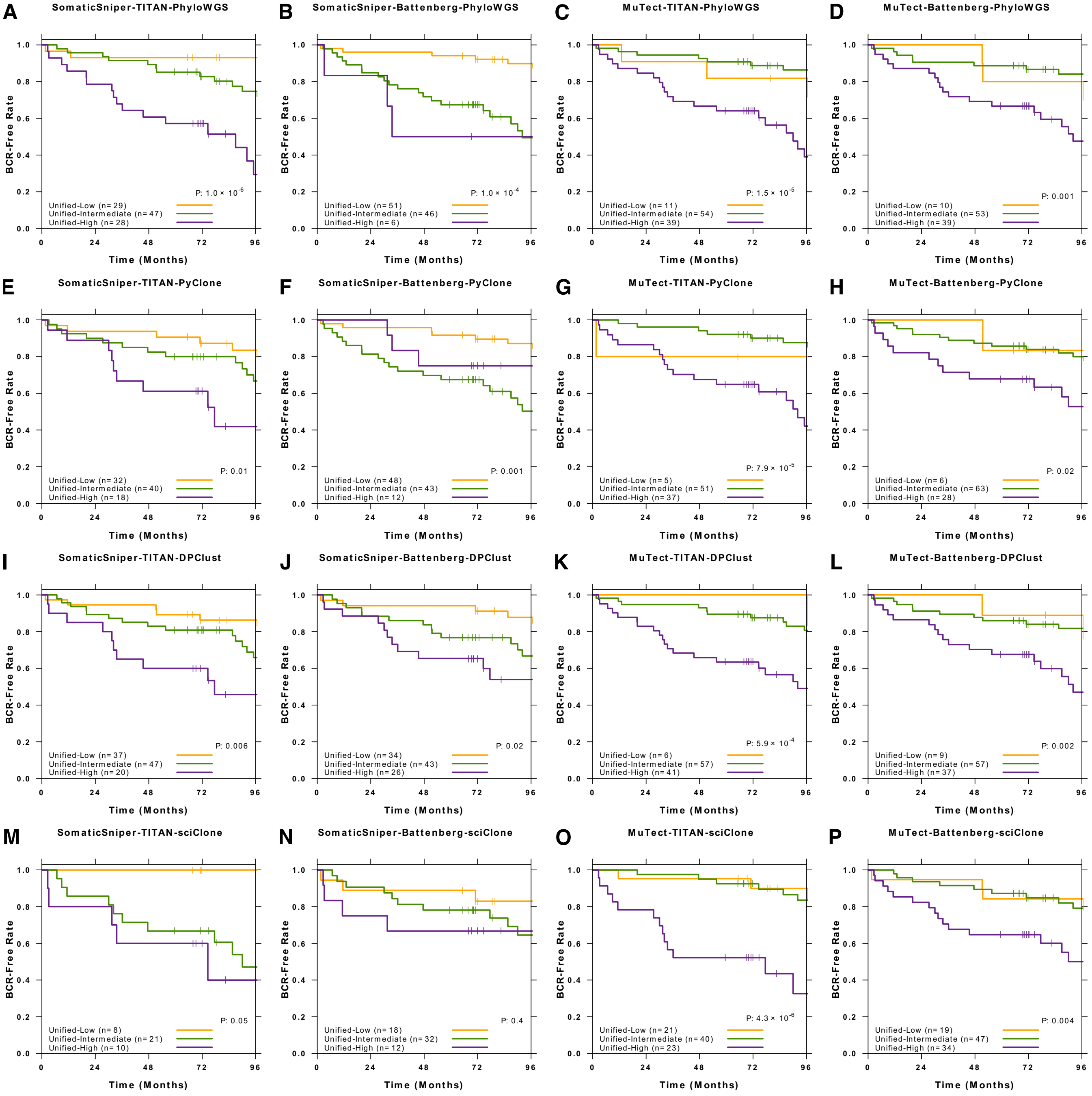


#### Supplementary Figure 11 – Unified Biomarker

Patient recurrence outcomes stratified using a unified biomarker, combining a previously developed multi-modal biomarker with clonality predicted by sixteen pipelines for single-region reconstruction **A-P**). Univariate modelling was carried out using log-rank tests, no covariates were included. Color of line in the KM-plots indicate patient groups. Sample size of each group is given as labeled.

Supplementary Figure 12

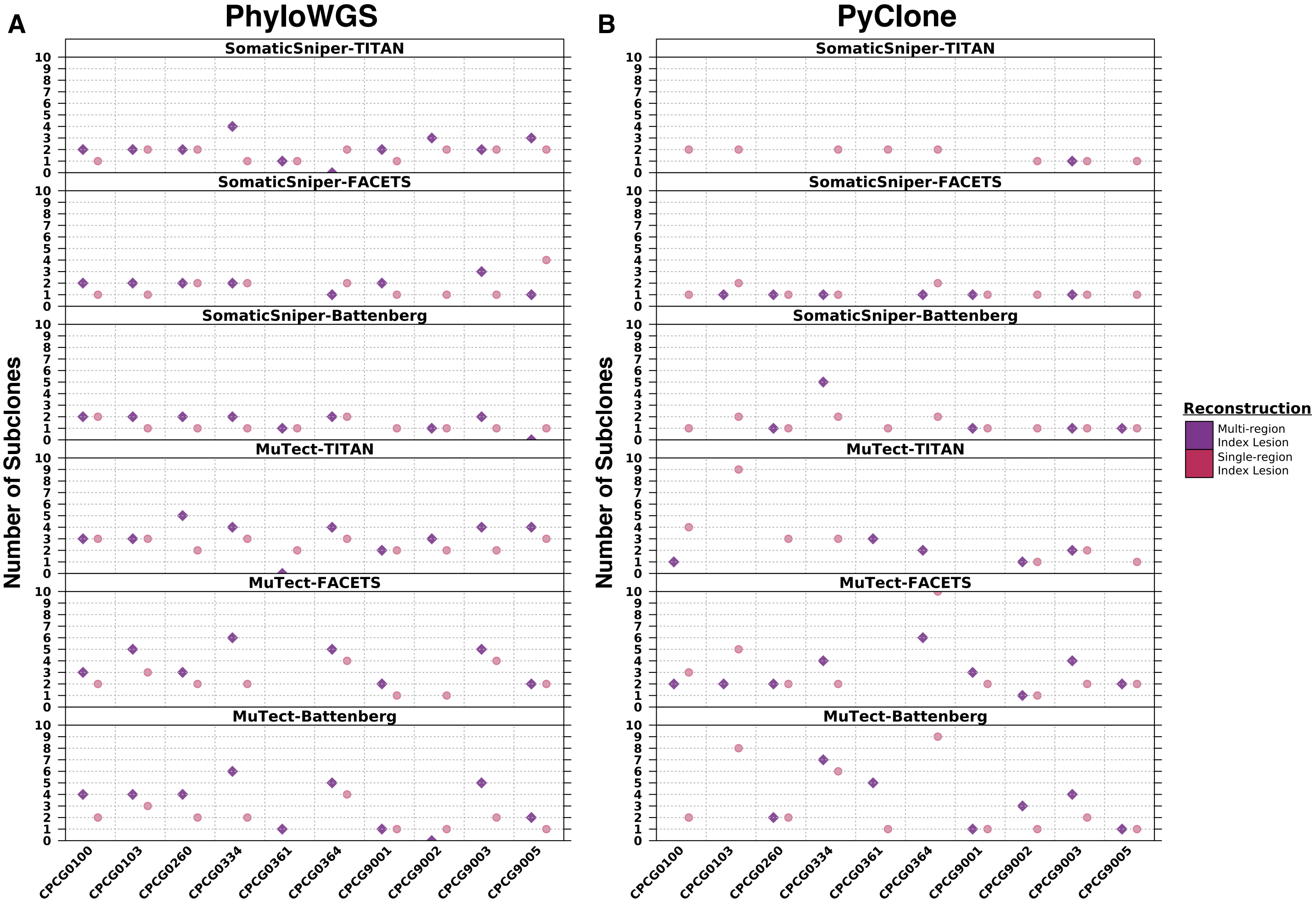


#### Supplementary Figure 12 – Index Lesion in Multi-Region Reconstructions

Number of subclones detected in the index lesion in single-region (pink circle) and multi-region (purple diamond) reconstructions based on pipelines using PhyloWGS **A**) and PyClone **B**). Single-region reconstructions indicate the number of subclones detected in the index lesion using only single-region reconstruction of the index lesion. Multi-region reconstructions indicate the number of subclones detected in the index lesion through multi-region reconstruction of all samples from the tumour. Missing values indicate a failed reconstruction, either single- or multi-region. Multi-region reconstruction SomaticSniper-TITAN-PhyloWGS: n=10 biologically independent samples; SomaticSniper-FACETS-PhyloWGS: n=8; SomaticSniper-Battenberg-PhyloWGS: n=9; MuTect-TITAN-PhyloWGS: n=10; MuTect-FACETS-PhyloWGS: n=8; MuTect-Battenberg-PhyloWGS: n=10; SomaticSniper-TITAN-PyClone: n=1; SomaticSniper-FACETS-PyClone: n=6; SomaticSniper-Battenberg-PyClone: n=5; MuTect-TITAN-PyClone: n=5; MuTect-FACETS-PyClone: n=9; MuTect-Battenberg-PyClone: n=7. Single-region reconstruction SomaticSniper-TITAN-PhyloWGS: n=10; SomaticSniper-FACETS-PhyloWGS: n=9; SomaticSniper-Battenberg-PhyloWGS: n=10; MuTect-TITAN-PhyloWGS: n=10; MuTect-FACETS-PhyloWGS: n=9; MuTect-Battenberg-PhyloWGS: n=9; SomaticSniper-TITAN-PyClone: n=8; SomaticSniper-FACETS-PyClone: n=9; SomaticSniper-Battenberg-PyClone: n=10; MuTect-TITAN-PyClone: n=7; MuTect-FACETS-PyClone: n=9; MuTect-Battenberg-PyClone: n=10.

#### Supplementary Figure 13

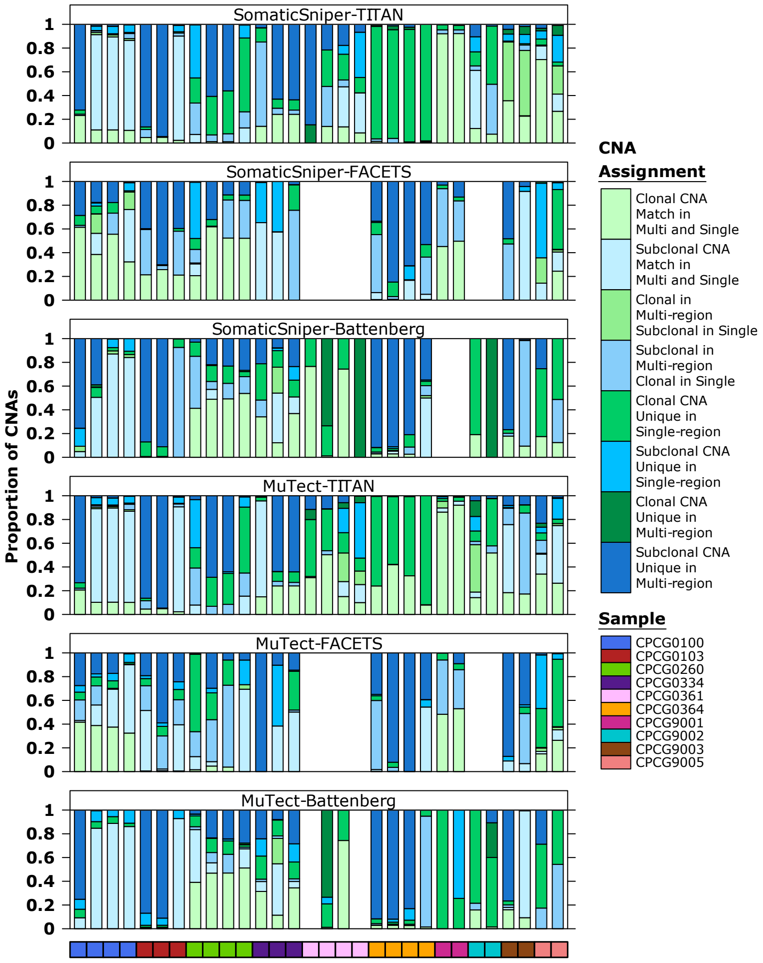


#### Supplementary Figure 13 – Single- and Multi-Region CNA Clonality

Comparison of clonal and subclonal CNA clonality predictions by single- and multi-region subclonal reconstructions in pipelines using PhyloWGS. CNAs were compared by 1.0 Mbp genomic bins between single-region reconstructions and their corresponding multi-region reconstructions. Each stacked bar represents one single-region and the covariate bar color indicates the identity of the sample. Missing bars indicated failed reconstructions, either single- or multi-region. CNAs were grouped into eight categories, delineated by stacked bar color. "Clonal CNA Match in Multi and Single" if the CNA was predicted to be clonal in both single- and multi-region reconstructions and 'Subclonal CNA Match in Multi and Single' if the CNA was predicted to be subclonal in both. CNAs with disagreeing clonality predictions in single- and multi-region reconstructions were grouped as 'Clonal in Multi-region Subclonal in Single' or *vice versa* as 'Subclonal in Multi-region Clonal in Single'. CNAs considered only in single- or multi-region reconstruction and thus did not have a clonality assignment in the reconstruction where it was not considered were given one of the categories: 'Clonal CNA Unique in Single-region', 'Subclonal CNA Unique in Single-region', 'Clonal CNA Unique in Multi-region', or 'Subclonal CNA Unique in Multi-region'. SomaticSniper-TITAN-PhyloWGS: n=30 biologically independent samples; SomaticSniper-FACETS-PhyloWGS: n=24; SomaticSniper-Battenberg-PhyloWGS: n=28; MuTect-TITAN-PhyloWGS: n=30; MuTect-FACETS-PhyloWGS: n=24; MuTect-Battenberg-PhyloWGS: n=28.

#### Supplementary Figure 14

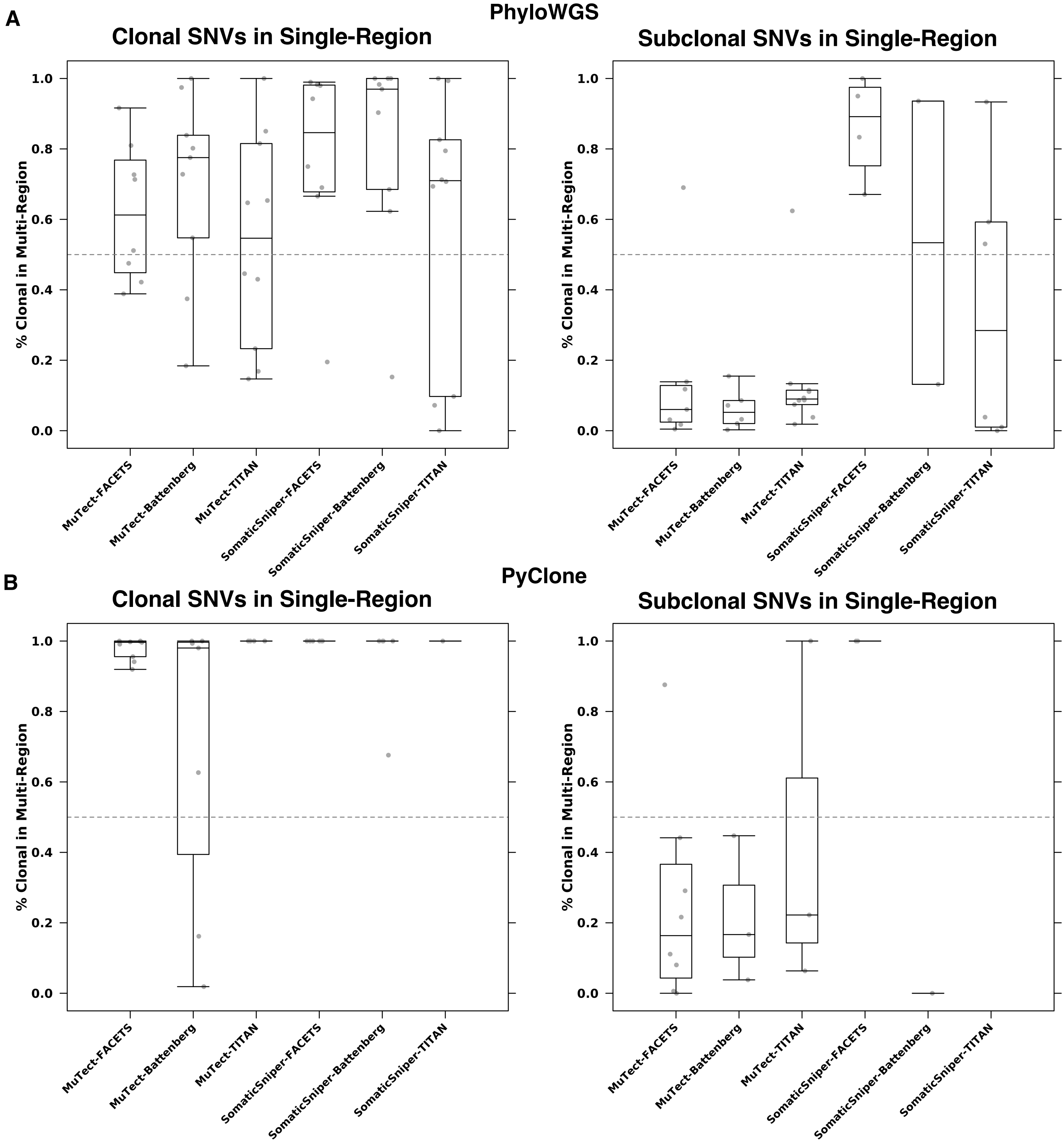


### Supplementary Figure 14 – Index Lesion SNVs in Multi-Region Analyses

Clonal and subclonal mutation cluster composition as defined by SNVs in index lesion single-region reconstructions compared to multi-region reconstructions by pipelines using PhyloWGS **A**) and PyClone **B**). Proportion of SNVs that were predicted to be clonal or subclonal in single-region reconstructions of the index lesion but were predicted to be clonal in multi-region reconstructions. All boxplots show the median (center line, 50$^{th}$ percentile), upper and lower quartiles (box limits, 75$^{th}$ and 25$^{th}$ percentile, respectively), and whiskers extend to the minimum and maximum values within 1.5 times the interquartile range (Tukey boxplots). All data points are represented using grey circles and outliers are those beyond the whiskers. Clonal SNVs in single-region MuTect-FACETS-PhyloWGS: n=8 biologically independent samples; MuTect-Battenberg-PhyloWGS: n=9; MuTect-TITAN-PhyloWGS: n=10; SomaticSniper-FACETS-PhyloWGS: n=8; SomaticSniper-Battenberg-PhyloWGS: n=9; SomaticSniper-TITAN-PhyloWGS: n=10; MuTect-FACETS-PyClone: n=9; MuTect-Battenberg-PyClone: n=7; MuTect-TITAN-PyClone: n=4; SomaticSniper-FACETS-PyClone: n=6; SomaticSniper-Battenberg-PyClone: n=5; SomaticSniper-TITAN-PyClone: n=1. Subclonal SNVs in single-region MuTect-FACETS-PhyloWGS: n=7; MuTect-Battenberg-PhyloWGS: n=6; MuTect-TITAN-PhyloWGS: n=10; SomaticSniper-FACETS-PhyloWGS: n=4; SomaticSniper-Battenberg-PhyloWGS: n=2; SomaticSniper-TITAN-PhyloWGS: n=6; MuTect-FACETS-PyClone: n=8; MuTect-Battenberg-PyClone: n=3; MuTect-TITAN-PyClone: n=3; SomaticSniper-FACETS-PyClone: n=2; SomaticSniper-Battenberg-PyClone: n=1; SomaticSniper-TITAN-PyClone: n=0. Source data are provided as a Source Data file.

### Supplementary Tables

#### Supplementary Table 1 – Failed Reconstructions

Number of reconstructions attempted, number of successful reconstructions and failure rate for all single- and multi-region subclonal reconstruction pipelines. Information is displayed separately for each of the twenty-two different mutation detection tool and subclonal reconstruction algorithm combinations. Summary information is calculated for single-region reconstruction of 293 tumours, single-region reconstruction of 30 samples from 10 tumours with multi-region sequencing, and multi-region reconstructions of the 10 tumours. Each sample that failed reconstruction is listed with the reason of failure.

| | |
|---|---|
| SomaticSniper-TITAN | PhyloWGS |
| Single-Region | Reconstruction |
| Total Samples | 293 |
| Completed Samples | 289 |
| Failure Rate | 0.0136518771331058 |
| Sample | Failure Reason |
| CPCG0166 | Polytumour |
| CPCG0520 | Polytumour |
| CPCG0528 | Polytumour |
| TCGA_7522_T1_wgs | Polytumour |

| | |
|---|---|
| SomaticSniper-TITAN | PhyloWGS |
| Multi-region | Tumours |
| Single-Region | Reconstruction |
| Total Samples | 30 |
| Completed Samples | 30 |
| Failure Rate | 0 |
| Multi-Region | Reconstruction |
| Total Samples | 10 |
| Completed Samples | 10 |
| Failure Rate | 0 |
| Sample | Failure Reason |

| | |
|---|---|
| SomaticSniper-Battenberg | PhyloWGS |
| Single-Region | Reconstruction |
| Total Samples | 293 |
| Completed Samples | 287 |
| Failure Rate | 0.0204778156996587 |
| Sample | Failure Reason |
| CPCG0117 | Failed Battenberg |
| CPCG0124 | Failed Battenberg |
| CPCG0205 | Failed Battenberg |
| CPCG0567 | Below 5 Mutations |
| CPCG0424 | Below 5 Mutations |
| CPCG0074 | Polytumour |

| | |
|---|---|
| SomaticSniper-Battenberg | PhyloWGS |
| Multi-region | Tumours |
| Single-Region | Reconstruction |
| Total Samples | 30 |
| Completed Samples | 30 |
| Failure Rate | 0 |
| Multi-Region | Reconstruction |
| Total Samples | 10 |
| Completed Samples | 9 |
| Failure Rate | 0.1 |
| Sample | Failure Reason |
| CPCG9001-MR | Below 5 Mutations |

| | |
|---|---|
| MuTect-TITAN | PhyloWGS |
| Single-Region | Reconstruction |
| Total Samples | 293 |
| Completed Samples | 289 |
| Failure Rate | 0.0136518771331058 |
| Sample | Failure Reason |
| CPCG0050 | Failed MuTect |
| CPCG0090 | Failed MuTect |
| CPCG0122 | Failed MuTect |
| TCGA_7522_T1_wgs | Polytumour |

| | |
|---|---|
| MuTect-TITAN | PhyloWGS |
| Multi-region | Tumours |
| Single-Region | Reconstruction |
| Total Samples | 30 |
| Completed Samples | 30 |
| Failure Rate | 0 |
| Multi-Region | Reconstruction |
| Total Samples | 10 |
| Completed Samples | 10 |
| Failure Rate | 0 |
| Sample | Failure Reason |

| | |
|---|---|
| MuTect-Battenberg | PhyloWGS |
| Single-Region | Reconstruction |
| Total Samples | 293 |
| Completed Samples | 281 |
| Failure Rate | 0.0409556313993174 |
| Sample | Failure Reason |
| CPCG0117 | Failed Battenberg |
| CPCG0124 | Failed Battenberg |
| CPCG0205 | Failed Battenberg |
| CPCG0050 | Failed MuTect |
| CPCG0090 | Failed MuTect |
| CPCG0122 | Failed MuTect |
| CPCG0089 | Excessive run-time |
| CPCG0587 | Excessive run-time |
| CPCG0575 | Excessive run-time |
| CPCG0361 | Below 10% CP |
| CPCG0575 | Polytumour |
| TCGA_5789_T1_wgs | Polytumour |

| | |
|---|---|
| MuTect-Battenberg | PhyloWGS |
| Multi-region | Tumours |
| Single-Region | Reconstruction |
| Total Samples | 30 |
| Completed Samples | 28 |
| Failure Rate | 0.0666666666666667 |
| Multi-Region | Reconstruction |
| Total Samples | 10 |
| Completed Samples | 10 |
| Failure Rate | 0 |
| Sample | Failure Reason |
| CPCG0361-F1 | Below 10% CP |
| CPCG0361-F4 | Polytumour |

| | |
|---|---|
| SomaticSniper-FACETS | PhyloWGS |
| Multi-region | Tumours |
| Single-Region | Reconstruction |
| Total Samples | 30 |
| Completed Samples | 26 |
| Failure Rate | 0.133333333333333 |
| Multi-Region | Reconstruction |
| Total Samples | 10 |
| Completed Samples | 8 |
| Failure Rate | 0.2 |
| Sample | Failure Reason |
| CPCG9002-MR | Polytumour |
| CPCG0361-F1 | No FACETS |
| CPCG0361-F2 | No FACETS |
| CPCG0361-F3 | No FACETS |
| CPCG0361-F4 | No FACETS |
| CPCG0361-MR | No FACETS |

| | |
|---|---|
| MuTect-FACETS | PhyloWGS |
| Multi-region | Tumours |
| Single-Region | Reconstruction |
| Total Samples | 30 |
| Completed Samples | 26 |
| Failure Rate | 0.133333333333333 |
| Multi-Region | Reconstruction |
| Total Samples | 10 |
| Completed Samples | 8 |
| Failure Rate | 0.2 |
| Sample | Failure Reason |
| CPCG9002-MR | Polytumour |
| CPCG0361-F1 | No FACETS |
| CPCG0361-F2 | No FACETS |
| CPCG0361-F3 | No FACETS |
| CPCG0361-F4 | No FACETS |
| CPCG0361-MR | No FACETS |

| | |
|---|---|
| SomaticSniper-TITAN | PyClone |
| Single-Region | Reconstruction |
| Total Samples | 293 |
| Completed Samples | 221 |
| Failure Rate | 0.245733788395904 |
| Sample | Failure Reason |
| Baca_03-728_T_wgs | No Input |
| Baca_05-2709_T_wgs | No Input |
| Baca_09-3983_T_wgs | No Input |
| Baca_STID0000000410_T_wgs | No Input |
| Berger_1783_T_wgs | No Input |
| Berger_2832_T_wgs | No Input |
| CPCG0015 | No Input |
| CPCG0019 | No Input |
| CPCG0022 | No Input |
| CPCG0027 | No Input |
| CPCG0030 | No Input |
| CPCG0046 | No Input |
| CPCG0072 | No Input |
| CPCG0074 | No Input |
| CPCG0084 | No Input |
| CPCG0089 | No Input |
| CPCG0120 | No Input |
| CPCG0123 | No Input |
| CPCG0154 | No Input |
| CPCG0199 | No Input |
| CPCG0205 | No Input |
| CPCG0212 | No Input |
| CPCG0219 | No Input |
| CPCG0234 | No Input |
| CPCG0237 | No Input |
| CPCG0246 | No Input |
| CPCG0251 | No Input |
| CPCG0259 | No Input |
| CPCG0274 | No Input |
| CPCG0342 | No Input |
| CPCG0344 | No Input |
| CPCG0353 | No Input |
| CPCG0358 | No Input |
| CPCG0373 | No Input |
| CPCG0450 | No Input |
| CPCG0458 | No Input |
| CPCG0525 | No Input |
| CPCG0529 | No Input |
| CPCG0559 | No Input |
| CPCG0562 | No Input |
| CPCG0570 | No Input |
| CPCG0580 | No Input |
| CPCG0584 | No Input |
| CPCG0590 | No Input |
| CPCG0595 | No Input |

| | |
|---|---|
| CPCG0597 | No Input |
| CPCG0600 | No Input |
| TCGA_5503_T1_wgs | No Input |
| TCGA_5506_T1_wgs | No Input |
| TCGA_5763_T1_wgs | No Input |
| TCGA_5771_T1_wgs | No Input |
| TCGA_6336_T1_wgs | No Input |
| TCGA_6370_T1_wgs | No Input |
| TCGA_7079_T1_wgs | No Input |
| TCGA_7233_T1_wgs | No Input |
| TCGA_7522_T1_wgs | No Input |
| TCGA_7737_T1_wgs | No Input |
| Weischenfeldt_02_T_wgs | No Input |
| Weischenfeldt_03_T_wgs | No Input |
| Weischenfeldt_06_T_wgs | No Input |
| Baca_07-5037_T_wgs | No Input |
| Baca_STID0000002621_T_wgs | No Input |
| CPCG0102 | No Input |
| CPCG0452 | No Input |
| CPCG0547 | No Input |
| CPCG0560 | No Input |
| CPCG0589 | No Input |
| Baca_09-396_T_wgs | Below 5 Mutations |
| CPCG0127 | Below 5 Mutations |
| CPCG0360 | Below 5 Mutations |
| CPCG0363 | Below 5 Mutations |
| CPCG0578 | Below 5 Mutations |

| | |
|---|---|
| SomaticSniper-TITAN | PyClone |
| Multi-region | Tumours |
| Single-Region | Reconstruction |
| Total Samples | 30 |
| Completed Samples | 18 |
| Failure Rate | 0.4 |
| Multi-Region | Reconstruction |
| Total Samples | 10 |
| Completed Samples | 1 |
| Failure Rate | 0.9 |
| Sample | Failure Reason |
| CPCG0100-F3 | Below 5 Mutations |
| CPCG0100-F6 | No Input |
| CPCG0103-F4 | No Input |
| CPCG0260-F1 | No Input |
| CPCG0260-F4 | No Input |
| CPCG0361-F3 | No Input |
| CPCG0361-F4 | No Input |
| CPCG0364-F3 | No Input |
| CPCG9001-P1 | No Input |
| CPCG9001-P2 | No Input |
| CPCG9002-P2 | No Input |
| CPCG9005-P2 | Below 5 Mutations |
| CPCG0100-MR | No shared input |

| | |
|---|---|
| CPCG0103-MR | No shared input |
| CPCG0260-MR | No shared input |
| CPCG0334-MR | No shared input |
| CPCG0361-MR | No shared input |
| CPCG0364-MR | No shared input |
| CPCG9001-MR | No shared input |
| CPCG9002-MR | No shared input |
| CPCG9005-MR | No shared input |

| | |
|---|---|
| SomaticSniper-Battenberg | PyClone |
| Single-Region | Reconstruction |
| Total Samples | 293 |
| Completed Samples | 277 |
| Failure Rate | 0.0546075085324232 |
| Sample | Failure Reason |
| CPCG0117 | Failed Battenberg |
| CPCG0124 | Failed Battenberg |
| Baca_STID0000000410_T_wgs | No Input |
| CPCG0074 | No Input |
| CPCG0120 | No Input |
| CPCG0212 | No Input |
| CPCG0219 | No Input |
| CPCG0274 | No Input |
| Weischenfeldt_011_T_wgs | No Input |
| Weischenfeldt_02_T_wgs | No Input |
| Weischenfeldt_09_T_wgs | No Input |
| CPCG0046 | No Input |
| CPCG0089 | No Input |
| Baca_09-396_T_wgs | Below 5 Mutations |
| Berger_1701_T_wgs | Below 5 Mutations |
| Berger_2832_T_wgs | Below 5 Mutations |

| | |
|---|---|
| SomaticSniper-Battenberg | PyClone |
| Multi-region | Tumours |
| Single-Region | Reconstruction |
| Total Samples | 30 |
| Completed Samples | 25 |
| Failure Rate | 0.166666666666667 |
| Multi-Region | Reconstruction |
| Total Samples | 10 |
| Completed Samples | 5 |
| Failure Rate | 0.5 |
| Sample | Failure Reason |
| CPCG0100-F3 | Below 5 Mutations |
| CPCG0100-F5 | No Input |
| CPCG0100-F6 | No Input |
| CPCG0103-F4 | No Input |
| CPCG0364-F4 | No Input |
| CPCG0100-MR | No shared inputs |
| CPCG0103-MR | No shared inputs |
| CPCG0361-MR | Below 5 Mutations |
| CPCG0364-MR | No shared inputs |
| CPCG9002-MR | No shared inputs |

| | |
|---|---|
| MuTect-TITAN | PyClone |
| Single-Region | Reconstruction |
| Total Samples | 293 |
| Completed Samples | 221 |
| Failure Rate | 0.245733788395904 |
| Sample | Failure Reason |
| CPCG0006 | Excessive Run-time |
| CPCG0020 | Excessive Run-time |
| CPCG0040 | Excessive Run-time |
| CPCG0047 | Excessive Run-time |
| CPCG0063 | Excessive Run-time |
| CPCG0166 | Excessive Run-time |
| CPCG0424 | Excessive Run-time |
| CPCG0451 | Excessive Run-time |
| CPCG0574 | Excessive Run-time |
| CPCG0050 | Failed MuTect |
| CPCG0090 | Failed MuTect |
| CPCG0122 | Failed MuTect |
| Baca_05-2709_T_wgs | No Input |
| Baca_STID0000000410_T_wgs | No Input |
| Berger_1783_T_wgs | No Input |
| CPCG0022 | No Input |
| CPCG0027 | No Input |
| CPCG0030 | No Input |
| CPCG0046 | No Input |
| CPCG0084 | No Input |
| CPCG0089 | No Input |
| CPCG0120 | No Input |
| CPCG0123 | No Input |
| CPCG0154 | No Input |
| CPCG0205 | No Input |
| CPCG0219 | No Input |
| CPCG0237 | No Input |
| CPCG0246 | No Input |
| CPCG0251 | No Input |
| CPCG0259 | No Input |
| CPCG0274 | No Input |
| CPCG0342 | No Input |
| CPCG0344 | No Input |
| CPCG0353 | No Input |
| CPCG0358 | No Input |
| CPCG0450 | No Input |
| CPCG0458 | No Input |
| CPCG0529 | No Input |
| CPCG0559 | No Input |
| CPCG0580 | No Input |
| CPCG0590 | No Input |
| CPCG0595 | No Input |
| CPCG0597 | No Input |
| CPCG0600 | No Input |
| TCGA_5503_T1_wgs | No Input |

| | |
|---|---|
| TCGA_5506_T1_wgs | No Input |
| TCGA_5763_T1_wgs | No Input |
| TCGA_5771_T1_wgs | No Input |
| TCGA_6370_T1_wgs | No Input |
| TCGA_7079_T1_wgs | No Input |
| TCGA_7233_T1_wgs | No Input |
| TCGA_7737_T1_wgs | No Input |
| Weischenfeldt_02_T_wgs | No Input |
| Baca_03-728_T_wgs | No Input |
| Baca_STID0000002621_T_wgs | No Input |
| CPCG0019 | No Input |
| CPCG0072 | No Input |
| CPCG0234 | No Input |
| CPCG0525 | No Input |
| CPCG0570 | No Input |
| CPCG0584 | No Input |
| CPCG0589 | No Input |
| TCGA_6336_T1_wgs | No Input |
| TCGA_7522_T1_wgs | No Input |
| Weischenfeldt_03_T_wgs | No Input |
| Weischenfeldt_06_T_wgs | No Input |
| Baca_07-5037_T_wgs | Below 5 Mutations |
| Berger_2832_T_wgs | Below 5 Mutations |
| CPCG0015 | Below 5 Mutations |
| CPCG0199 | Below 5 Mutations |
| CPCG0560 | Below 5 Mutations |
| CPCG0562 | Below 5 Mutations |

| | |
|---|---|
| MuTect-TITAN | PyClone |
| Multi-region | Tumours |
| Single-Region | Reconstruction |
| Total Samples | 30 |
| Completed Samples | 25 |
| Failure Rate | 0.166666666666667 |
| Multi-Region | Reconstruction |
| Total Samples | 10 |
| Completed Samples | 5 |
| Failure Rate | 0.5 |
| Sample | Failure Reason |
| CPCG0103-F4 | No Input |
| CPCG0260-F4 | Below 5 Mutations |
| CPCG0364-F1 | Excessive run-time |
| CPCG9001-P1 | No Input |
| CPCG9001-P2 | No Input |
| CPCG0103-MR | No shared input |
| CPCG0260-MR | No shared input |
| CPCG0334-MR | No shared input |
| CPCG9001-MR | No shared input |
| CPCG9005-MR | No shared input |

| | |
|---|---|
| MuTect-Battenberg | PyClone |
| Single-Region | Reconstruction |
| Total Samples | 293 |
| Completed Samples | 262 |
| Failure Rate | 0.10580204778157 |
| Sample | Failure Reason |
| CPCG0006 | Excessive Run-time |
| CPCG0019 | Excessive Run-time |
| CPCG0020 | Excessive Run-time |
| CPCG0027 | Excessive Run-time |
| CPCG0040 | Excessive Run-time |
| CPCG0047 | Excessive Run-time |
| CPCG0063 | Excessive Run-time |
| CPCG0072 | Excessive Run-time |
| CPCG0084 | Excessive Run-time |
| CPCG0166 | Excessive Run-time |
| CPCG0234 | Excessive Run-time |
| CPCG0249 | Excessive Run-time |
| CPCG0346 | Excessive Run-time |
| CPCG0388 | Excessive Run-time |
| CPCG0424 | Excessive Run-time |
| CPCG0574 | Excessive Run-time |
| CPCG0577 | Excessive Run-time |
| CPCG0589 | Excessive Run-time |
| CPCG0050 | Failed MuTect |
| CPCG0090 | Failed MuTect |
| CPCG0122 | Failed MuTect |
| CPCG0117 | Failed Battenberg |
| CPCG0124 | Failed Battenberg |
| CPCG0046 | No Input |
| CPCG0074 | No Input |
| CPCG0120 | No Input |
| CPCG0219 | No Input |
| CPCG0274 | No Input |
| Weischenfeldt_011_T_wgs | No Input |
| Weischenfeldt_02_T_wgs | No Input |
| Weischenfeldt_09_T_wgs | No Input |

| | |
|---|---|
| MuTect-Battenberg | PyClone |
| Multi-region | Tumours |
| Single-Region | Reconstruction |
| Total Samples | 30 |
| Completed Samples | 28 |
| Failure Rate | 0.0666666666666667 |
| Multi-Region | Reconstruction |
| Total Samples | 10 |
| Completed Samples | 7 |
| Failure Rate | 0.3 |
| Sample | Failure Reason |
| CPCG0100-F5 | No Input |
| CPCG0100-F6 | No Input |

| | |
|---|---|
| CPCG0103-MR | No Input |
| CPCG0100-MR | No shared input |
| CPCG0364-MR | No shared input |

| | |
|---|---|
| SomaticSniper-FACETS | PyClone |
| Multi-region | Tumours |
| Single-Region | Reconstruction |
| Total Samples | 30 |
| Completed Samples | 26 |
| Failure Rate | 0.133333333333333 |
| Multi-Region | Reconstruction |
| Total Samples | 10 |
| Completed Samples | 6 |
| Failure Rate | 0.4 |
| Sample | Failure Reason |
| CPCG0361-F1 | No FACETS |
| CPCG0361-F2 | No FACETS |
| CPCG0361-F3 | No FACETS |
| CPCG0361-F4 | No FACETS |
| CPCG0100-MR | Below 5 Mutations |
| CPCG9002-MR | Below 5 Mutations |
| CPCG9005-MR | Below 5 Mutations |
| CPCG0361-MR | No FACETS |

| | |
|---|---|
| MuTect-FACETS | PyClone |
| Multi-region | Tumours |
| Single-Region | Reconstruction |
| Total Samples | 30 |
| Completed Samples | 25 |
| Failure Rate | 0.166666666666667 |
| Multi-Region | Reconstruction |
| Total Samples | 10 |
| Completed Samples | 9 |
| Failure Rate | 0.1 |
| Sample | Failure Reason |
| CPCG0103-F4 | Excessive Memory Requirementy |
| CPCG0361-F1 | No FACETS |
| CPCG0361-F2 | No FACETS |
| CPCG0361-F3 | No FACETS |
| CPCG0361-F4 | No FACETS |
| CPCG0361-MR | No FACETS |

| | |
|---|---|
| SomaticSniper-TITAN | DPClust |
| Single-Region | Reconstruction |
| Total Samples | 293 |
| Completed Samples | 293 |
| Failure Rate | 0 |
| Sample | Failure Reason |

| | |
|---|---|
| SomaticSniper-Battenberg | DPClust |
| Single-Region | Reconstruction |
| Total Samples | 293 |
| Completed Samples | 291 |
| Failure Rate | 0.0068259385665529 |
| Sample | Failure Reason |
| CPCG0124 | Failed Battenberg |
| CPCG0117 | Failed Battenberg |

| | |
|---|---|
| MuTect-TITAN | DPClust |
| Single-Region | Reconstruction |
| Total Samples | 293 |
| Completed Samples | 283 |
| Failure Rate | 0.0341296928327645 |
| Sample | Failure Reason |
| CPCG0050 | Failed MuTect |
| CPCG0090 | Failed MuTect |
| CPCG0122 | Failed MuTect |
| CPCG0166 | Excessive Memory Requirement |
| CPCG0574 | Excessive Memory Requirement |
| CPCG0063 | Excessive Memory Requirement |
| CPCG0089 | Excessive Memory Requirement |
| CPCG0424 | Excessive Memory Requirement |
| CPCG0084 | Excessive Memory Requirement |
| CPCG0234 | Excessive Memory Requirement |

| | |
|---|---|
| MuTect-Battenberg | DPClust |
| Single-Region | Reconstruction |
| Total Samples | 293 |
| Completed Samples | 288 |
| Failure Rate | 0.0170648464163823 |
| Sample | Failure Reason |
| CPCG0050 | Failed MuTect |
| CPCG0090 | Failed MuTect |
| CPCG0122 | Failed MuTect |
| CPCG0117 | Failed Battenberg |
| CPCG0124 | Failed Battenberg |

| | |
|---|---|
| SomaticSniper-TITAN | sciClone |
| Single-Region | Reconstruction |
| Total Samples | 293 |
| Completed Samples | 100 |
| Failure Rate | 0.658703071672355 |
| Sample | Failure Reason |
| Baca_03-1426_T_wgs | Below 5 Mutations |
| Baca_07-4814_T_wgs | Below 5 Mutations |
| Baca_07-5318_T_wgs | Below 5 Mutations |
| Baca_08-2153_T_wgs | Below 5 Mutations |
| Baca_08-217_T_wgs | Below 5 Mutations |
| Baca_08-5852_T_wgs | Below 5 Mutations |
| Baca_09-146_T_wgs | Below 5 Mutations |
| Baca_09-628_T_wgs | Below 5 Mutations |
| Baca_STID0000003042_T_wgs | Below 5 Mutations |
| Berger_3027_T_wgs | Below 5 Mutations |
| CPCG0003 | Below 5 Mutations |
| CPCG0007 | Below 5 Mutations |
| CPCG0094 | Below 5 Mutations |
| CPCG0095 | Below 5 Mutations |
| CPCG0098 | Below 5 Mutations |
| CPCG0122 | Below 5 Mutations |
| CPCG0185 | Below 5 Mutations |
| CPCG0189 | Below 5 Mutations |
| CPCG0191 | Below 5 Mutations |
| CPCG0194 | Below 5 Mutations |
| CPCG0208 | Below 5 Mutations |
| CPCG0213 | Below 5 Mutations |
| CPCG0232 | Below 5 Mutations |
| CPCG0237 | Below 5 Mutations |
| CPCG0250 | Below 5 Mutations |
| CPCG0260 | Below 5 Mutations |
| CPCG0262 | Below 5 Mutations |
| CPCG0263 | Below 5 Mutations |
| CPCG0265 | Below 5 Mutations |
| CPCG0341 | Below 5 Mutations |
| CPCG0346 | Below 5 Mutations |
| CPCG0356 | Below 5 Mutations |
| CPCG0358 | Below 5 Mutations |
| CPCG0365 | Below 5 Mutations |
| CPCG0366 | Below 5 Mutations |
| CPCG0371 | Below 5 Mutations |
| CPCG0374 | Below 5 Mutations |
| CPCG0378 | Below 5 Mutations |
| CPCG0381 | Below 5 Mutations |
| CPCG0387 | Below 5 Mutations |
| CPCG0391 | Below 5 Mutations |
| CPCG0411 | Below 5 Mutations |
| CPCG0412 | Below 5 Mutations |
| CPCG0451 | Below 5 Mutations |
| CPCG0454 | Below 5 Mutations |

| | |
|---|---|
| CPCG0520 | Below 5 Mutations |
| CPCG0528 | Below 5 Mutations |
| CPCG0551 | Below 5 Mutations |
| CPCG0565 | Below 5 Mutations |
| CPCG0574 | Below 5 Mutations |
| CPCG0575 | Below 5 Mutations |
| CPCG0581 | Below 5 Mutations |
| CPCG0591 | Below 5 Mutations |
| CPCG0592 | Below 5 Mutations |
| TCGA_5789_T1_wgs | Below 5 Mutations |
| TCGA_7744_T1_wgs | Below 5 Mutations |
| Weischenfeldt_03_T_wgs | Below 5 Mutations |
| Baca_02-1431_T_wgs | No Input |
| Baca_03-728_T_wgs | No Input |
| Baca_05-1657_T_wgs | No Input |
| Baca_05-2709_T_wgs | No Input |
| Baca_07-3258_T_wgs | No Input |
| Baca_07-5021_T_wgs | No Input |
| Baca_07-5037_T_wgs | No Input |
| Baca_08-716_T_wgs | No Input |
| Baca_09-37_T_wgs | No Input |
| Baca_09-396_T_wgs | No Input |
| Baca_09-3983_T_wgs | No Input |
| Baca_STID0000000410_T_wgs | No Input |
| Baca_STID0000002525_T_wgs | No Input |
| Baca_STID0000002621_T_wgs | No Input |
| Berger_0508_T_wgs | No Input |
| Berger_0581_T_wgs | No Input |
| Berger_1701_T_wgs | No Input |
| Berger_1783_T_wgs | No Input |
| Berger_2832_T_wgs | No Input |
| Berger_3043_T_wgs | No Input |
| CPCG0001 | No Input |
| CPCG0015 | No Input |
| CPCG0019 | No Input |
| CPCG0022 | No Input |
| CPCG0027 | No Input |
| CPCG0030 | No Input |
| CPCG0046 | No Input |
| CPCG0048 | No Input |
| CPCG0057 | No Input |
| CPCG0072 | No Input |
| CPCG0074 | No Input |
| CPCG0075 | No Input |
| CPCG0078 | No Input |
| CPCG0084 | No Input |
| CPCG0087 | No Input |
| CPCG0089 | No Input |
| CPCG0102 | No Input |
| CPCG0117 | No Input |
| CPCG0120 | No Input |
| CPCG0121 | No Input |

| | |
|---|---|
| CPCG0123 | No Input |
| CPCG0127 | No Input |
| CPCG0154 | No Input |
| CPCG0158 | No Input |
| CPCG0166 | No Input |
| CPCG0182 | No Input |
| CPCG0198 | No Input |
| CPCG0199 | No Input |
| CPCG0205 | No Input |
| CPCG0206 | No Input |
| CPCG0210 | No Input |
| CPCG0212 | No Input |
| CPCG0219 | No Input |
| CPCG0234 | No Input |
| CPCG0236 | No Input |
| CPCG0246 | No Input |
| CPCG0251 | No Input |
| CPCG0255 | No Input |
| CPCG0259 | No Input |
| CPCG0266 | No Input |
| CPCG0267 | No Input |
| CPCG0268 | No Input |
| CPCG0274 | No Input |
| CPCG0339 | No Input |
| CPCG0340 | No Input |
| CPCG0342 | No Input |
| CPCG0344 | No Input |
| CPCG0348 | No Input |
| CPCG0350 | No Input |
| CPCG0352 | No Input |
| CPCG0353 | No Input |
| CPCG0357 | No Input |
| CPCG0360 | No Input |
| CPCG0362 | No Input |
| CPCG0363 | No Input |
| CPCG0368 | No Input |
| CPCG0373 | No Input |
| CPCG0375 | No Input |
| CPCG0380 | No Input |
| CPCG0401 | No Input |
| CPCG0413 | No Input |
| CPCG0414 | No Input |
| CPCG0424 | No Input |
| CPCG0432 | No Input |
| CPCG0433 | No Input |
| CPCG0450 | No Input |
| CPCG0452 | No Input |
| CPCG0458 | No Input |
| CPCG0492 | No Input |
| CPCG0525 | No Input |
| CPCG0529 | No Input |
| CPCG0547 | No Input |

| | |
|---|---|
| CPCG0559 | No Input |
| CPCG0560 | No Input |
| CPCG0562 | No Input |
| CPCG0567 | No Input |
| CPCG0570 | No Input |
| CPCG0577 | No Input |
| CPCG0578 | No Input |
| CPCG0580 | No Input |
| CPCG0584 | No Input |
| CPCG0587 | No Input |
| CPCG0588 | No Input |
| CPCG0589 | No Input |
| CPCG0590 | No Input |
| CPCG0595 | No Input |
| CPCG0596 | No Input |
| CPCG0597 | No Input |
| CPCG0600 | No Input |
| TCGA_5503_T1_wgs | No Input |
| TCGA_5506_T1_wgs | No Input |
| TCGA_5750_T1_wgs | No Input |
| TCGA_5763_T1_wgs | No Input |
| TCGA_5771_T1_wgs | No Input |
| TCGA_6336_T1_wgs | No Input |
| TCGA_6365_T1_wgs | No Input |
| TCGA_6370_T1_wgs | No Input |
| TCGA_7075_T1_wgs | No Input |
| TCGA_7079_T1_wgs | No Input |
| TCGA_7169_T1_wgs | No Input |
| TCGA_7233_T1_wgs | No Input |
| TCGA_7522_T1_wgs | No Input |
| TCGA_7737_T1_wgs | No Input |
| TCGA_7740_T1_wgs | No Input |
| TCGA_7791_T1_wgs | No Input |
| TCGA_8258_T1_wgs | No Input |
| Weischenfeldt_010_T_wgs | No Input |
| Weischenfeldt_011_T_wgs | No Input |
| Weischenfeldt_01_T_wgs | No Input |
| Weischenfeldt_02_T_wgs | No Input |
| Weischenfeldt_04_T_wgs | No Input |
| Weischenfeldt_05_T_wgs | No Input |
| Weischenfeldt_06_T_wgs | No Input |
| Weischenfeldt_07_T_wgs | No Input |
| Weischenfeldt_08_T_wgs | No Input |
| Weischenfeldt_09_T_wgs | No Input |

| | |
|---|---|
| SomaticSniper-TITAN | sciClone |
| Multi-region | Tumours |
| Single-Region | Reconstruction |
| Total Samples | 30 |
| Completed Samples | 3 |
| Failure Rate | 0.9 |
| Multi-Region | Reconstruction |

| | |
|---|---|
| Total Samples | 10 |
| Completed Samples | 0 |
| Failure Rate | 1 |
| Sample | Failure Reason |
| CPCG0260-B1F2 | Below 5 Mutations |
| CPCG0334-B1F3 | Below 5 Mutations |
| CPCG0361-B1F1 | Below 5 Mutations |
| CPCG0361-B1F2 | Below 5 Mutations |
| CPCG0364-B1F2 | Below 5 Mutations |
| CPCG0100-B1F3 | No Input |
| CPCG0100-B1F5 | No Input |
| CPCG0100-B1F6 | No Input |
| CPCG0103-B1F2 | No Input |
| CPCG0103-B1F4 | No Input |
| CPCG0260-B1F1 | No Input |
| CPCG0260-B1F3 | No Input |
| CPCG0260-B1F4 | No Input |
| CPCG0334-B1F1 | No Input |
| CPCG0334-B1F2 | No Input |
| CPCG0361-B1F3 | No Input |
| CPCG0361-B1F4 | No Input |
| CPCG0364-B1F3 | No Input |
| CPCG0364-B1F4 | No Input |
| CPCG9001-B1P1 | No Input |
| CPCG9001-B1P2 | No Input |
| CPCG9002-B1P1 | No Input |
| CPCG9002-B1P2 | No Input |
| CPCG9003-B1P1 | No Input |
| CPCG9003-B1P2 | No Input |
| CPCG9005-B1P1 | No Input |
| CPCG9005-B1P2 | No Input |
| CPCG0100-MR | No Input |
| CPCG0103-MR | No Input |
| CPCG0260-MR | No Input |
| CPCG0334-MR | No Input |
| CPCG0361-MR | No Input |
| CPCG0364-MR | No Input |
| CPCG9001-MR | No Input |
| CPCG9002-MR | No Input |
| CPCG9003-MR | No Input |
| CPCG9005-MR | No Input |

| | |
|---|---|
| SomaticSniper-Battenberg | sciClone |
| Single-Region | Reconstruction |
| Total Samples | 293 |
| Completed Samples | 150 |
| Failure Rate | 0.488054607508532 |
| Sample | Failure Reason |
| CPCG0117 | Failed Battenberg |
| CPCG0124 | Failed Battenberg |
| CPCG0205 | Failed Battenberg |
| Baca_02-1431_T_wgs | Below 5 Mutations |
| Baca_03-1426_T_wgs | Below 5 Mutations |
| Baca_03-728_T_wgs | Below 5 Mutations |
| Baca_04-1243_T_wgs | Below 5 Mutations |
| Baca_05-2709_T_wgs | Below 5 Mutations |
| Baca_07-3258_T_wgs | Below 5 Mutations |
| Baca_07-4814_T_wgs | Below 5 Mutations |
| Baca_08-2153_T_wgs | Below 5 Mutations |
| Baca_08-5852_T_wgs | Below 5 Mutations |
| Baca_08-784_T_wgs | Below 5 Mutations |
| Baca_STID0000003127_T_wgs | Below 5 Mutations |
| Berger_3043_T_wgs | Below 5 Mutations |
| CPCG0006 | Below 5 Mutations |
| CPCG0007 | Below 5 Mutations |
| CPCG0019 | Below 5 Mutations |
| CPCG0027 | Below 5 Mutations |
| CPCG0046 | Below 5 Mutations |
| CPCG0048 | Below 5 Mutations |
| CPCG0057 | Below 5 Mutations |
| CPCG0075 | Below 5 Mutations |
| CPCG0078 | Below 5 Mutations |
| CPCG0084 | Below 5 Mutations |
| CPCG0087 | Below 5 Mutations |
| CPCG0095 | Below 5 Mutations |
| CPCG0098 | Below 5 Mutations |
| CPCG0121 | Below 5 Mutations |
| CPCG0127 | Below 5 Mutations |
| CPCG0154 | Below 5 Mutations |
| CPCG0158 | Below 5 Mutations |
| CPCG0182 | Below 5 Mutations |
| CPCG0189 | Below 5 Mutations |
| CPCG0191 | Below 5 Mutations |
| CPCG0194 | Below 5 Mutations |
| CPCG0198 | Below 5 Mutations |
| CPCG0199 | Below 5 Mutations |
| CPCG0206 | Below 5 Mutations |
| CPCG0213 | Below 5 Mutations |
| CPCG0234 | Below 5 Mutations |
| CPCG0250 | Below 5 Mutations |
| CPCG0255 | Below 5 Mutations |
| CPCG0265 | Below 5 Mutations |
| CPCG0268 | Below 5 Mutations |

| | |
|---|---|
| CPCG0341 | Below 5 Mutations |
| CPCG0348 | Below 5 Mutations |
| CPCG0352 | Below 5 Mutations |
| CPCG0358 | Below 5 Mutations |
| CPCG0363 | Below 5 Mutations |
| CPCG0371 | Below 5 Mutations |
| CPCG0373 | Below 5 Mutations |
| CPCG0374 | Below 5 Mutations |
| CPCG0380 | Below 5 Mutations |
| CPCG0381 | Below 5 Mutations |
| CPCG0387 | Below 5 Mutations |
| CPCG0391 | Below 5 Mutations |
| CPCG0401 | Below 5 Mutations |
| CPCG0411 | Below 5 Mutations |
| CPCG0412 | Below 5 Mutations |
| CPCG0528 | Below 5 Mutations |
| CPCG0529 | Below 5 Mutations |
| CPCG0557 | Below 5 Mutations |
| CPCG0561 | Below 5 Mutations |
| CPCG0565 | Below 5 Mutations |
| CPCG0575 | Below 5 Mutations |
| CPCG0580 | Below 5 Mutations |
| CPCG0581 | Below 5 Mutations |
| CPCG0590 | Below 5 Mutations |
| CPCG0591 | Below 5 Mutations |
| CPCG0592 | Below 5 Mutations |
| CPCG0595 | Below 5 Mutations |
| TCGA_5506_T1_wgs | Below 5 Mutations |
| TCGA_5750_T1_wgs | Below 5 Mutations |
| TCGA_5763_T1_wgs | Below 5 Mutations |
| TCGA_5771_T1_wgs | Below 5 Mutations |
| TCGA_5789_T1_wgs | Below 5 Mutations |
| TCGA_6336_T1_wgs | Below 5 Mutations |
| TCGA_6365_T1_wgs | Below 5 Mutations |
| TCGA_6370_T1_wgs | Below 5 Mutations |
| TCGA_7075_T1_wgs | Below 5 Mutations |
| TCGA_7079_T1_wgs | Below 5 Mutations |
| TCGA_7169_T1_wgs | Below 5 Mutations |
| TCGA_7233_T1_wgs | Below 5 Mutations |
| TCGA_7522_T1_wgs | Below 5 Mutations |
| TCGA_7737_T1_wgs | Below 5 Mutations |
| TCGA_8258_T1_wgs | Below 5 Mutations |
| Weischenfeldt_01_T_wgs | Below 5 Mutations |
| Weischenfeldt_02_T_wgs | Below 5 Mutations |
| Weischenfeldt_03_T_wgs | Below 5 Mutations |
| Weischenfeldt_04_T_wgs | Below 5 Mutations |
| Weischenfeldt_05_T_wgs | Below 5 Mutations |
| Weischenfeldt_06_T_wgs | Below 5 Mutations |
| Weischenfeldt_08_T_wgs | Below 5 Mutations |
| Weischenfeldt_09_T_wgs | Below 5 Mutations |
| Baca_01-28_T_wgs | No Input |
| Baca_09-37_T_wgs | No Input |

| | |
|---|---|
| Baca_09-396_T_wgs | No Input |
| Baca_STID0000000410_T_wgs | No Input |
| Baca_STID0000003042_T_wgs | No Input |
| Berger_0508_T_wgs | No Input |
| Berger_0581_T_wgs | No Input |
| Berger_1701_T_wgs | No Input |
| Berger_1783_T_wgs | No Input |
| Berger_2832_T_wgs | No Input |
| Berger_3027_T_wgs | No Input |
| CPCG0074 | No Input |
| CPCG0083 | No Input |
| CPCG0089 | No Input |
| CPCG0120 | No Input |
| CPCG0166 | No Input |
| CPCG0185 | No Input |
| CPCG0196 | No Input |
| CPCG0208 | No Input |
| CPCG0212 | No Input |
| CPCG0219 | No Input |
| CPCG0274 | No Input |
| CPCG0344 | No Input |
| CPCG0353 | No Input |
| CPCG0368 | No Input |
| CPCG0375 | No Input |
| CPCG0413 | No Input |
| CPCG0414 | No Input |
| CPCG0424 | No Input |
| CPCG0433 | No Input |
| CPCG0437 | No Input |
| CPCG0450 | No Input |
| CPCG0454 | No Input |
| CPCG0492 | No Input |
| CPCG0525 | No Input |
| CPCG0550 | No Input |
| CPCG0562 | No Input |
| CPCG0567 | No Input |
| CPCG0574 | No Input |
| CPCG0578 | No Input |
| CPCG0584 | No Input |
| CPCG0587 | No Input |
| TCGA_5503_T1_wgs | No Input |
| TCGA_7740_T1_wgs | No Input |
| TCGA_7791_T1_wgs | No Input |
| Weischenfeldt_010_T_wgs | No Input |
| Weischenfeldt_011_T_wgs | No Input |
| Weischenfeldt_07_T_wgs | No Input |

| | |
|---|---|
| SomaticSniper-Battenberg | sciClone |
| Multi-region | Tumours |
| Single-Region | Reconstruction |
| Total Samples | 30 |
| Completed Samples | 5 |

| | |
|---|---|
| Failure Rate | 0.833333333333333 |
| Multi-Region | Reconstruction |
| Total Samples | 10 |
| Completed Samples | 0 |
| Failure Rate | 1 |
| Sample | Failure Reason |
| CPCG0260-B1F2 | Below 5 Mutations |
| CPCG0260-B1F3 | Below 5 Mutations |
| CPCG0260-B1F4 | Below 5 Mutations |
| CPCG0334-B1F3 | Below 5 Mutations |
| CPCG0361-B1F2 | Below 5 Mutations |
| CPCG0364-B1F2 | Below 5 Mutations |
| CPCG0364-B1F3 | Below 5 Mutations |
| CPCG9002-B1P2 | Below 5 Mutations |
| CPCG9003-B1P1 | Below 5 Mutations |
| CPCG9003-B1P2 | Below 5 Mutations |
| CPCG9005-B1P1 | Below 5 Mutations |
| CPCG9005-B1P2 | Below 5 Mutations |
| CPCG0100-B1F3 | No Input |
| CPCG0100-B1F5 | No Input |
| CPCG0100-B1F6 | No Input |
| CPCG0103-B1F2 | No Input |
| CPCG0103-B1F4 | No Input |
| CPCG0334-B1F2 | No Input |
| CPCG0361-B1F1 | No Input |
| CPCG0361-B1F3 | No Input |
| CPCG0361-B1F4 | No Input |
| CPCG0364-B1F4 | No Input |
| CPCG9001-B1P1 | No Input |
| CPCG9001-B1P2 | No Input |
| CPCG9002-B1P1 | No Input |
| CPCG0100-MR | No Input |
| CPCG0103-MR | No Input |
| CPCG0260-MR | No Input |
| CPCG0334-MR | No Input |
| CPCG0361-MR | No Input |
| CPCG0364-MR | No Input |
| CPCG9001-MR | No Input |
| CPCG9002-MR | No Input |
| CPCG9003-MR | No Input |
| CPCG9005-MR | No Input |

| | |
|---|---|
| MuTect-TITAN | sciClone |
| Single-Region | Reconstruction |
| Total Samples | 293 |
| Completed Samples | 182 |
| Failure Rate | 0.378839590443686 |
| Sample | Failure Reason |
| CPCG0050 | Failed MuTect |
| CPCG0090 | Failed MuTect |
| CPCG0122 | Failed MuTect |
| Baca_02-1431_T_wgs | Below 5 Mutations |
| Baca_03-728_T_wgs | Below 5 Mutations |
| Baca_07-5037_T_wgs | Below 5 Mutations |
| Baca_08-217_T_wgs | Below 5 Mutations |
| Baca_08-784_T_wgs | Below 5 Mutations |
| Baca_09-37_T_wgs | Below 5 Mutations |
| Baca_09-396_T_wgs | Below 5 Mutations |
| Baca_STID0000000410_T_wgs | Below 5 Mutations |
| Baca_STID0000002525_T_wgs | Below 5 Mutations |
| Baca_STID0000002621_T_wgs | Below 5 Mutations |
| Berger_1701_T_wgs | Below 5 Mutations |
| CPCG0003 | Below 5 Mutations |
| CPCG0006 | Below 5 Mutations |
| CPCG0007 | Below 5 Mutations |
| CPCG0015 | Below 5 Mutations |
| CPCG0019 | Below 5 Mutations |
| CPCG0022 | Below 5 Mutations |
| CPCG0027 | Below 5 Mutations |
| CPCG0030 | Below 5 Mutations |
| CPCG0046 | Below 5 Mutations |
| CPCG0072 | Below 5 Mutations |
| CPCG0087 | Below 5 Mutations |
| CPCG0102 | Below 5 Mutations |
| CPCG0123 | Below 5 Mutations |
| CPCG0154 | Below 5 Mutations |
| CPCG0182 | Below 5 Mutations |
| CPCG0199 | Below 5 Mutations |
| CPCG0205 | Below 5 Mutations |
| CPCG0206 | Below 5 Mutations |
| CPCG0246 | Below 5 Mutations |
| CPCG0251 | Below 5 Mutations |
| CPCG0260 | Below 5 Mutations |
| CPCG0268 | Below 5 Mutations |
| CPCG0340 | Below 5 Mutations |
| CPCG0342 | Below 5 Mutations |
| CPCG0352 | Below 5 Mutations |
| CPCG0353 | Below 5 Mutations |
| CPCG0358 | Below 5 Mutations |
| CPCG0362 | Below 5 Mutations |
| CPCG0373 | Below 5 Mutations |
| CPCG0413 | Below 5 Mutations |
| CPCG0433 | Below 5 Mutations |

| | |
|---|---|
| CPCG0450 | Below 5 Mutations |
| CPCG0492 | Below 5 Mutations |
| CPCG0565 | Below 5 Mutations |
| CPCG0570 | Below 5 Mutations |
| CPCG0577 | Below 5 Mutations |
| CPCG0578 | Below 5 Mutations |
| CPCG0588 | Below 5 Mutations |
| CPCG0589 | Below 5 Mutations |
| CPCG0592 | Below 5 Mutations |
| CPCG0597 | Below 5 Mutations |
| CPCG0600 | Below 5 Mutations |
| TCGA_5750_T1_wgs | Below 5 Mutations |
| TCGA_5771_T1_wgs | Below 5 Mutations |
| TCGA_6365_T1_wgs | Below 5 Mutations |
| TCGA_7169_T1_wgs | Below 5 Mutations |
| TCGA_7522_T1_wgs | Below 5 Mutations |
| TCGA_8258_T1_wgs | Below 5 Mutations |
| Weischenfeldt_010_T_wgs | Below 5 Mutations |
| Baca_05-2709_T_wgs | No Input |
| Baca_07-3258_T_wgs | No Input |
| Baca_07-5318_T_wgs | No Input |
| Baca_09-3983_T_wgs | No Input |
| Berger_0581_T_wgs | No Input |
| Berger_1783_T_wgs | No Input |
| Berger_2832_T_wgs | No Input |
| Berger_3043_T_wgs | No Input |
| CPCG0001 | No Input |
| CPCG0057 | No Input |
| CPCG0074 | No Input |
| CPCG0078 | No Input |
| CPCG0089 | No Input |
| CPCG0117 | No Input |
| CPCG0120 | No Input |
| CPCG0198 | No Input |
| CPCG0212 | No Input |
| CPCG0219 | No Input |
| CPCG0274 | No Input |
| CPCG0432 | No Input |
| CPCG0458 | No Input |
| CPCG0525 | No Input |
| CPCG0529 | No Input |
| CPCG0559 | No Input |
| CPCG0560 | No Input |
| CPCG0562 | No Input |
| CPCG0580 | No Input |
| CPCG0584 | No Input |
| CPCG0590 | No Input |
| CPCG0595 | No Input |
| CPCG0596 | No Input |
| TCGA_5503_T1_wgs | No Input |
| TCGA_5506_T1_wgs | No Input |
| TCGA_5763_T1_wgs | No Input |

| | |
|---|---|
| TCGA_6336_T1_wgs | No Input |
| TCGA_6370_T1_wgs | No Input |
| TCGA_7079_T1_wgs | No Input |
| TCGA_7233_T1_wgs | No Input |
| TCGA_7737_T1_wgs | No Input |
| Weischenfeldt_011_T_wgs | No Input |
| Weischenfeldt_01_T_wgs | No Input |
| Weischenfeldt_02_T_wgs | No Input |
| Weischenfeldt_04_T_wgs | No Input |
| Weischenfeldt_05_T_wgs | No Input |
| Weischenfeldt_06_T_wgs | No Input |
| Weischenfeldt_07_T_wgs | No Input |
| Weischenfeldt_08_T_wgs | No Input |
| Weischenfeldt_09_T_wgs | No Input |

| | |
|---|---|
| MuTect-TITAN | sciClone |
| Multi-region | Tumours |
| Single-Region | Reconstruction |
| Total Samples | 30 |
| Completed Samples | 11 |
| Failure Rate | 0.633333333333333 |
| Multi-Region | Reconstruction |
| Total Samples | 10 |
| Completed Samples | 0 |
| Failure Rate | 1 |
| Sample | Failure Reason |
| CPCG0260-B1F1 | Below 5 Mutations |
| CPCG0260-B1F4 | Below 5 Mutations |
| CPCG0334-B1F1 | Below 5 Mutations |
| CPCG0361-B1F4 | Below 5 Mutations |
| CPCG0364-B1F3 | Below 5 Mutations |
| CPCG9002-B1P1 | Below 5 Mutations |
| CPCG9003-B1P1 | Below 5 Mutations |
| CPCG9003-B1P2 | Below 5 Mutations |
| CPCG9005-B1P1 | Below 5 Mutations |
| CPCG0100-B1F3 | No Input |
| CPCG0100-B1F5 | No Input |
| CPCG0100-B1F6 | No Input |
| CPCG0103-B1F4 | No Input |
| CPCG0361-B1F3 | No Input |
| CPCG0364-B1F4 | No Input |
| CPCG9001-B1P1 | No Input |
| CPCG9001-B1P2 | No Input |
| CPCG9002-B1P2 | No Input |
| CPCG9005-B1P2 | No Input |
| CPCG0100-MR | No Input |
| CPCG0103-MR | No Input |
| CPCG0260-MR | No Input |
| CPCG0334-MR | No Input |
| CPCG0361-MR | No Input |
| CPCG0364-MR | No Input |
| CPCG9001-MR | No Input |

| | |
|---|---|
| CPCG9002-MR | No Input |
| CPCG9003-MR | No Input |
| CPCG9005-MR | No Input |

| | |
|---|---|
| MuTect-Battenberg | sciClone |
| Single-Region | Reconstruction |
| Total Samples | 293 |
| Completed Samples | 257 |
| Failure Rate | 0.122866894197952 |
| Sample | Failure Reason |
| CPCG0050 | Failed MuTect |
| CPCG0090 | Failed MuTect |
| CPCG0122 | Failed MuTect |
| CPCG0117 | Failed Battenberg |
| CPCG0124 | Failed Battenberg |
| CPCG0205 | Failed Battenberg |
| Baca_05-2709_T_wgs | Below 5 Mutations |
| Baca_08-784_T_wgs | Below 5 Mutations |
| Baca_09-37_T_wgs | Below 5 Mutations |
| Baca_STID0000003042_T_wgs | Below 5 Mutations |
| Berger_0508_T_wgs | Below 5 Mutations |
| Berger_0581_T_wgs | Below 5 Mutations |
| Berger_1783_T_wgs | Below 5 Mutations |
| Berger_3027_T_wgs | Below 5 Mutations |
| Berger_3043_T_wgs | Below 5 Mutations |
| CPCG0006 | Below 5 Mutations |
| CPCG0074 | Below 5 Mutations |
| CPCG0083 | Below 5 Mutations |
| CPCG0189 | Below 5 Mutations |
| CPCG0212 | Below 5 Mutations |
| CPCG0413 | Below 5 Mutations |
| CPCG0437 | Below 5 Mutations |
| CPCG0450 | Below 5 Mutations |
| CPCG0492 | Below 5 Mutations |
| CPCG0550 | Below 5 Mutations |
| Weischenfeldt_010_T_wgs | Below 5 Mutations |
| Weischenfeldt_011_T_wgs | Below 5 Mutations |
| Weischenfeldt_05_T_wgs | Below 5 Mutations |
| Weischenfeldt_06_T_wgs | Below 5 Mutations |
| Berger_1701_T_wgs | No Input |
| Berger_2832_T_wgs | No Input |
| Weischenfeldt_01_T_wgs | No Input |
| Weischenfeldt_02_T_wgs | No Input |
| Weischenfeldt_04_T_wgs | No Input |
| Weischenfeldt_07_T_wgs | No Input |
| Weischenfeldt_09_T_wgs | No Input |

| | |
|---|---|
| MuTect-Battenberg | sciClone |
| Multi-region | Tumours |
| Single-Region | Reconstruction |
| Total Samples | 30 |
| Completed Samples | 20 |
| Failure Rate | 0.333333333333333 |
| Multi-Region | Reconstruction |
| Total Samples | 10 |

| | |
|---|---|
| Completed Samples | 2 |
| Failure Rate | 0.8 |
| Sample | Failure Reason |
| CPCG0100-B1F5 | Below 5 Mutations |
| CPCG0103-B1F4 | Below 5 Mutations |
| CPCG0364-B1F4 | Below 5 Mutations |
| CPCG9001-B1P1 | Below 5 Mutations |
| CPCG9001-B1P2 | Below 5 Mutations |
| CPCG9002-B1P1 | Below 5 Mutations |
| CPCG9005-B1P1 | Below 5 Mutations |
| CPCG0100-B1F3 | No Input |
| CPCG0100-B1F6 | No Input |
| CPCG9005-B1P2 | No Input |
| CPCG0100-MR | No Input |
| CPCG0103-MR | No Input |
| CPCG0260-MR | Below 5 Mutations |
| CPCG0364-MR | No Input |
| CPCG9001-MR | No Input |
| CPCG9002-MR | No Input |
| CPCG9003-MR | Below 5 Mutations |
| CPCG9005-MR | No Input |

| | |
|---|---|
| SomaticSniper-FACETS | sciClone |
| Multi-region | Tumours |
| Single-Region | Reconstruction |
| Total Samples | 30 |
| Completed Samples | 5 |
| Failure Rate | 0.833333333333333 |
| Multi-Region | Reconstruction |
| Total Samples | 10 |
| Completed Samples | 0 |
| Failure Rate | 1 |
| Sample | Failure Reason |
| CPCG0361-F1 | No FACETS |
| CPCG0361-F2 | No FACETS |
| CPCG0361-F3 | No FACETS |
| CPCG0361-F4 | No FACETS |
| CPCG0361-MR | No FACETS |
| CPCG0260-B1F3 | Below 5 Mutations |
| CPCG0260-B1F4 | Below 5 Mutations |
| CPCG0334-B1F3 | Below 5 Mutations |
| CPCG0364-B1F2 | Below 5 Mutations |
| CPCG0364-B1F3 | Below 5 Mutations |
| CPCG9003-B1P1 | Below 5 Mutations |
| CPCG9005-B1P1 | Below 5 Mutations |
| CPCG9005-B1P2 | Below 5 Mutations |
| CPCG0100-B1F3 | No Input |
| CPCG0100-B1F5 | No Input |
| CPCG0100-B1F6 | No Input |
| CPCG0100-MR | No Input |
| CPCG0103-B1F2 | No Input |
| CPCG0103-B1F4 | No Input |
| CPCG0103-MR | No Input |
| CPCG0260-B1F2 | No Input |
| CPCG0260-MR | No Input |
| CPCG0334-B1F2 | No Input |
| CPCG0334-MR | No Input |
| CPCG0364-B1F4 | No Input |
| CPCG0364-MR | No Input |
| CPCG9001-B1P1 | No Input |
| CPCG9001-B1P2 | No Input |
| CPCG9001-MR | No Input |
| CPCG9002-B1P1 | No Input |
| CPCG9002-B1P2 | No Input |
| CPCG9002-MR | No Input |
| CPCG9003-B1P2 | No Input |
| CPCG9003-MR | No Input |
| CPCG9005-MR | No Input |

| | |
|---|---|
| MuTect-FACETS | sciClone |
| Multi-region | Tumours |
| Single-Region | Reconstruction |
| Total Samples | 30 |
| Completed Samples | 20 |
| Failure Rate | 0.333333333333333 |
| Multi-Region | Reconstruction |
| Total Samples | 10 |
| Completed Samples | 2 |
| Failure Rate | 0.8 |
| Sample | Failure Reason |
| CPCG0361-F1 | No FACETS |
| CPCG0361-F2 | No FACETS |
| CPCG0361-F3 | No FACETS |
| CPCG0361-F4 | No FACETS |
| CPCG0361-MR | No FACETS |
| CPCG0100-B1F6 | Below 5 Mutations |
| CPCG0260-MR | Below 5 Mutations |
| CPCG9002-B1P1 | Below 5 Mutations |
| CPCG9005-B1P1 | Below 5 Mutations |
| CPCG9005-B1P2 | Below 5 Mutations |
| CPCG0100-B1F3 | No Input |
| CPCG0100-MR | No Input |
| CPCG0364-B1F4 | No Input |
| CPCG0364-MR | No Input |
| CPCG0103-MR | No shared input |
| CPCG9001-MR | No shared input |
| CPCG9005-MR | No Input |
| CPCG9002-MR | No Input |
